# Seal milk oligosaccharides rival human milk complexity and exhibit functional dynamics during lactation

**DOI:** 10.1101/2025.03.20.644374

**Authors:** Chunsheng Jin, Jon Lundstrøm, Carmen R. Cori, Shih-Yun Guu, Alexander R. Bennett, Mirjam Dannborg, Johan Bengtsson-Palme, Rachel Hevey, Kay-Hooi Khoo, Daniel Bojar

**Affiliations:** Proteomics Core Facility at Sahlgrenska Academy, University of Gothenburg, Gothenburg, 405 30; Sweden; Department of Chemistry and Molecular Biology; University of Gothenburg; Gothenburg, 405 30; Sweden. Wallenberg Centre for Molecular and Translational Medicine; University of Gothenburg; Gothenburg, 405 30; Sweden; Dept. Pharmaceutical Sciences, University of Basel, Klingelbergstr. 50, 4056, Basel, Switzerland; Institute of Biological Chemistry, Academia Sinica, Taipei 11529, Taiwan; Department of Medical Biochemistry, Institute of Biomedicine, University of Gothenburg, Gothenburg, 405 30; Sweden; Division of Systems and Synthetic Biology, Department of Life Sciences, SciLifeLab, Chalmers University of Technology, 412 96, Gothenburg, Sweden. Department of Infectious Diseases, Institute of Biomedicine, The Sahlgrenska Academy, University of Gothenburg, Guldhedsgatan 10A, 413 46, Gothenburg, Sweden. Centre for Antibiotic Resistance Research (CARe), Gothenburg, Sweden

## Abstract

Breast milk oligosaccharides are crucial for neonatal development and health. Yet most milk research focuses on humans, or domesticated mammals that are historically poor in milk oligosaccharide complexity. Here, we perform an exhaustive mass spectrometry-driven structural characterization of milk oligosaccharides in a wild mammal, Atlantic grey seals (*Halichoerus grypus*), throughout their lactation period. Characterizing and quantifying 332 milk oligosaccharides, including 166 novel structures, we reveal seals to rival human milk in complexity, with seal free oligosaccharides reaching unprecedented 28 monosaccharides in size. Glycomics and metabolomics time course analysis establishes a concerted regulatory process reshaping the seal milk glycome throughout lactation,similar as in human milk. Functional analysis of herein newly characterized structures reveals anti-biofilm effects and immunomodulatory functions of seal milk oligosaccharides. We envision these findings to overturn long-held assumptions about milk complexity of non-human mammals and enable insights into the functional relevance of complex carbohydrates in breast milk.

## Introduction

Milk oligosaccharides (MOs), soluble glycans resulting from the elaboration of lactose in mammalian breast milk, are key contributors to infant development and health (*1*). The exact sequences of MOs in the resulting milk glycome can have measurable impacts on protecting neonates from pathogens, via competitive inhibition or nurturing the initial microbiome by selecting for MO degraders such as *Bifidobacteria* (*2*). Critically, structural diversity in MOs— including features such as fucosylation, sialylation, and sulfation—directly determines their functional specificity, enabling tailored interactions with pathogens, immune cells, and commensal microbes across species. An evolutionary tuning of the milk glycome across species (*3–5*) suggests that understudied mammals, particularly those with distinct ecological pressures, may harbor unique MO repertoires with novel bioactivities.

The current best estimate for the pan-mammalian milk glycome, accounting for structural ambiguities, is >650 known structures (*4*), of which ∼100 have been recently newly characterized by our group (*3*). A large portion of the remaining structures have been discovered in humans, due to an incredible research focus on human breast milk over the last century. Currently, around 300 unique MO structures have been characterized in human milk (*4*), with the largest human MO reaching 18 monosaccharide building blocks (*6*), but it is important to note that any individual human milk sample contains far fewer identifiable structures.

One reason for our lacking knowledge in non-human MOs is that sample access can be a bottleneck, especially given that domesticated mammals (e.g., cows, goats, sheep), which would be readily accessible, have much lower oligosaccharide levels than their wild counterparts (*5, 7, 8*), further hampering MO discovery. We have noted previously that reported milk glycome diversities are essentially a function of the number of published studies on a given species (*4*) and expect the true biochemical diversity of non-human milk to be far larger than currently assumed.

Marine mammals, in particular, represent an underexplored frontier: their unique evolutionary trajectories and extreme environments (e.g., high pathogen exposure, rapid postnatal development) likely drive the evolution of specialized MOs with potent protective or developmental roles. Further, they present a sampling challenge, contributing to the lack of studies in this direction. Our previous findings on dolphin MOs (*3*) support this hypothesis.

Seals are an especially promising MO reservoir because they do not rely on carbohydrates as an energy source in their milk (*9*), leading to a high ratio of oligosaccharides to lactose, which in turn facilitates the characterization of many unique structures. Thus, MOs of seal species such as hooded seal (*Cystophora cristata*) (*10*), Australian fur seal (*Arctocephalus pusillus doriferus*) (*10*), Arctic harbor seal (*Phoca vitulina vitulina*) (*11*), and bearded seal (*Erignathus barbatus*) (*12*) have been previously characterized, typically identifying fucosylated structures, which are often used as decoy receptors (*13*). Yet even in this group of model species for rich milk glycomes, we lack a thorough characterization of (i) the full structural MO complexity and (ii) their change over time during lactation.

We here for the first time present the full milk glycome of the Atlantic grey seal (*Halichoerus grypus*), with hundreds of free milk oligosaccharides in a longitudinal dataset following the lactation period. Previously, only six short milk oligosaccharides from this species had been reported in metabolomics data (*14, 15*). Next to a great degree of structural diversity and novelty (increasing the number of all known MOs by >20%), as well as the hitherto largest characterized MOs, we unveil concerted changes in the glycome profile throughout lactation via time course analysis, as well as a multi-omics analysis with paired metabolomics data. We then engaged in functional experiments to show that some of the structures that are (i) novel and (ii) changing during lactation have potent anti-biofilm and immunomodulatory functions. We envision that the extraordinary richness in milk oligosaccharides of the milk of *H. grypus* can serve as a model system to improve our understanding of lactation and the health impact of the milk glycome.

## Results

### Seal milk harbors substantial glycan sequence diversity

To map the species diversity of milk oligosaccharides (MOs) in *H. grypus* (grey seal), we analyzed samples from five individuals at multiple timepoints during the lactation period (days 2, 7, 13, and 17/18/19 after birth), for a total of 20 biological samples. It is important to note that the lactation period differs dramatically in seals (*16*), with the four days of *Cystophora cristata* being the shortest of any mammal. *H. grypus* lactates 17 days on average, making this a complete dataset of the entire lactation period.

We measured all our samples via neutral/acidic fractionation, lactose depletion, and exoglycosidase digestion by PGC-ESI-MS/MS and MS^n^ and complemented this with permethylation to target sulfated MOs (*17*) (LC-MS/MS, MALDI-MS) for a representative acidic fraction. Overall, this represented a plethora of mass spectrometry measurements, allowing us to measure and quantify 332 unique milk oligosaccharides in seal milk, of which we structurally characterized 240 (Table S1, Fig. S1). This single study has thus made *H. grypus* the species with the second-most characterized MOs, behind *Homo sapiens* (Fig. 1a). As expected from the evolutionary loss of the CMAH gene in pinnipeds (*18*), we detected no Neu5Gc-containing glycan in *H. grypus* milk. Throughout lactation, ∼50% of the total abundance derived from fucosylated, non-sialylated, glycans, while up to 40% stemmed from sialofucosylated structures. Non-fucosylated, sialylated glycans remained low in abundance (1-4%). This preponderance of fucosylated structures was more reminiscent of human milk than of MOs from domesticated mammals, e.g., bovine MOs (*1*), yet also matched the characterized glycans in the closely related seal species *Phoca vitulina* (*11*).

**Figure 1.**
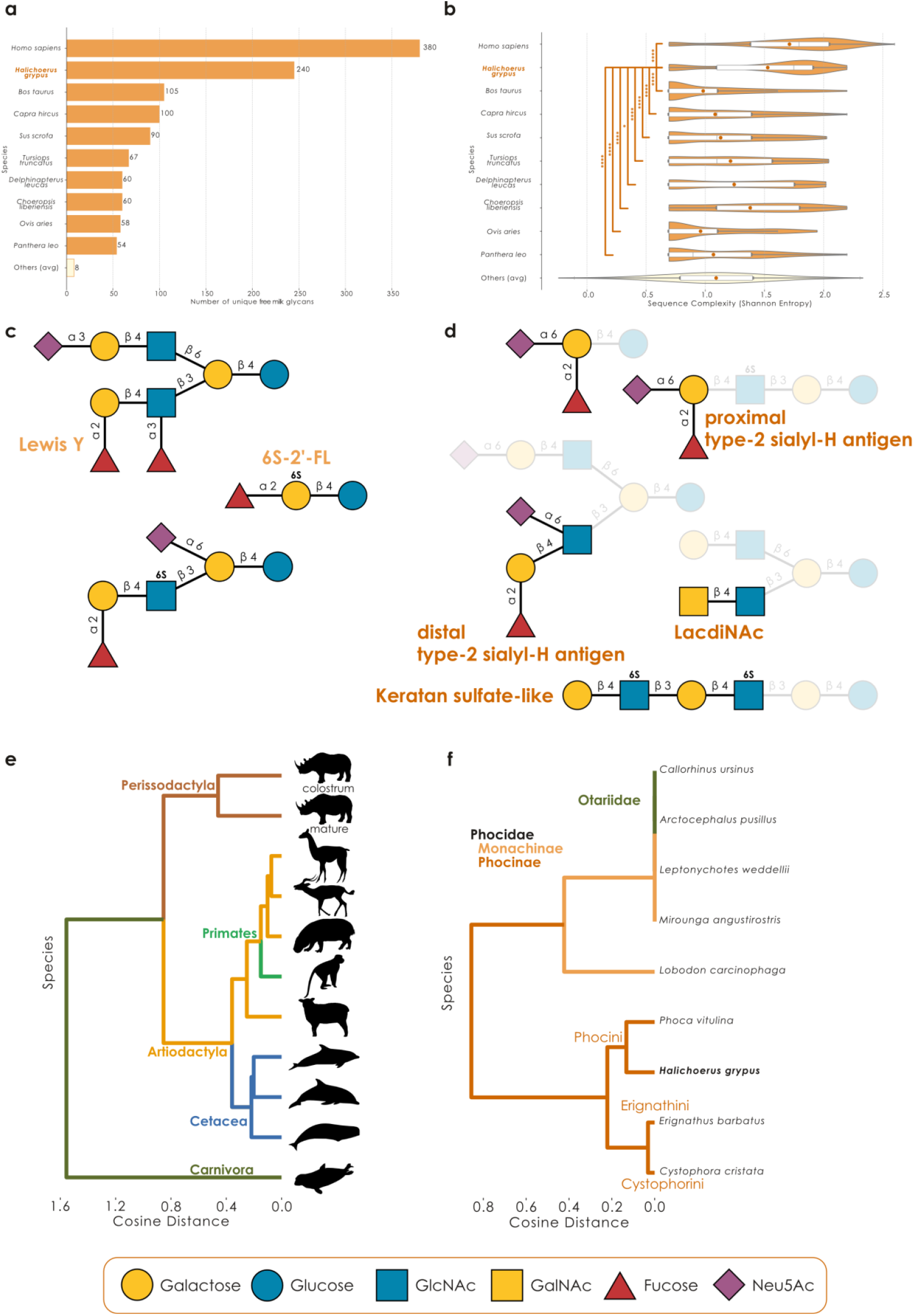
Grey seal milk exhibits a complex but characteristic glycome. **a)** Species with the most known free milk oligosaccharides. We displayed the 10 species with the most unique, structurally characterized MOs, as well as the average of the remaining species, via a bar graph. **b)** Alpha diversity (Shannon entropy) of structural epitopes, glycan sizes, and glycan branching across species. Data are shown as overlaid violin plots and boxplots. Lines in the boxplot indicate the median and dots the mean. The edges of the box describe the interquartile range (25% to 75%) and whiskers extend this to maximally 1.5x the interquartile range. Statistical comparisons between*H. grypus* and the other species have been performed as two-tailed Mann Whitney U-tests with a Benjamini-Hochberg correction. **c)** Representative structures identified in the milk of *H. grypus*, chosen because of either their novelty or exhibiting unusual motifs. **d)** New structural motifs identified in the milk of *H. grypus*. **e)** Phylogenetic tree of free milk glycomes. Pairwise cosine distances of motif abundances for all species with available comprehensive and quantitative milk glycomics (*3*) were used to create a dendrogram via UPGMA. **f)** Phylogenetic tree of seal milk glycomes. Dendrograms were constructed similarly to (e), yet only using the presence/absence of terminal motifs in all known seal milk glycans instead. All glycans in this article are drawn with GlycoDraw (*20*) and comply with the Symbol Nomenclature For Glycans (SNFG).

With 166 of the 240 structures (69%) being entirely novel—even accounting for ambiguities— we argued that seal milk still presented an entirely different sequence distribution from human MOs. A biodiversity analysis of structural epitopes, glycan sizes, and glycan branching across all species with measured MOs in the glycowork database (*4, 19*) indeed revealed that seal MOs also exhibited the second-highest alpha diversity, just behind *H. sapiens* (Fig. 1b). Given the drastic difference in sampling (decades of studying thousands of human milk samples with diverse methods vs 20 samples measured in one study), we even argue that *H. grypus* has the potential to eclipse human MOs in diversity and complexity, which would overturn the assumption of lower MO complexity in non-human species.

To that effect, we noted a range of unusual structures among our new discoveries (Fig. 1c), including the presence of novel or striking substructures/motifs, discussed further below. Especially noteworthy structures here included a sulfated version of the ubiquitous and, arguably most famous, MO 2’-fucosyllactose (2’-FL), Fucα1-2Gal6Sβ1-4Glc, which we have previously discovered in bottlenose dolphin milk (*3*). Sulfation frequently acts as an enhancer and modulator for binding specific lectins (*21*) and we speculated that 6S-2’-FL could present a more potent form of 2’FL for its typical anti-viral functions (*13, 22*). To that effect, we analyzed experimental glycan array data of 2’-FL and 6S-2’-FL and indeed uncovered both coronavirus as well as influenza virus proteins with strongly enhanced binding to 6S-2’-FL compared to 2’-FL (Fig. S2a, Table S2), supporting our hypothesis.

Additionally, we were excited to discover giant MOs in the milk of *H. grypus*, with several highly branched structures reaching 28 monosaccharides in length, such as Neu5Ac_6_Hex_12_HexNAc_10_ (Fig. S3; Table S3). This was substantially longer than the current record (*6*) of a human MO of size 18, and is among the largest mammalian non-polysaccharide glycans of any type (Table S4). We note that the scaffold of these mega-MOs allows for multivalent presentation of classic MO epitopes (Fig. S3), such as sialylation and/or fucosylation, and thus could make these molecules highly potent as soluble receptor decoys or signaling molecules themselves.

*H. grypus* uses lactose (Galβ1-4Glc) as a core, which is further elongated with multiple Galβ1-4GlcNAc (*N*-acetyllactosamine, LacNAc, type 2), namely poly-LacNAc. These poly-LacNAc chains serve as linear and extended scaffolds for diverse modifications, such as fucosylation, sialylation, and sulfation. Regarding noteworthy motifs/substructures in seal MOs, we first note an abundance of known structural motifs, such as the Sialyl-Lewis X, Lewis Y, type-2 H antigen, B antigen, Galili antigen, I antigen, and i antigen motifs (Fig. 1c, Table S1). Next, tying into our recent discovery of the LacdiNAc motif (GalNAcβ1-4GlcNAc) as a new and common MO building block (*3*), we report here that *H. grypus* milk also exhibits LacdiNAc-containing MOs (Fig. 1d, Fig. S1d), with 10 herein characterized seal MOs carrying this substructure (five of them entirely novel sequences; Table S5). We further note one, novel, structure with a sialylated LacdiNAc motif here as well.

Our in-depth mass spectrometry workflows also revealed the presence of two new terminal motifs, Fucα1-2(Neu5Acα2-6)Galβ1-4GlcNAc/Glc and Fucα1-2Galβ1-4(Neu5Acα2-6)GlcNAc (Fig. 1d, Fig. S1a, Fig. S1e), which, due to their biochemical provenance, we term proximal and distal type-2 sialyl-H antigen, respectively, referring to the glycan epitope that comprises blood group O, Fucα1-2Galβ1-4GlcNAc. Using the comprehensive glycan database within glycowork (*19*), we find that the proximal type-2 sialyl-H antigen has only been reported once so far, in glycosphingolipids of human colon adenocarcinoma (*23*), yet never in milk oligosaccharides. On the other hand, the distal type-2 sialyl-H antigen has only been described in glycosphingolipids of acute myeloid leukemia (*24*) and in MOs of another seal species (*11*), *Phoca vitulina*, raising the possibility (discussed below) of this as a more general seal motif. Both the H antigen and sialic acid moieties are potent anti-pathogen tools in MOs (*25*) and we speculated that their combination in the same molecules could enhance their potency.

Analyzing the curated glycan array binding data stored in the glycowork library (Fig. S2b, Table S2), we then indeed found that proximal sialylation changed the binding behavior of the type-2 H antigen (e.g., abrogating LEL binding, leaving UEA-I binding unaffected, and increasing AAL binding). In the case of distal sialylation, we again found increased AAL binding, as well as Siglec-1 and Siglec-15 binding (Fig. S2c, Table S2), supporting the biological relevance of these new motifs.

Sulfation is very common modification in seal MOs. Sulfated milk oligosaccharides have been reported as a minor constituent of HMOs, for instance in internal 6-sulfo Lewis X (*4, 26*). In our previous study, various mono-sulfated MOs were also detected in other mammals, including marine mammals (*3*). Here, we report for the first time the presence of keratan sulfate-like milk oligosaccharides (Fig. 1d, Fig. S1c), which could play key roles in supporting mucosal barrier function by mimicking host tissue glycosaminoglycans (*27*), fostering a GAG-degrading microbiome, or potentially aiding in growth factor signaling. Intriguingly, with the discovery of similar MOs with more repeat units, a new keratan sulfate subtype (*28*), such as KS-IV, would need to be created for keratan sulfate originating from a lactose core. In total, we noted 53 unique sulfated structures in *H. grypus* milk, 48 of which were entirely new (Fig. S4-5, Table S6), mostly via sulfation of the C6 of internal GlcNAc residues (Fig. S1b-c). This, by far, made *H. grypus* the species with the most known sulfated MOs (compared to the 22 characterized sulfated MOs in *H. sapiens* (*4*)). This also further highlighted the promise of applying the relatively new technique of sulfoglycomics to more milk samples and indicates that the prevalence of sulfated MOs has been underestimated thus far.

In general, non-sulfated seal MOs showed a deterministic pattern of extending sequences via branching and further decoration (Fig. 2), which then continued on to the measured giant MOs (Fig. S3; Table S3). Next to modulating binding, as mentioned above, we thus speculated that sulfation—changing the electrostatic and steric environment of a monosaccharide—may also affect glycan extension. Specifically with the case of GlcNAc6S, we indeed observed that MOs containing sulfated GlcNAc, usually connected to a branchpoint galactose, were less branched than those containing unmodified GlcNAc at this position (Fig. S4; Wilcoxon signed-rank test of extending sulfated/non-sulfated structures, controlled for their length: p < 0.001 that GlcNAc6S-modified structures are extended less often), leading us to the tentative conclusion that GlcNAc sulfation may be a mechanism in seal MOs to modulate and control branching, due to a change in substrate presentation to glycosyltransferases.

**Figure 2.**
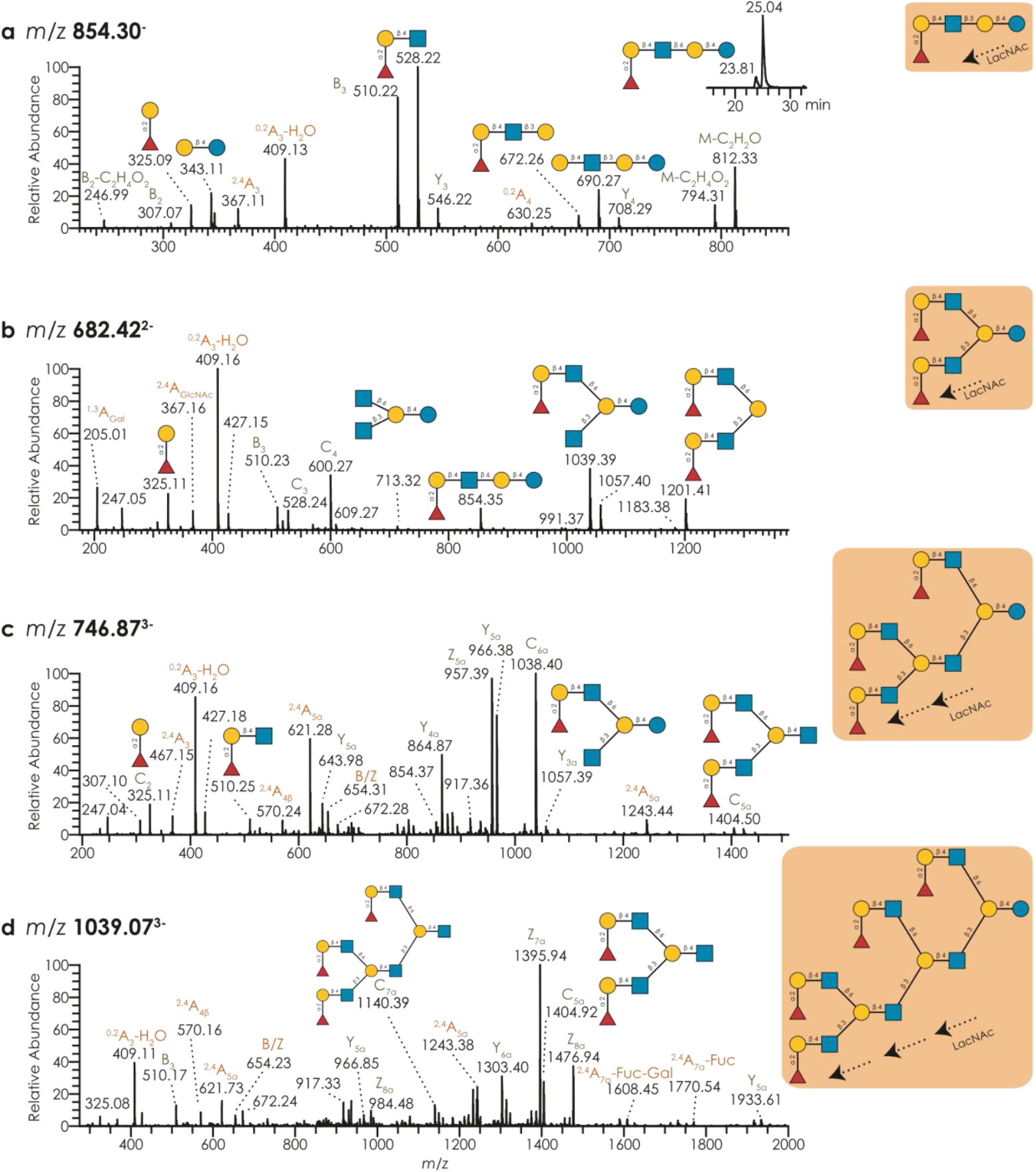
Seal milk oligosaccharides are extended in a repeating pattern of branched poly-LacNAc. **a-d)** The lactose is extended with one or multiple LacNAc units (a-d) and further branched with one or more LacNAc units (b-d). For the examples of neutral, fucosylated MO structures at *m/z* 854.3 (a), 682.42 (b), 746.87 (c), and 1039.07 (d), we show representative and annotated MS^2^ spectra, as well as the full determined sequence. Further, the chromatogram in (a) shows the separation of the two GlcNAc linkage isomers in retention time. MS/MS spectra of I-antigen containing structures are rich in B/C ions carrying this epitope (e.g., *m/z* 1201 in b, *m/z* 1038 and 1404 in c, and *m/z* 1140 and 1404 in d), ^2,4^A cleavage of GlcNAc adjacent to the branched Gal residue (e.g., *m/z* 621 in c and *m/z* 1770 in d), and D ions which indicate the size of the C6 branch caused by double cleavages of branched Gal residues (e.g., *m/z* 654 in c and d). All fragments in this work are provided in the Domon-Costello nomenclature (*29*).

In previous work (*3*), we have shown that our careful investigation of various milk glycomes resulted in well-comparable data and recapitulated DNA-based phylogenetic relationships between species. Since *H. grypus* is part of Carnivora, a taxonomic order we have not investigated in our prior work, we were curious to probe whether this would be reflected in our glycan-based phylogenetic tree. Indeed, *H. grypus* (as the only representative of Carnivora in our dataset) clustered separately from the ungulate clade (Artiodactyla and Perissodactyla; Fig. 1e), particularly driven by the high levels of type 2 H-antigen and Poly-LacNAc in seal milk, combined with the absence of motifs such as Sd^a^ (especially prevalent in Perissodactyla and Cetacea).

While our *H. grypus* dataset here presents the only quantitative seal MO dataset thus far, we were still interested in comparing the motif distributions of the various investigated seal species, to probe whether even finer taxonomic information can be found in their milk glycomes. For this, we used the absence/presence of motifs in identified milk glycans to calculate distances between seal milk glycomes, and construct a corresponding phylogenetic tree (Fig. 1f). Interestingly, this not only captured the main distinction within pinnipeds, of the families of eared seals (Otariidae) and earless seals (Phocidae)—diverging about 25 million years ago (*30*)—but even distinguished the Phocidae sub-families of Monachinae (34 chromosomes) and Phocinae (32 chromosomes). Especially for the Otariidae/Phocidae distinction, this is likely grounded in an alpha-lactalbumin mutation in the otariids that prevents them from forming elaborate MOs (*16*).

Among seals with investigated MOs, the closest neighbor to our *H. grypus* data in this clustering were *Phoca vitulina* seals. As both these species belong to the Phocinae sub-group of Phocini, we conclude that even fine-grained evolutionary information is reflected in the milk glycomes of these species. This clustering of *H. grypus* and *P. vitulina* was in part driven by their shared expression of the distal type-2 sialyl-H antigen, one of the unusual structures mentioned above (Fig. 1d). Overall, our glycan-derived seal taxonomy matched genomic phylogenies of this clade (*31*), especially considering that we lacked quantitative glycan data for most of these species. We thus conclude that milk glycans are exquisite repositories of evolutionary information, due to their adaptation to specific environmental niches.

### The seal milk glycome is changing throughout lactation

Next, we wanted to make use of the unique opportunity of having milk samples of the same seal individuals throughout their entire lactation period. For humans and cows, it has been shown repeatedly that the milk glycome is dynamic and changes over time (*32, 33*), to fit the changing needs of the infant. Since it was not known whether the same was true for seals, especially given their relatively brief lactation period, we next investigated whether our milk glycomes would cluster according to their time point (Table S7). For this, we used hierarchical clustering of CLR-transformed relative glycan abundances (Table S8), since glycomics data are compositional data which results in, otherwise unaccounted for, data dependencies (*34*). Overall, we observed a strong clustering of samples by their time point (Fig. 3a), substantiated by high clustering metrics, which indicated that the seal milk glycome also undergoes a concerted change during lactation that is conserved across individuals.

**Figure 3.**
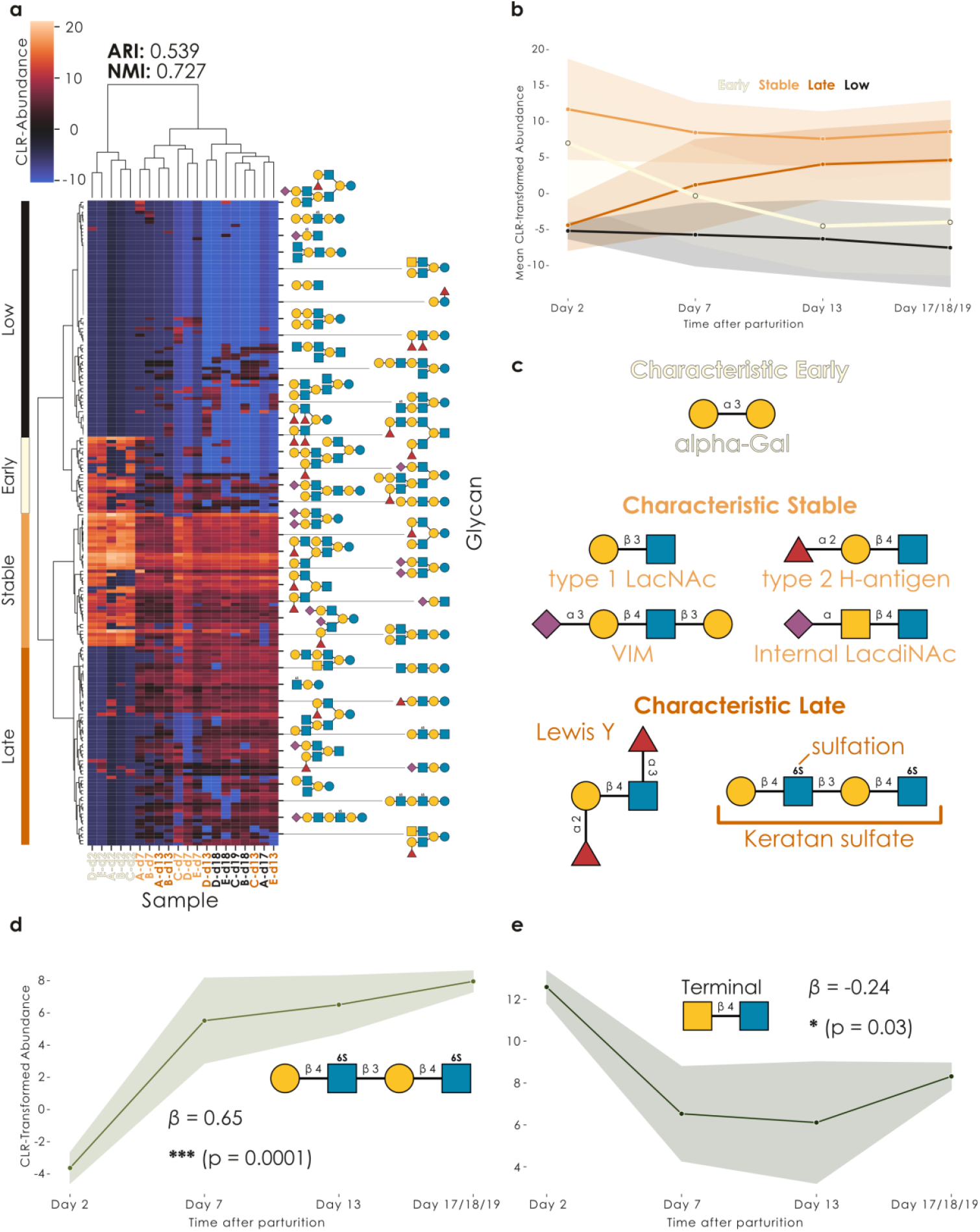
The seal milk glycome is changing during lactation. **a)** Seal milk samples cluster by lactation time point. Using CLR-transformed glycomics data from five individuals (A-E) and four timepoints (d2, d7, d13, d17/18/19; total N = 20), we engaged in hierarchical clustering of samples via Ward’s variance minimization algorithm. Representative glycans are shown for periodic rows. Successful clustering was assessed via the adjusted Rand index (ARI; ranging from -1 to 1) and normalized mutual information (NMI; ranging from 0 to 1). The four main glycan clusters resulting from row-wise clustering are labeled. **b)** Distinct clusters of glycans change abundance in a concerted manner during lactation. Data are shown as the abundance means of all glycans in the cluster for that timepoint, connected by a line plot with a 95% confidence band, shaded by cluster identity.**c)** Specific motifs characterize the temporally regulated glycans during lactation. For each cluster (Early, Stable, Late), we used the *quantify_motifs* function in glycowork (v1.5) (*19*) to obtain cluster-specific motif abundances and then determined which motifs were most cluster-specific via an ANOVA, using the *get_glycanova* function from glycowork. Representative, cluster-specific motifs are shown for each cluster via their SNFG depiction. All shown motifs are significantly different from all other clusters (p < 0.05), determined via Tukey’s Honestly Significant Difference (HSD) post-hoc tests, followed by a two-stage Benjamini-Hochberg correction for multiple testing. **d-e)** Terminal epitopes change consistently through lactation. We used the *get_time_series* function of glycowork (v1.5) to analyze the expression of keratan sulfate-like (d) and terminal LacdiNAc-containing glycans(e) throughout the lactation period. Shown are line plots of the CLR-transformed motif abundances, with a 95% confidence band, as well as the regression coefficient (*β*) and its significance, from fitting a degree 1 polynomial function to the time series. *p<0.05, ***p<0.001

We were next intrigued by the observation that several clusters of MOs seemed to exhibit distinct temporal dynamics (Fig. 3a), which we further investigated by analyzing the total glycan abundance in each cluster (the four top-level row clusters from the hierarchical clustering) over time (Fig. 3b). This analysis indeed led to the conclusion that there were three temporally interesting clusters (as well as a fourth group of glycans with very low and sporadic expression): a cluster of glycans exclusively expressed in early (colostrum-like) milk (Early), another cluster with relatively stably expressed MOs (Stable), and a last glycan cluster with late MOs that were not found in early milk and increased in abundance over the later stages of lactation (Late).

Reasoning that these different MO clusters fulfilled different functions in seal milk, similar to what is documented for human milk (*33*), we next set out to investigate which functional MO motifs distinguished each cluster from the others. We caution that, in addition, each cluster of course also contained many other motifs, yet these were not necessarily characteristic of that cluster. Overall, an ANOVA-based workflow of motif-level abundances allowed us to show that early MOs exhibited high levels of the alpha-Gal motif (Galα1-3Gal), while late glycans were enriched for the Lewis Y antigen and sulfated MOs, especially the mentioned keratan sulfate-like structures (Fig. 3c). We note that sulfated MOs in general have also been reported to increase in later lactation stages in cow milk (*35*). Lastly, the stable cluster was characterized by motifs such as the type 2 H-antigen and internal LacdiNAc structures. This analysis then confirmed our initial hypothesis that the different temporal clusters exhibited a unique array of functional moieties in their MOs, predisposing them for fulfilling different niche functions during pup development.

While analyzing these different temporal dynamics of the clusters can yield insights into the regulation of the lactation cascade, we next wanted to analyze how the milk glycome as a whole changed during lactation. With the example of fucosylation (Fig. S6), we revealed that some of its dynamics can be complex to disentangle, such as with a general decrease in fucosylated glycans during lactation, which was mainly driven by decreasing Fucα1-2Gal-containing glycans (important for blood group epitopes), even though Fucα1-3GlcNAc-containing glycans (important for Lewis antigen epitopes) exhibited the opposite trend.

Taking this analysis method to some of the motifs we newly discovered here, we for instance found, in accordance with our cluster analysis (Fig. 3c), that keratan sulfate-like MOs exhibited an increasing abundance even when considering the entire milk glycome (Fig. 3d). Further, we noted a general decrease in the abundance of LacdiNAc-terminated MOs during lactation (Fig. 3e), indicating that their levels seemed to be highest in colostrum-like milk. This decrease in LacdiNAc-containing MOs during lactation resembled a similar decrease in LacdiNAc which has been reported in protein-linked glycans in cow milk (*36*), raising interesting possibilities of cross-connections between different glycan types in milk during lactation.

Lastly, we returned to our milk glycome clustering (Fig. 3a) to make another observation: Next to temporal clusters of MOs, it also seemed to us that the milk glycome of *H. grypus* became more diverse in general, in later lactation stages. We formally analyzed this via three different alpha diversity indices of the milk glycomes and conclusively confirmed this observation (Fig. S7a-c). Next, we wanted to make sure that the clustering of the samples on our heatmap was not only due to this increase in diversity and engaged in an ANOSIM analysis of the beta diversities of our milk glycomes (Fig. S7d), which confirmed that the glycan sequence content also changed throughout lactation, supported by our earlier analyses as well. Overall, we report that the seal milk glycome (i) becomes more diverse throughout lactation, (ii) changes its repertoire of available functional groups, and (iii) exhibits clusters of glycans with concerted changes, e.g., only present during the early phase, that hint at a regulated and conserved process.

### The changing seal milk metabolome is reflected in the glycome

Since the same seal milk samples have been used for metabolomics measurements in an earlier study (*15*), we next set out to investigate whether we could find links between the milk metabolome (Table S9) and the milk glycome in *H. grypus*, given that glycosylation is metabolically regulated (*37*). Using our established approach of cross-correlating CLR-transformed systems biology datasets (*34*), we did indeed find many significant correlations of glycan substructures/motifs and milk metabolites throughout the lactation period (Fig. 4a).

**Figure 4.**
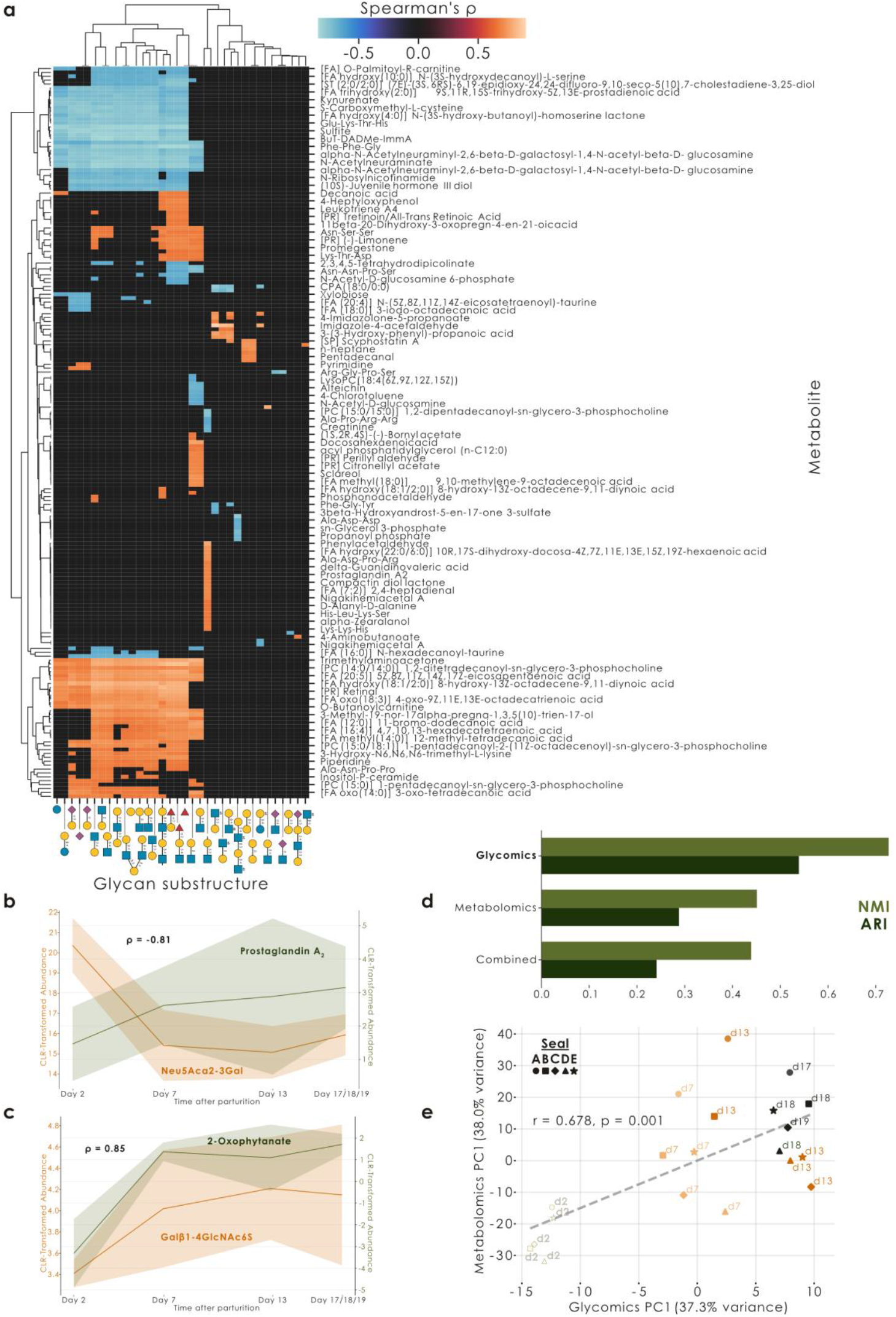
The changing milk metabolome is reflected in the glycome. **a)** Seal milk glycan substructures and metabolites correlate throughout the lactation period. We used the *get_SparCC* function from glycowork (v1.5) for a cross-correlation analysis of CLR-transformed glycomics and metabolomics data. Shown is a hierarchical clustering of the resulting Spearman’s ρ correlation coefficients, with only significant correlation coefficients shown (p < 0.05 of two-tailed *t*-tests, corrected for multiple testing by a two-stage Benjamini–Hochbergprocedure). **b-c)** Features of the milk glycome and metabolome are strongly correlated during lactation. For the example of Neu5Acα2-3Gal / Prostaglandin A_2_ (b) and Galβ1-4GlcNAc6S / 2-Oxophytanate (c), we show their CLR-transformed abundance during the lactation period (line: mean, confidence band: 95% CI), as well as their regularized partial correlation as Spearman’s ρ. **d)** Milk glycomes are more characteristic of lactation stage than milk metabolomes. Metrics (normalized mutual information, NMI; adjusted Rand index, ARI) are shown for clustering lactation timepoints by using CLR-transformed glycomics, metabolomics, or glycomics+metabolomics data. **e)** Joint glycome-metabolome transitions mark lactation progression. Principal component analysis (PCA) was performed on CLR-transformed glycomics and metabolomics data. Correlation between glycomics PC1 (37.3% variance) and metabolomics PC1 (38.0% variance) is shown as Pearson’s r. Samples are colored by sampling day (d2-d19) and exhibit different shapes for each seal (A-E).

Given the high degree of overlap between shared glycan substructures, as well as biosynthetically related metabolites, we then refined this by calculating regularized partial correlations, correcting for “bystander” correlations and enriching for more direct effects (Table S10). Overall, we noticed strong positive correlations (Spearman’s ρ of 0.7-0.9) between several MO substructures and membrane lipids (phosphatidylcholines, phosphatidylethanolamines, phosphatidylinositols, sphingolipids) as well as fatty acid components (acylcarnitines), which could indicate the coordinated delivery of energy via milk fat and developmental/protective factors via the MOs.

Specific regularized partial correlations for instance included a strong negative correlation of sialylated MOs (Neu5Acα2-3Gal) and the anti-inflammatory (*38*) eicosanoid prostaglandin A_2_(Fig. 4b), with the latter increasing in later stages of lactation. Additionally, we noted a strong positive correlation of sulfated glycans (Galβ1-4GlcNAc6S) and 2-oxophytanate (Fig. 4c), a metabolite produced during the oxidation of phytanic acid, derived from a fish diet (*39*). This could be a product from lipid catabolism, since grey seals are capital breeders and fast for the entire lactation period. Increasing lipid catabolism is known to increase oxidative stress, whereas sulfated glycans have been linked to mitigating oxidative stress (*40*), creating an intriguing connection for future research into sulfated glycans as a protective factor here.

Given that the milk metabolome is also known to change over time (*41*), we wanted to compare which systems biology modality, the milk glycome or metabolome, carried clearer information about the lactation stage. Similar to our previous clustering analysis (Fig. 3a), we thus clustered samples by their glycome, metabolome, or both (Fig. 4d). Interestingly, this resulted in the glycome being the most informative modality for determining lactation stage (ARI: 0.539, NMI: 0.727), even superior to the combination of glycome and metabolome (ARI: 0.228, NMI: 0.429). We suspect that this latter result was due to the added noise by the metabolome, because PCA-driven denoising indeed improved the clustering by the combined features (ARI: 0.399, NMI: 0.550), yet these metrics still did not reach the performance of the milk glycome by itself, perhaps because the metabolome clustered by individual, rather than lactation stage. We note that there is still a shared component of variation in both ‘omics layers that correlates similarly with lactation stage (Fig. 4e), explaining ∼38% of variance, and loading highest on our keratan sulfate-like MOs, distal sialyl-H antigen, and fatty acids, respectively, indicating a shared physiological program.

Overall, this discrepancy in information content—despite the metabolome exhibiting ∼5x the number of features of the glycome—clearly indicates the (i) physiological relevance and (ii)concerted regulation of the MOs during lactation, as well as the incredible information richness of glycans in general.

### Changing structures in seal milk glycome exhibit immunomodulatory and anti-biofilm properties

Based on the prominence of LacdiNAc-containing structures in our seal milk samples (Fig. 1)—and the relative novelty of finding this motif to be conserved in milk oligosaccharides (*3*)—we decided to more closely investigate functional properties of this class of molecules. MOs have known and potent effects on modulating immune cell activity, which often is even conserved across species (*3, 42*). In absence of an available *H. grypus* immune cell line, we thus tested the impact of LacdiNAc on human macrophages.

As a baseline for comparisons, we first differentiated THP-1 monocyte cells into naïve macrophage-like cells (M0 macrophages). These were then activated with LPS for classical (M1) activation or IL-4/IL-13 for alternative (M2) activation, leading to distinct cytokine profiles across macrophage populations (Fig. 5a). Next, we co-treated naïve, M1-, and M2-stimulated macrophages with LacdiNAc (GalNAcβ1-4GlcNAc) and the extremely closely related LacNAc (Galβ1-4GlcNAc) moiety. In addition to these disaccharides, we synthesized a full LacdiNAc-containing MO (LdiNnT, lacto-*N,N*-neotetraose), for comparison with the conserved MO LNnT (lacto-*N*-neotetraose), differing by only one *N*-acetyl group.

**Figure 5.**
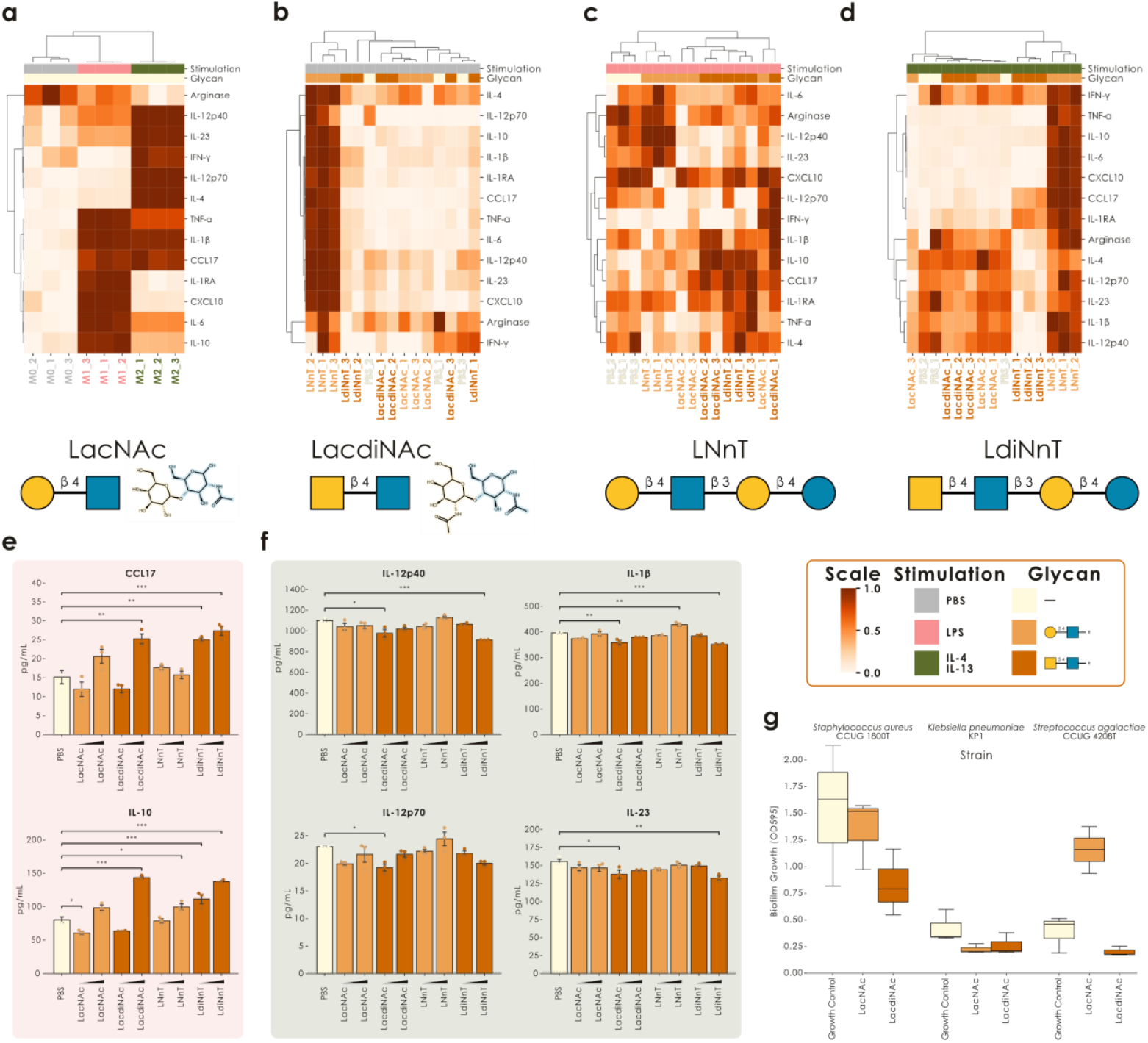
LacdiNAc exhibits unique immunomodulatory and anti-biofilm effects. **a)** Cytokine profile of naïve (M0, PBS) vs M1-polarized (LPS) vs M2-polarized (IL-4/IL-13) macrophages. **b-d)** Cytokine profiles showing the immunomodulatory effects of LacNAc/LacdiNAc and LNnT/LdiNnT on M0- (b), M1- (c), and M2-polarized(d) macrophages. Row dimensions were normalized to 0-1 before clustering by correlation distance. **e)** Quantification of CCL17 and IL-10 cytokine production upon 0.1 – 1 mM LacNAc/LacdiNAc and LNnT/LdiNnT treatment of M1-polarized macrophages. **f)** Quantification of IL-12p40, IL-12p70, IL-1β, and IL-23 cytokine production upon 0.1 – 1 mM LacNAc/LacdiNAc and LNnT/LdiNnT treatment of M2-polarized macrophages.The dashed line indicates the limit of detection as determined by the standard curve of each analyte. **g)** Biofilm formation (measured as OD595 absorption, adjusted for blank medium controls) of three bacterial strains grown in the absence or presence of 1 mg/mL LacNAc or LacdiNAc. Significant differences were established via a one-way ANOVA with post-hoc Tukey’s HSD tests. ***, p < 0.001; **, p < 0.01; *, p < 0.05

With the possible exception of LNnT, which was produced by fermentation and thus may contain trace endotoxins, treatment with (physiologically relevant concentrations (*43*) of) LacNAc/LacdiNAc structures had little to no effect on cytokine production in naïve macrophages (Fig. 5b, Fig. S8). In contrast, we observed numerous significant immunomodulatory effects of LacdiNAc-containing structures (both *N,N*-acetyllactosamine and lacto-*N,N*-neotetraose) but not LacNAc-containing structures (*N*-acetyllactosamine and lacto-*N*-neotetraose) in activated macrophages. Specifically, in M1-polarized macrophages, LacdiNAc upregulated CCL17 and IL-10 (Fig. 5c, Fig. 5e), while in M2-polarized macrophages, it downregulated IL-12p40, IL-12p70, IL-1β, and IL-23 (Fig. 5d, Fig. 5f). All results can also be found in Table S11.

Previously, LacdiNAc has been proposed as a parasite pattern and as a ligand for galectin-3 on macrophages (*44*). We contend here, however, that galectin-3 cannot be viewed as the only LacdiNAc receptor responsible for our observed immunomodulation because: (i) the mentioned work did specifically not identify galectin-3-mediated effects in the herein used THP-1 cells; and (ii) galectin-3 in fact bound LacNAc with at least similar affinity to LacdiNAc (*44*), while we observed significant differences in the effects of LacNAc vs LacdiNAc in our assay. We thus hypothesize that another LacdiNAc receptor exists on THP-1 cells that contributes to the immunomodulatory effect of this glycan moiety.

Next to immunomodulatory effects, an emerging property of MOs in recent years has been the inhibition of biofilm formation (*45–47*), with important implications for antibiotic resistance development, which we can confirm here, for instance with the well-characterized 6’-sialyl-lactose (Fig. S9). Since LacdiNAc-containing glycans have never been assessed in this regard to the best of our knowledge, we continued our comparison of LacNAc and LacdiNAc with respect to their action on biofilms. Assessing three pathologically relevant bacterial strains (*Klebsiella pneumoniae* KP1, *Streptococcus agalactiae* CCUG 4208T, *Staphylococcus aureus* CCUG 1800T), we report that LacdiNAc—but not LacNAc—inhibited biofilm formation in *S. agalactiae* and *S. aureus*, while both glycan moieties inhibited *K. pneumoniae* biofilm formation (Fig. 5g). Importantly for potential resistance development, none of the added glycan moieties affected bacterial growth (Fig. S10), potentially indicating either a signaling-mediated effect of the added glycan moieties or an inhibition of biofilm-associated lectins.

These encouraging findings implied that (i) more anti-pathogenic molecules are still available in mammalian milk for biomining purposes and (ii) minute chemical differences, such as between LacNAc and LacdiNAc, coupled with pronounced biological differences, speak to a specific recognition mechanism mediating this effect, which could eventually be targeted. It is also interesting to note here that mucin *O*-glycans, which can contain similar epitopes, have been speculated to modulate *S. aureus* virulence and adhesion *in vivo* (*48, 49*). Therefore, we continued this line of research by probing the anti-biofilm properties of the recently discovered glucuronyl-lactose (*3*) as another new MO, which then demonstrated promising anti-biofilm properties in *K. pneumoniae* and *S. aureus* that could not be observed with free glucuronic acid (Fig. S11). We thus conclude that anti-biofilm properties could be common in yet-to-be-discovered MOs and deem this a promising reservoir for mining anti-pathogenic compounds.

## Discussion

With an in-depth case study of the multifaceted and longitudinal milk glycome of the Atlantic grey seal, *Halichoerus grypus*, we here show that human-level complexity of milk oligosaccharide biochemistry and regulation can be found elsewhere in the animal kingdom, somewhat revising currently held paradigms of the exceptional state of human breast milk. We also note that we here, yet again, extend the number of all known MO structures by >20%, demonstrating how much remains to be learned about glycan biosynthesis and biodiversity. Finally, the highly potent functional properties of structural motifs in these newly discovered MOs, if nothing else, should motivate the further exploration of the MO repertoires of more mammals.

As noted previously (*3, 9*), aquatic and semi-aquatic mammals (such as *H. grypus*) exhibit a pronounced MO complexity and we envision that future efforts to map the pan-mammalian milk glycome will benefit most from investigating such species. Our efforts here notwithstanding, the current rate of discovering >50% new sequences in each newly characterized mammal indicates that there is still much to be discovered in the biochemical diversity of milk oligosaccharides, which could also shed light on biosynthetic constraints and yield functional molecules with biomedical relevance.

We especially note that the rather common properties of new MO building blocks as precision probes for biomedically relevant applications such as immunomodulation or anti-biofilm action is reminiscent of the recently emerging consensus of mucin glycans as anti-virulence signaling molecules (*50, 51*). Particularly given their role as already soluble molecules, MOs are not only soluble decoys for viruses and bacteria but also prime signaling molecules with established functions, and given the partly shared sequence content with mucin *O*-glycans, we envision that there could be substantial synergy in studying these anti-pathogen signaling functions across mucin glycans and MOs. Our findings, such as increased binding of the new 6S-2’-FL to the dendritic cell immunoreceptor 1 (DCIR1) compared to 2’-FL (Fig. S2a), provide ample opportunities for starting this line of inquiry.

Deep quantitative glycomics datasets of non-human samples are still a rarity and—especially with (i) longitudinal data, (ii) milk, and (iii) stemming from several individuals such as in our work here—present a strong point of novelty of our study and enable the kind of sophisticated analyses that constitute the future of glycomics (*52*). Crucially, this provides an exhaustive overview of available glycan epitopes in the free milk glycome. While this already facilitates analyses across, e.g., neutral and acidic fractions of MOs, we envision that future efforts will also contribute the corresponding *N*- and *O*-glycans from proteins in these milk samples. This could, for instance, yield insights into global vs class-specific regulation of glycan expression. We envision that more such multi-glycomics datasets, enabled by new workflows (*53*), will thus yield a more exhaustive understanding of the regulation, dynamics, and functions of the milk glycome and its roles in physiology.

## Methods

### Sample processing

Milk samples were originally collected in a previous study (*14*) from a seal colony on the Isle of May, Scotland. Sample processing began with diluting 500 μL of breast milk with an identical volume of distilled water, followed by centrifugation at 4000g for 30 minutes at 4°C and a removal of the fat layer. We then added two volumes of cold 96% ethanol to one volume of skimmed milk, followed by overnight incubation at 4°C to achieve protein precipitation. The sample underwent another centrifugation at 4000g for 30 minutes at 4°C, after which the supernatant containing milk oligosaccharides (MOs) was transferred to a fresh tube, frozen, and lyophilized. The protein pellet was kept for protein-linked glycan analysis.

Lyophilized milk oligosaccharides were resuspended in water at a concentration of 250 μL per mL of original milk. Residual proteins were eliminated using a spin-filter with a 10 kDa cutoff, operated at 11,000 rpm for 10 minutes (Sigma-Aldrich). Reduction for all glycan types was performed overnight at 50°C using 0.5 M NaBH_4_ and 20 mM NaOH. Desalting was accomplished using cation exchange resin (AG50WX8, Bio-Rad) packed onto a ZipTip C18 tip (Sigma-Aldrich). Following SpeedVac drying, methanol was introduced to remove any remaining borate through evaporation. The samples were subsequently resuspended in water at 250 μL per mL of original milk. The resulting glycans were analyzed via LC-MS/MS (see below) using 3 μL per measurement, with or without additional fractionation.

Further fractionation of the released MOs into neutral and acidic components was achieved using DEAE Sephadex A-25 (GH Healthcare). Since lactose, the predominant MO in the neutral fraction, would suppress other minor neutral MO signals during LC-MS/MS, it was eliminated using a carbon solid-phase cartridge (HyperSep Hypercarb SPE cartridges 25 mL, Thermo Scientific, Sweden). The cartridge was prepared with three 500 μL washes of 90% MeCN containing 0.1% TFA, followed by three 500 μL washes of 0.1% TFA. After MO application, lactose was eluted using three 500 μL washes of 8% MeCN with 0.1% TFA. The neutral MOs were subsequently eluted using three 500 μL washes of 65% MeCN containing 0.1% TFA, dried via centrifugation evaporation, and stored at -20°C until analysis, maintaining a concentration of 250 μL per ml of original milk.

### Glycomics

LC-MS/MS analysis of all glycan types was performed from stock solutions containing 250 μL sample per mL of original milk. Glycans, at 3 μL per sample in water, underwent separation on an in-house packed column (10 cm × 250 μm) containing 5 μm porous graphitized carbon particles (Hypercarb; Thermo-Hypersil). Following injection, elution used an acetonitrile gradient (Buffer A: 10 mM ammonium bicarbonate; Buffer B: 10 mM ammonium bicarbonate in 80% acetonitrile). The gradient, ranging from 0-45% Buffer B, ran for 46 minutes, followed by a 100% Buffer B wash step and 24-minute Buffer A equilibration.

Sample analysis was conducted in negative ion mode using an LTQ linear ion trap mass spectrometer (Thermo Electron), equipped with an IonMax standard ESI source featuring a stainless-steel needle maintained at -3.5 kV. Compressed air served as the nebulizer gas, while the heated capillary was maintained at 270°C with a -50 kV capillary voltage. The process included a full scan (*m/z* 340 or 380-2000, two micro scans, maximum 100 ms, target value of 30,000) followed by data-dependent MS^2^ scans (two micro scans, maximum 100 ms, target value of 10,000) using normalized collision energy of 35%, an isolation window of 2.5 units, activation q = 0.25, and 30 ms activation time. The MS^2^ threshold was set at 300 counts. Data acquisition and processing employed Xcalibur software (Version 2.0.7). Comparison of glycan abundances between samples involved quantifying individual glycan structures relative to total content by integrating the extracted ion chromatogram peak area. The area under the curve (AUC) for each structure was normalized to the total AUC and expressed as a percentage. Peak area processing was performed using Progenesis QI (Nonlinear Dynamics Ltd). MOs were identified from their MS/MS spectra by manual annotation together with exoglycosidase verification as described previously (*3, 54, 55*). Detected glycans and where to find them for the neutral and acidic fractions are recorded in Tables S12-13.

### Permethylation and MS data acquisition and processing

One of the MO samples (JC_231115MA5; seal B, day 7, acidic fraction) was permethylated and then fractionated by a Waters® OASIS MAX cartridge into neutral, mono-sulfated, and multiply sulfated glycan pools, exactly as described before (*56*). Each of the permethylated MO sample fractions was initially screened by MALDI-MS to obtain an overall profile and to identify the major structures present by glycan compositions. Sample aliquots were mixed 1:1 with matrix (2,5-dihydrobenzyonic acid for positive mode analysis of non-sulfated glycans, and 10 mg/mL of 3,4-diaminobenzophenone for sulfated glycans in negative mode) in 50% acetonitrile and spotted onto the MALDI plate for data acquisition on an AB SCIEX MALDI TOF/TOF 5800 system. For nanoLC-nanoESI-MS analysis, the MO samples were dissolved in 10% acetonitrile/0.1% formic acid, applied via autosampler to an Ultimate™ 3000 RSLC system connected to an Orbitrap Fusion™ Tribrid™ Mass Spectrometer (ThermoFisher Scientific) via a PicoView nanosprayer (New Objective, Woburn, MA), and separated with a constant flow rate of 300 nL/min at 50°C on a ReproSil-Pur 120 C18-AQ column (120 Å, 1.9 µm, 75 µm X 200 mm, Dr. Maisch). The solvent system used was buffer A (100% H_2_O with 0.1% formic acid) and buffer B (100% acetonitrile with 0.1% formic acid), with a 60 min linear gradient of 30% to 80% B. The Orbitrap Fusion Tribrid instrument settings were as described previously (*57*) using a HCD-MS^2^-product dependent MS^3^ data dependent acquisition method for positive mode, with full MS and HCD MS^2^ (stepped collision energy at 10, 15, 20) acquired in the Orbitrap at 120,000 and 30,000 resolution, respectively, and CID MS^3^ (30% normalized collision energy) in the ion trap for the targeted MS^2^ product ions at *m/z* 638.3382, 825.4227, and 999.5119, corresponding to Fuc_1_Hex_1_HexNAc_1_^+^,NeuAc_1_Hex_1_HexNAc_1_^+^,and Neu_1_Fuc_1_Hex_1_HexNAc_1_^+^, respectively. For analysis of sulfated MO in negative mode, the data dependent HCD-MS^2^ were acquired with a stepped collision energy at 45, 55, 65. All MS and MS/MS data were manually assigned according to previously established fragmentation patterns (*56–58*).

### Data analysis

All data preprocessing and motif analyses were performed using the Python package glycowork (*19*) (version 1.5). Analyses using relative abundances all followed the same preprocessing workflow, which corresponds to the *preprocess_data* function in glycowork: outlier datapoints were removed via Winsorization and missing data were imputed via a MissForest-based, machine learning-driven imputation strategy. Then, data were transformed via a center-log ratio (CLR) transform, using a γ value of 0.1, to correct for the compositional data nature of glycomics data (*34*). Glycan motifs were annotated via glycowork and motif abundances were obtained via the *quantify_motifs* function in glycowork. Our standard feature set here included “known” (named literature motifs) and “size_branch” (sizes and branching level of glycans), with any deviations from this noted in the respective figure legend.

### Synthesis of LacdiNAc-containing milk oligosaccharides

The total synthesis of LdiNnT from lactose was carried out through a sequential 13-step strategy involving glycosylation, deprotection, and purification. Full details on the experimental procedures, physical data, and ^1^H, ^13^C, ^1^H-^1^H COSY, and ^1^H-^13^C HSQC NMR spectra of all compounds are provided in the Supporting Information (SI).

### Immunomodulation of polarized macrophages

THP-1 cells (ATCC, TIB-202) were cultivated in Roswell Park Memorial Institute 1640 medium (RPMI 1640; Gibco, A1049101) supplemented with 10% (v/v) fetal bovine serum (FBS; Nordic Biolabs, FBS-HI-12A) and 1% (v/v) penicillin-streptomycin (Sigma-Aldrich, P4333-100ML) in a humidified atmosphere containing 5% CO_2_ at 37 °C. 4 × 10^4^ cells/well were seeded in a 96-well plate and differentiated into naïve macrophage-like cells (M0 macrophages) by treatment with 25 nM phorbol-12-myristate-13-acetate (PMA; Sigma-Aldrich, P8139-1MG) for 48 hours. After an additional 24-hour rest period in fresh medium, the cells were challenged with 100 ng/mL lipopolysaccharide (LPS; Sigma-Aldrich, L4391-1MG) (M1-polarized macrophages) or 20 ng/mL interleukin 4 (IL-4; Sigma-Aldrich, SRP3093-20UG) and 20 ng/mL interleukin 13 (IL-13; Sigma-Aldrich, SRP3274-10UG) (M2-polarized macrophages) in the absence or presence of 0.1 – 1 mM chemically pure MOs (*N*-acetyllactosamine: LacNAc, Sigma-Aldrich, A7791-5MG; *N,N*-acetyllactosamine: LacdiNAc, GlycoNZ, GNZ-0001-sp; lacto-*N*-neotetraose: LNnT, Elicityl, GLY021-95%; lacto-*N,N*-neotetraose: LdiNnT, synthesized in-house). After 24 hours, the culture supernatants were collected and analyzed for the levels of key macrophage cytokines using a multiplex immunoassay based on fluorescence-encoded beads (LEGENDplex; Biolegend, 740503) according to the manufacturer’s instructions. The samples were measured on an Accuri C6 Plus Flow Cytometer (BD) and the results were analyzed using LEGENDplex Data Analysis Software Suite (Biolegend, Qognit).

### Bacterial strains and crystal violet assay for biofilm quantification

*Staphylococcus aureus* CCUG 1800T and *Streptococcus agalactiae* CCUG 4208T were obtained from the Culture Collection University of Gothenburg (CCUG, https://www.ccug.se/). *Klebsiella pneumoniae* KP1 were obtained from Dr. Scott Rice (University of Technology, Sydney, Australia). *K. pneumoniae* KP1 and *S. aureus* CCUG 1800T were maintained in brain-heart infusion broth (Sigma-Aldrich, 53286-500G) and on brain-heart infusion agar. *S. agalactiae* CCUG 4208T were maintained in tryptic-soy broth (Sigma-Aldrich, 22092-500G) and on tryptic soy agar. All strains were preserved as glycerol stocks at -80°C.

The impact on biofilm formation by milk oligosaccharides were assessed by crystal violet staining. Overnight cultures from single colonies were diluted to a final OD_600_ of 0.01 (*K*. pneumoniae and *S. aureus*) and 0.1 (*S. agalactiae*) in a 96-well plate with a total well volume of 100 µL. Milk oligosaccharides (*N*-acetyllactosamine: LacNAc, Sigma-Aldrich, A7791-5MG; *N,N*-acetyllactosamine: LacdiNAc, GlycoNZ, GNZ-0001-PA; 2’-fucosyllactose: 2’-FL, Sigma-Aldrich, SMB00933-50MG; 6’-sialyl-d-lactose: 6’-SL, Sigma-Aldrich, 40817-1MG; D-glucuronic acid: GlcA, Sigma-Aldrich, G5269-10G; glucoronyllactose: GlcA-Lac, Elicityl) were added to a final concentration of 1 mg/mL. Plates were sealed with a breathable, sterile film and incubated statically for 24 h at 37°C to allow for biofilm formation. After 24 h, OD_600_ was measured using a Varioskan LUX Multimode Microplate reader. The planktonic cells were then removed, and each well was washed thrice with 150 µL 1×PBS. The remaining biomass was stained with a 0.1% crystal violet solution for 60 min at room temperature. After staining, the wells were washed once with 300 µL 1×PBS, followed by two washes with 150 µL 1×PBS. The stain was solubilized by adding 150 µL 33% acetic acid for 30 min and absorbance was read at 595 nm using a Varioskan LUX Multimode Microplate reader. The assay was performed once (n=1) for LacNAc, LacdiNAc, and 6’-SL, twice (n=2) for GlcA and GlcA-Lac, and thrice (n=3) for 2’-FL, with three technical replicates for each condition. All values were corrected by subtracting the values of the blanks, and the mean values was used for plotting.

### Statistical analysis

All statistical testing has been done in Python 3.12.6 using the glycowork package (version 1.5), the statsmodels package (version 0.14) and the scipy package (version 1.11). Data normalization and motif quantification was done with glycowork (version 1.5). All statistical operations on glycomics and metabolomics data have been performed on CLR-transformed data. Testing differences between two groups used Welch’s t-test, while testing differences between more than two groups used ANOVA or ANOSIM, followed by Tukey’s Honestly Significant Difference post-hoc tests in the former case. Regressions were performed via Spearman correlation analyses of the data, or of the residuals in the case of regularized partial correlations. All multiple testing correction has been performed via a two-stage Benjamini-Hochberg correction.

## Data availability

All data used in this article can either be found in supplemental tables or as stored datasets within glycowork (*19*). The glycomics MS raw files have been deposited in the GlycoPOST database under the ID GPST000556, with the corresponding retention times and compositions found in Tables S12-13.

## Code availability

Code and documentation are available via https://github.com/BojarLab/glycowork.

## Acknowledgement

The authors would like to thank Dr. Malcolm Kennedy for generous sample donations and James Urban for facilitating sample acquisition and transport. This work was supported by a Branco Weiss Fellowship – Society in Science awarded to D.B.; by the Knut and Alice Wallenberg Foundation; the Hasselblad Foundation; and the University of Gothenburg, Sweden. C.C. and R.H. gratefully acknowledge support from the Swiss National Science Foundation (project 320030-231409) and the University of Basel, Switzerland. We thank SciLifeLab and BioMS (Swedish research council) for providing financial support to the Proteomics Core Facility, Sahlgrenska Academy. K-H.K. was supported by Academia Sinica grant AS-IR-113-L04. We thank the Academia Sinica Common Mass Spectrometry Facilities for Proteomics and Protein Modification Analysis funded by the Academia Sinica Core Facility and Innovative Instrument Project grant AS-CFII-108-107, for MS data collection. The funders had no role in study design, data collection and analysis, decision to publish or preparation of the manuscript.

## Contributions

D. B. conceptualization; A.R.B., D. B., and J. L. formal analysis; C.C. and R.H. synthesis design & NMR analysis; C.C., D. B., J. L., and R.H. resources; D. B. data curation; A.R.B., D. B., C. J., and J. L. writing–original draft; A.R.B., C. C., D. B., C. J., J. B.-P., J. L., K.-H. K., M. D., R.H., and S.-Y. G. writing–review & editing; A.R.B., C. J., D. B., J. L., K.-H. K., and S.-Y.G. visualization; D. B., J. B.-P., K.-H. K., and R.H. supervision; D. B. funding acquisition; C. J., D.B., J. L., K.-H. K., M. D., and S.-Y. G. methodology; C. J., J. L, K.-H. K., and M. D. validation.

## References

1. T. Urashima, K. Ajisaka, T. Ujihara, E. Nakazaki, Recent advances in the science of human milk oligosaccharides. BBA Advances 7, 100136 (2025).

2. L. Bode, The functional biology of human milk oligosaccharides. Early Human Development 91, 619–622 (2015).

3. C. Jin, J. Lundstrøm, E. Korhonen, A. S. Luis, D. Bojar, Breast Milk Oligosaccharides Contain Immunomodulatory Glucuronic Acid and LacdiNAc. Molecular & Cellular Proteomics, 100635 (2023).

4. L. Thomès, V. Karlsson, J. Lundstrøm, D. Bojar, Mammalian milk glycomes: Connecting the dots between evolutionary conservation and biosynthetic pathways. Cell Reports 42, 112710 (2023).

5. J. Lundstrøm, D. Bojar, The evolving world of milk oligosaccharides: Biochemical diversity understood by computational advances. Carbohydrate Research 537, 109069 (2024).

6. C. A. Remoroza, Y. Liang, T. D. Mak, Y. Mirokhin, S. L. Sheetlin, X. Yang, J. V. San Andres, M. L. Power, S. E. Stein, Increasing the Coverage of a Mass Spectral Library of Milk Oligosaccharides Using a Hybrid-Search-Based Bootstrapping Method and Milks from a Wide Variety of Mammals. Anal. Chem. 92, 10316–10326 (2020).

7. T. Urashima, E. Taufik, K. Fukuda, S. Asakuma, Recent Advances in Studies on Milk Oligosaccharides of Cows and Other Domestic Farm Animals. Bioscience, Biotechnology, and Biochemistry 77, 455–466 (2013).

8. S. S. van Leeuwen, E. M. te Poele, A. C. Chatziioannou, E. Benjamins, A. Haandrikman, L. Dijkhuizen, Goat Milk Oligosaccharides: Their Diversity, Quantity, and Functional Properties in Comparison to Human Milk Oligosaccharides. J. Agric. Food Chem. 68, 13469–13485 (2020).

9. T. Urashima, R. Horiuchi, M. Sakanaka, T. Katayama, K. Fukuda, Lactose or milk oligosaccharide: which is significant among mammals? Animal Frontiers 13, 14–23 (2023).

10. T. Urashima, M. Arita, M. Yoshida, T. Nakamura, I. Arai, T. Saito, J. P. Y. Arnould, K. M. Kovacs, C. Lydersen, Chemical characterisation of the oligosaccharides in hooded seal (Cystophora cristata) and Australian fur seal (Arctocephalus pusillus doriferus) milk. Comparative Biochemistry and Physiology Part B: Biochemistry and Molecular Biology 128, 307–323 (2001).

11. T. Urashima, T. Nakamura, K. Yamaguchi, J. Munakata, I. Arai, T. Saito, C. Lydersen, K. M. Kovacs, Chemical characterization of the oligosaccharides in milk of high Arctic harbour seal (Phoca vitulina vitulina). Comparative Biochemistry and Physiology Part A: Molecular & Integrative Physiology 135, 549–563 (2003).

12. T. Urashima, T. Nakamura, D. Nakagawa, M. Noda, I. Arai, T. Saito, C. Lydersen, K. M. Kovacs, Characterization of oligosaccharides in milk of bearded seal (Erignathus barbatus). Comparative Biochemistry and Physiology Part B: Biochemistry and Molecular Biology 138, 1–18 (2004).

13. L. Thomès, D. Bojar, The Role of Fucose-Containing Glycan Motifs Across Taxonomic Kingdoms. Front. Mol. Biosci. 8, 755577 (2021).

14. A. D. Lowe, S. Bawazeer, D. G. Watson, S. McGill, R. J. S. Burchmore, P. P. Pomeroy, M. W. Kennedy, Rapid changes in Atlantic grey seal milk from birth to weaning – immune factors and indicators of metabolic strain. Sci Rep 7, 16093 (2017).

15. D. G. Watson, P. P. Pomeroy, M. W. Kennedy, Atlantic Grey Seal Milk Shows Continuous Changes in Key Metabolites and Indicators of Metabolic Transition in Pups From Birth to Weaning. Front. Mar. Sci. 7, 596904 (2021).

16. J. P. Avery, S. A. Zinn, Extraordinary diversity of the pinniped lactation triad: lactation and growth strategies of seals, sea lions, fur seals, and walruses. Animal Frontiers 13, 93–102 (2023).

17. S.-Y. Yu, S.-W. Wu, H.-H. Hsiao, K.-H. Khoo, Enabling techniques and strategic workflow for sulfoglycomics based on mass spectrometry mapping and sequencing of permethylated sulfated glycans. Glycobiology 19, 1136–1149 (2009).

18. M. O. Altman, P. Gagneux, Absence of Neu5Gc and Presence of Anti-Neu5Gc Antibodies in Humans—An Evolutionary Perspective. Front. Immunol. 10, 789 (2019).

19. L. Thomès, R. Burkholz, D. Bojar, Glycowork: A Python package for glycan data science and machine learning. Glycobiology, cwab067 (2021).

20. J. Lundstrøm, J. Urban, L. Thomès, D. Bojar, GlycoDraw: a python implementation for generating high-quality glycan figures. Glycobiology, cwad063 (2023).

21. J. Jung, J. R. Enterina, D. T. Bui, F. Mozaneh, P.-H. Lin, Nitin, C.-W. Kuo, E. Rodrigues, A. Bhattacherjee, P. Raeisimakiani, G. C. Daskhan, C. D. St. Laurent, K.-H. Khoo, L. K. Mahal, W. F. Zandberg, X. Huang, J. S. Klassen, M. S. Macauley, Carbohydrate Sulfation As a Mechanism for Fine-Tuning Siglec Ligands. ACS Chem. Biol. 16, 2673–2689 (2021).

22. Y. Yu, Y. Lasanajak, X. Song, L. Hu, S. Ramani, M. L. Mickum, D. J. Ashline, B. V. V. Prasad, M. K. Estes, V. N. Reinhold, R. D. Cummings, D. F. Smith, Human Milk Contains Novel Glycans That Are Potential Decoy Receptors for Neonatal Rotaviruses. Molecular & Cellular Proteomics 13, 2944–2960 (2014).

23. H. Korekane, S. Tsuji, S. Noura, M. Ohue, Y. Sasaki, S. Imaoka, Y. Miyamoto, Novel fucogangliosides found in human colon adenocarcinoma tissues by means of glycomic analysis. Analytical Biochemistry 364, 37–50 (2007).

24. D. Wang, T. Zhang, K. Madunić, A. A. De Waard, C. Blöchl, O. A. Mayboroda, M. Griffioen, R. M. Spaapen, C. G. Huber, G. S. M. Lageveen-Kammeijer, M. Wuhrer, Glycosphingolipid-Glycan Signatures of Acute Myeloid Leukemia Cell Lines Reflect Hematopoietic Differentiation. J. Proteome Res. 21, 1029–1040 (2022).

25. R. E. Moore, L. L. Xu, S. D. Townsend, Prospecting Human Milk Oligosaccharides as a Defense Against Viral Infections. ACS Infect. Dis. 7, 254–263 (2021).

26. Y. Guérardel, W. Morelle, Y. Plancke, J. Lemoine, G. Strecker, Structural analysis of three sulfated oligosaccharides isolated from human milk. Carbohydrate Research 320, 230–238 (1999).

27. P. S. Rawat, A. S. Seyed Hameed, X. Meng, W. Liu, Utilization of glycosaminoglycans by the human gut microbiota: participating bacteria and their enzymatic machineries. Gut Microbes 14, 2068367 (2022).

28. B. Caterson, J. Melrose, Keratan sulfate, a complex glycosaminoglycan with unique functional capability. Glycobiology 28, 182–206 (2018).

29. B. Domon, C. E. Costello, A systematic nomenclature for carbohydrate fragmentations in FAB-MS/MS spectra of glycoconjugates. Glycoconjugate J 5, 397–409 (1988).

30. K. Nyakatura, O. R. Bininda-Emonds, Updating the evolutionary history of Carnivora (Mammalia): a new species-level supertree complete with divergence time estimates. BMC Biol 10, 12 (2012).

31. J. W. Higdon, O. R. Bininda-Emonds, R. M. Beck, S. H. Ferguson, Phylogeny and divergence of the pinnipeds (Carnivora: Mammalia) assessed using a multigene dataset. BMC Evol Biol 7, 216 (2007).

32. E. M. Quinn, T. F. O’Callaghan, J. T. Tobin, J. P. Murphy, K. Sugrue, H. Slattery, M. O’Donovan, R. M. Hickey, Changes to the Oligosaccharide Profile of Bovine Milk at the Onset of Lactation. Dairy 1, 284–296 (2020).

33. C. Thum, C. R. Wall, G. A. Weiss, W. Wang, I. M.-Y. Szeto, L. Day, Changes in HMO Concentrations throughout Lactation: Influencing Factors, Health Effects and Opportunities. Nutrients 13, 2272 (2021).

34. A. R. Bennett, J. Lundstrøm, S. Chatterjee, M. Thaysen-Andersen, D. Bojar, Compositional data analysis enables statistical rigor in comparative glycomics. Nat Commun 16, 795 (2025).

35. T. E. Gray, K. Narayana, A. M. Garner, S. A. Bakker, R. K. H. Yoo, A. J. Fischer-Tlustos, M. Steele, W. F. Zandberg, Analysis of the biosynthetic flux in bovine milk oligosaccharides reveals competition between sulfated and sialylated species and the existence of glucuronic acid-containing analogues. Food Chemistry 361, 130143 (2021).

36. I. Gazi, K. R. Reiding, A. Groeneveld, J. Bastiaans, T. Huppertz, A. J. R. Heck, LacdiNAc to LacNAc: remodelling of bovine α-lactalbumin N-glycosylation during the transition from colostrum to mature milk. Glycobiology 34, cwae062 (2024).

37. A. F. Scheper, J. Schofield, R. Bohara, T. Ritter, A. Pandit, Understanding glycosylation: Regulation through the metabolic flux of precursor pathways. Biotechnology Advances 67, 108184 (2023).

38. T. Ohmura, Y. Tian, N. Sarich, Y. Ke, A. Meliton, A. S. Shah, K. Andreasson, K. G. Birukov, A. Birukova, Regulation of lung endothelial permeability and inflammatory responses by prostaglandin A2: role of EP4 receptor. MBoC 28, 1622–1635 (2017).

39. P. J. Brown, G. Mei, F. B. Gibberd, D. Burston, P. D. Mayne, J. E. McClinchy, M. Sidey, Diet and Refsum’s disease. The determination of phytanic acid and phytol in certain foods and the application of this knowledge to the choice of suitable convenience foods for patients with Refsum’s disease. J Human Nutrition Diet 6, 295–305 (1993).

40. H. Nagase, Y. Katagiri, K. Oh-hashi, H. Geller, Y. Hirata, Reduced Sulfation Enhanced Oxytosis and Ferroptosis in Mouse Hippocampal HT22 Cells. Biomolecules 10, 92 (2020).

41. K. O. Poulsen, F. Meng, E. Lanfranchi, J. F. Young, C. Stanton, C. A. Ryan, A. L. Kelly, U. K. Sundekilde, Dynamic Changes in the Human Milk Metabolome Over 25 Weeks of Lactation. Front. Nutr. 9, 917659 (2022).

42. F. Rosa, A. K. Sharma, M. Gurung, D. Casero, K. Matazel, L. Bode, C. Simecka, A. A. Elolimy, P. Tripp, C. Randolph, T. W. Hand, K. D. Williams, T. LeRoith, L. Yeruva, Human Milk Oligosaccharides Impact Cellular and Inflammatory Gene Expression and Immune Response. Front. Immunol. 13, 907529 (2022).

43. N. Sprenger, L. Y. Lee, C. A. De Castro, P. Steenhout, S. K. Thakkar, Longitudinal change of selected human milk oligosaccharides and association to infants’ growth, an observatory, single center, longitudinal cohort study. PLoS ONE 12, e0171814 (2017).

44. T. K. van den Berg, H. Honing, N. Franke, A. van Remoortere, W. E. C. M. Schiphorst, F.-T. Liu, A. M. Deelder, R. D. Cummings, C. H. Hokke, I. van Die, LacdiNAc-Glycans Constitute a Parasite Pattern for Galectin-3-Mediated Immune Recognition. J Immunol 173, 1902–1907 (2004).

45. D. L. Ackerman, R. S. Doster, J.-H. Weitkamp, D. M. Aronoff, J. A. Gaddy, S. D. Townsend, Human Milk Oligosaccharides Exhibit Antimicrobial and Antibiofilm Properties against Group B Streptococcus. ACS Infect. Dis. 3, 595–605 (2017).

46. A. Bhowmik, P. Chunhavacharatorn, S. Bhargav, A. Malhotra, A. Sendrayakannan, P. Kharkar, N. Nirmal, A. Chauhan, Human Milk Oligosaccharides as Potential Antibiofilm Agents: A Review. Nutrients 14, 5112 (2022).

47. R. E. Moore, S. K. Spicer, J. A. Talbert, S. D. Manning, S. D. Townsend, J. A. Gaddy, Anti-biofilm Activity of Human Milk Oligosaccharides in Clinical Strains of Streptococcus agalactiae with Diverse Capsular and Sequence Types. ChemBioChem 24, e202200643 (2023).

48. J. Ricciuto, S. R. Heimer, M. S. Gilmore, P. Argüeso, Cell Surface O-Glycans Limit Staphylococcus aureus Adherence to Corneal Epithelial Cells. Infect Immun 76, 5215–5220 (2008).

49. K. M. Jacob, S. Hernández-Villamizar, N. D. Hammer, G. Reguera, Mucin-induced surface dispersal of Staphylococcus aureus and Staphylococcus epidermidis via quorum-sensing dependent and independent mechanisms. mBio 15, e01562–24 (2024).

50. B. X. Wang, J. Takagi, A. McShane, J. H. Park, K. Aoki, C. Griffin, J. Teschler, G. Kitts, G. Minzer, M. Tiemeyer, R. Hevey, F. Yildiz, K. Ribbeck, Host-derived O-glycans inhibit toxigenic conversion by a virulence-encoding phage in Vibrio cholerae. The EMBO Journal 42, e111562 (2023).

51. J. Takagi, K. Aoki, B. S. Turner, S. Lamont, S. Lehoux, N. Kavanaugh, M. Gulati, A. Valle Arevalo, T. J. Lawrence, C. Y. Kim, B. Bakshi, M. Ishihara, C. J. Nobile, R. D. Cummings, D. J. Wozniak, M. Tiemeyer, R. Hevey, K. Ribbeck, Mucin O-glycans are natural inhibitors of Candida albicans pathogenicity. Nat Chem Biol 18, 762–773 (2022).

52. D. Bojar, F. Lisacek, Glycoinformatics in the Artificial Intelligence Era. Chem. Rev. 122, 15971–15988 (2022).

53. E. S. X. Moh, S. Dalal, N. J. DeBono, L. Kautto, K. Wongtrakul-Kish, N. H. Packer, SSSMuG: Same Sample Sequential Multi-Glycomics. Anal. Chem., acs.analchem.3c04928 (2024).

54. A. V. Everest-Dass, J. L. Abrahams, D. Kolarich, N. H. Packer, M. P. Campbell, Structural Feature Ions for Distinguishing N- and O-Linked Glycan Isomers by LC-ESI-IT MS/MS. J. Am. Soc. Mass Spectrom. 24, 895–906 (2013).

55. R. P. Estrella, J. M. Whitelock, N. H. Packer, N. G. Karlsson, Graphitized Carbon LC™MS Characterization of the Chondroitin Sulfate Oligosaccharides of Aggrecan. Anal. Chem. 79, 3597–3606 (2007).

56. H.-C. Tseng, C.-T. Hsiao, N. Yamakawa, Y. Guérardel, K.-H. Khoo, Discovery Sulfoglycomics and Identification of the Characteristic Fragment Ions for High-Sensitivity Precise Mapping of Adult Zebrafish Brain–Specific Glycotopes. Front. Mol. Biosci. 8, 771447 (2021).

57. C.-T. Hsiao, P.-W. Wang, H.-C. Chang, Y.-Y. Chen, S.-H. Wang, Y. Chern, K.-H. Khoo, Advancing a High Throughput Glycotope-centric Glycomics Workflow Based on NAnoLC-MS2-product Dependent-MS3 ANAlysis of Permethylated Glycans*. Molecular & Cellular Proteomics 16, 2268–2280 (2017).

58. C.-W. Cheng, C.-C. Chou, H.-W. Hsieh, Z. Tu, C.-H. Lin, C. Nycholat, M. Fukuda, K.-H. Khoo, Efficient Mapping of Sulfated Glycotopes by Negative Ion Mode nanoLC–MS/MS-Based Sulfoglycomic Analysis of Permethylated Glycans. Anal. Chem. 87, 6380–6388 (2015).

