## Supplemental Figures for "Seal milk oligosaccharides rival human milk complexity and exhibit functional dynamics during lactation"

### Supplementary Figures

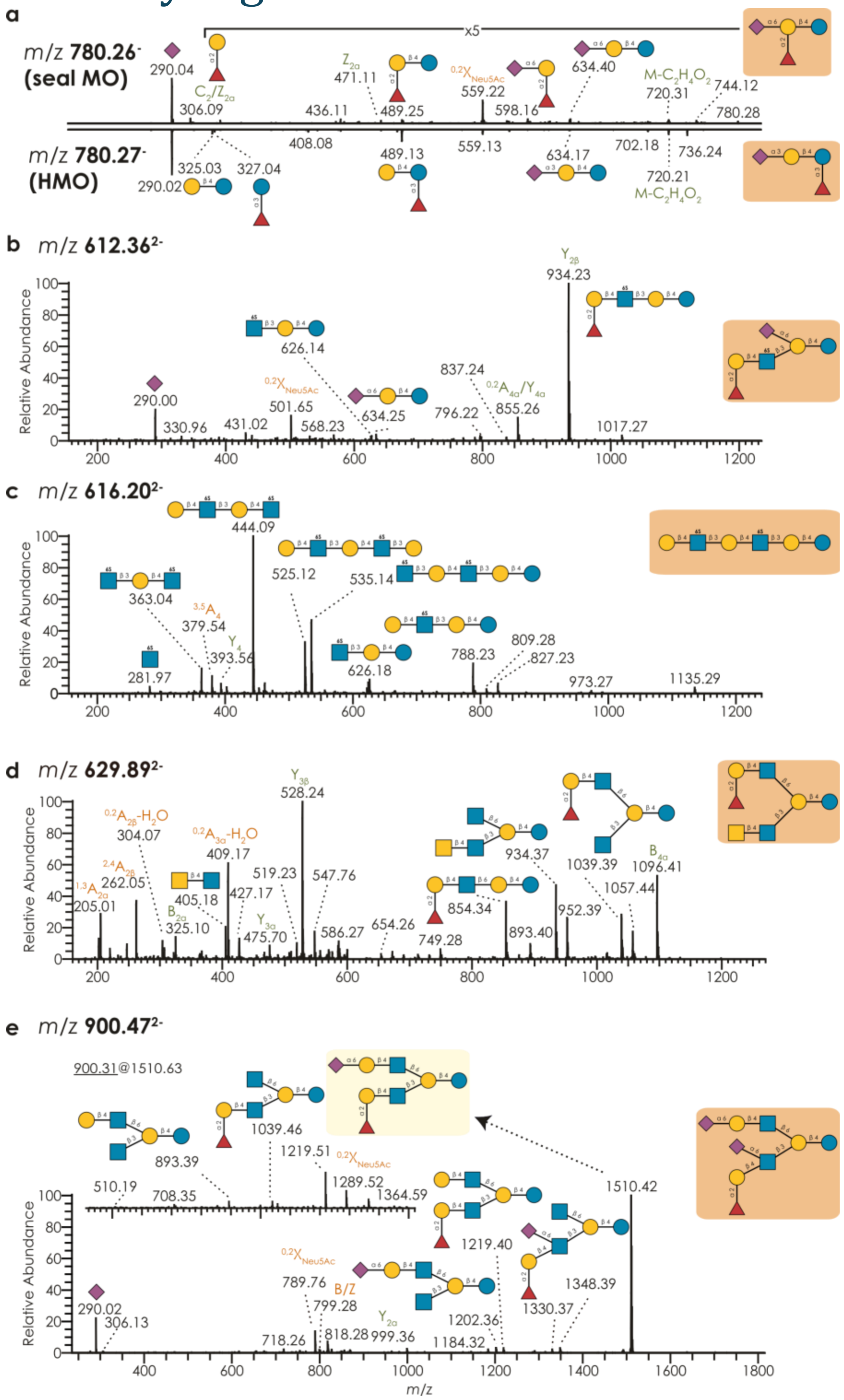

**Figure S1. Newly characterized milk oligosaccharides in grey seal milk. a-e)** For example novel structures of *m/z* 780.26 (a), 612.36 (b), 616.2 (c), 629.89 (d), and 900.47 (e), we show a representative, annotated $MS^2$ spectrum, along with their full determined sequence. An acquired $MS^3$ spectrum of the *m/z* 1510.63 fragment is further shown for (e) as an inlay. a) Two sialylated FLs are detected. One shows the same $MS^2$ spectrum as the human sialylated 3'FL. The $MS^2$ spectrum of the other contains diagnostic ions at *m/z* 306 and 559, suggesting α2,6-linked Neu5Ac and fragments at *m/z* 598 and 634 suggest doubly-substituted Gal. b-c) Fragments at *m/z* 501 ($^{0,2}X_{Neu5Ac}$) and 634 indicate α2,6-linked Neu5Ac (6'SL). Fragments at *m/z* 626 suggest sulfated GlcNAc from LNnT, while fragments at *m/z* 363 imply sulfation of both GlcNAc residues. d) LacdiNAc-containing MOs usually show $C_2$ ions at *m/z* 405 and $^{0,2}A_{GlcNAc}$-$H_2O$ ions at *m/z* 304. The H type 2 chain was diagnosed via *m/z* 205 ($^{1,3}A_{Gal}$), 427 ($^{0,2}A_{GlcNAc}$), and 409 ($^{0,2}A_{GlcNAc}$-$H_2O$) fragments. e) $MS^2$ spectrum of proximal type 2 sialyl-H. The size of C6 branch was assigned by D ions at *m/z* 799. The $^{0,2}X_{Neu5Ac}$ at *m/z* 789 ($MS^2$) and 1289 ($MS^3$) highly indicate α2,6-linkage of Neu5Ac.

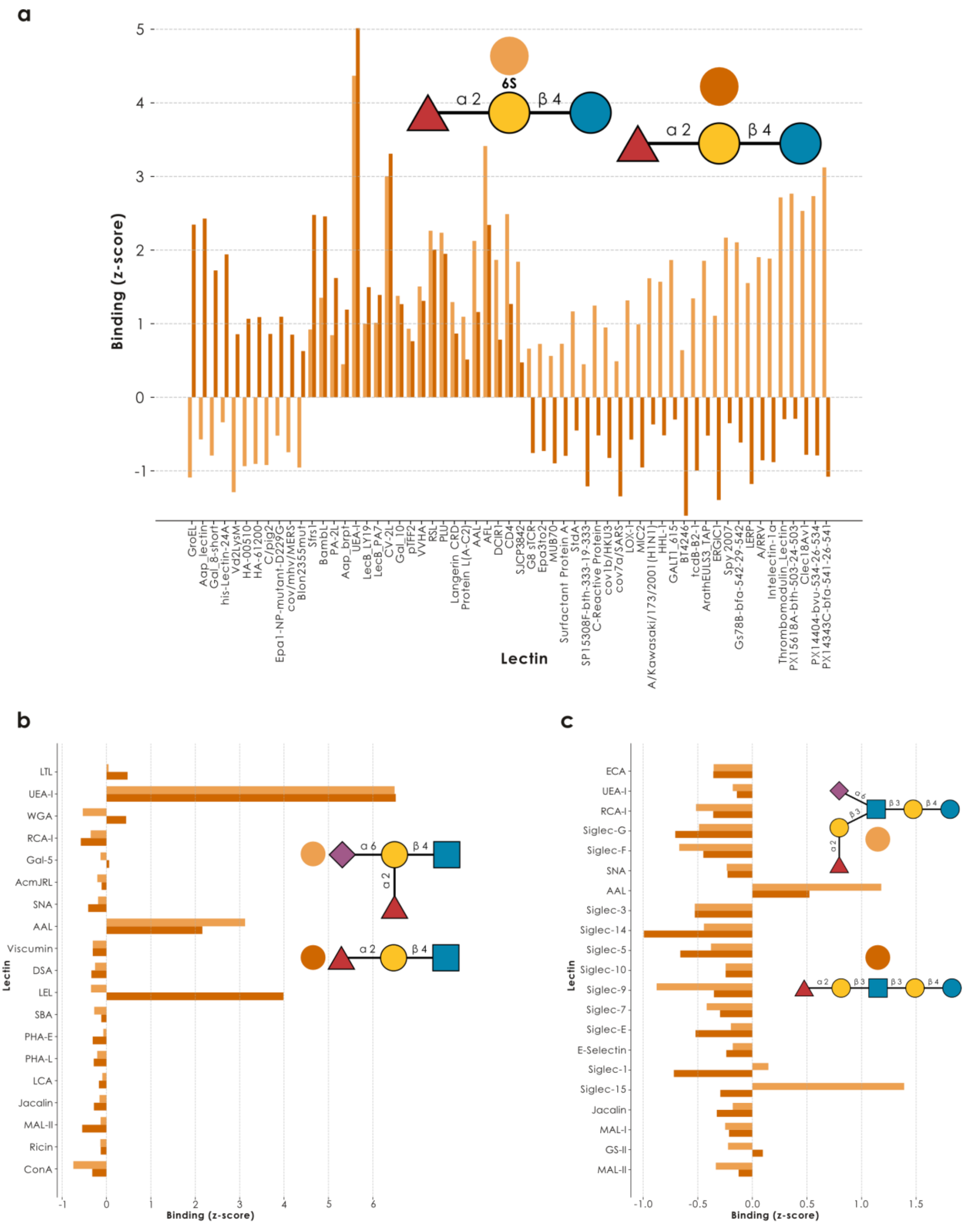

**Figure S2. New milk oligosaccharide modifications affect lectin binding. a-c)** For 6-sulfo-2'-fucosyllactose (6S-2'-FL; a) as well as proximal (b) and distal (c) type-2 sialyl-H antigen, we obtained their glycan array binding values to those lectins with information for both molecules. For 6S-2'FL, we then further filtered this by at least one molecule exhibiting a z-score above zero.

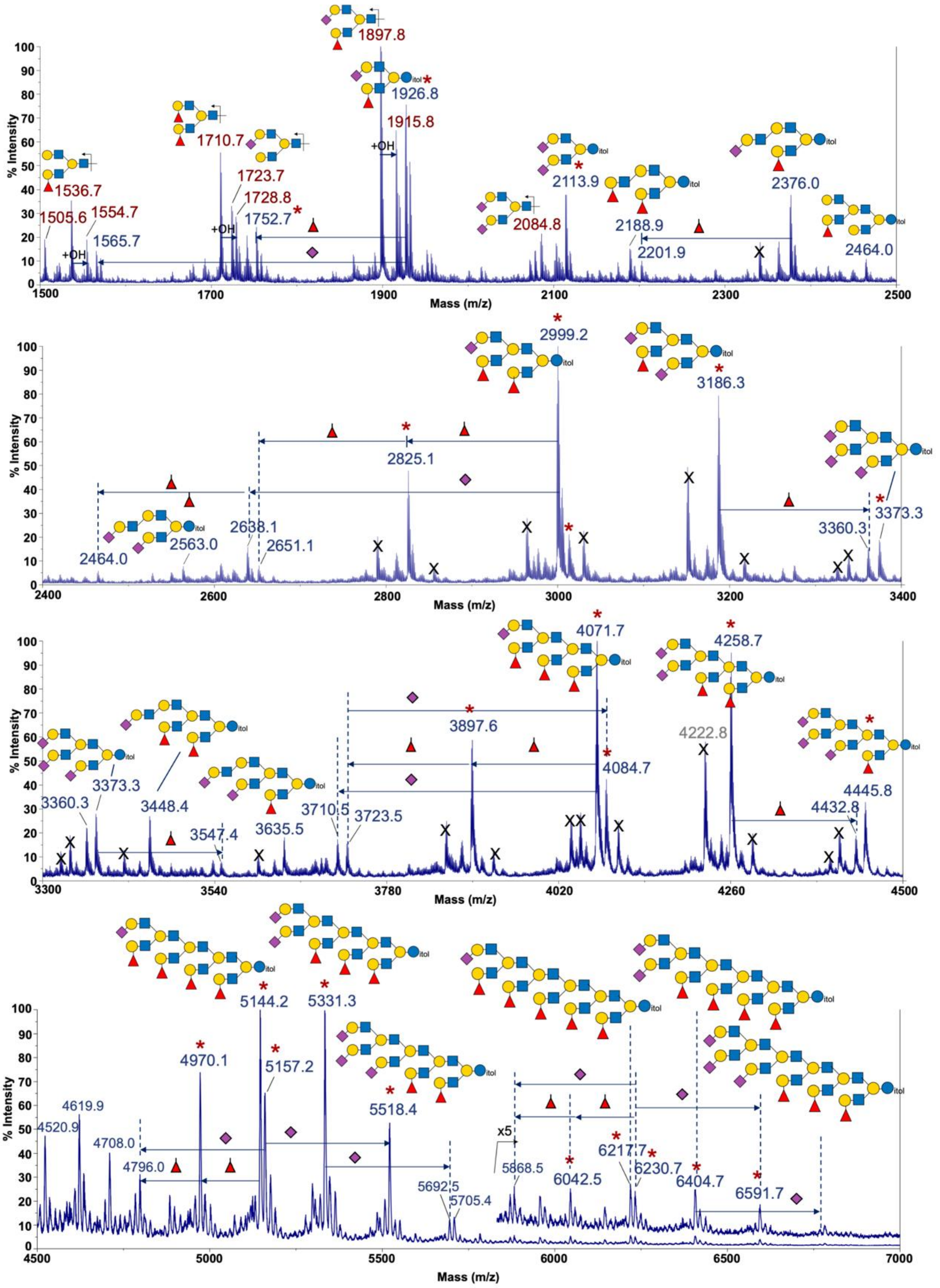

**Figure S3. Giant milk oligosaccharides reside in seal milk.** For a representative acidic fraction of our seal milk (JC_231115MA5, seal B, day 7), we report a characteristic branching pattern that seems to repeat and forms structures up to 28 monosaccharides (suggested structure candidates shown) via MALDI-MS. The intact $[M+Na]^+$ molecular ions were annotated in blue, to be distinguished from in-source produced oxonium ions annotated in red. The smallest components detected were found to carry two LacNAcs on the reducing end lactose core, with additional $Fuc_1$, $Neu5Ac_1$, $Fuc_1Neu5Ac_1$, or $Neu5Ac_2$. $MS^2$ data from subsequent LC-MS/MS analysis confirmed their branched nature, showing that the two LacNAcs were mostly fucosylated or sialylated at the terminal Gal, as annotated here for the peaks at *m/z* 1926.8 and 2113.9. Further extension with one LacNAc was possible but the major peak series, marked by asterisks, strikingly increased by a $Fuc_1LacNAc_2$ unit, giving rise to a series of MO extended by a polyLacNAc chain of varying length and branched at each Gal position. Each of the terminal LacNAc units was capped with Fuc or Neu5Ac. An exemplary HCD-$MS^2$ spectrum afforded by a triply protonated version (*m/z* $993.5^{3+}$) of the major peak at *m/z* 2999.2 is shown in Fig S4A.

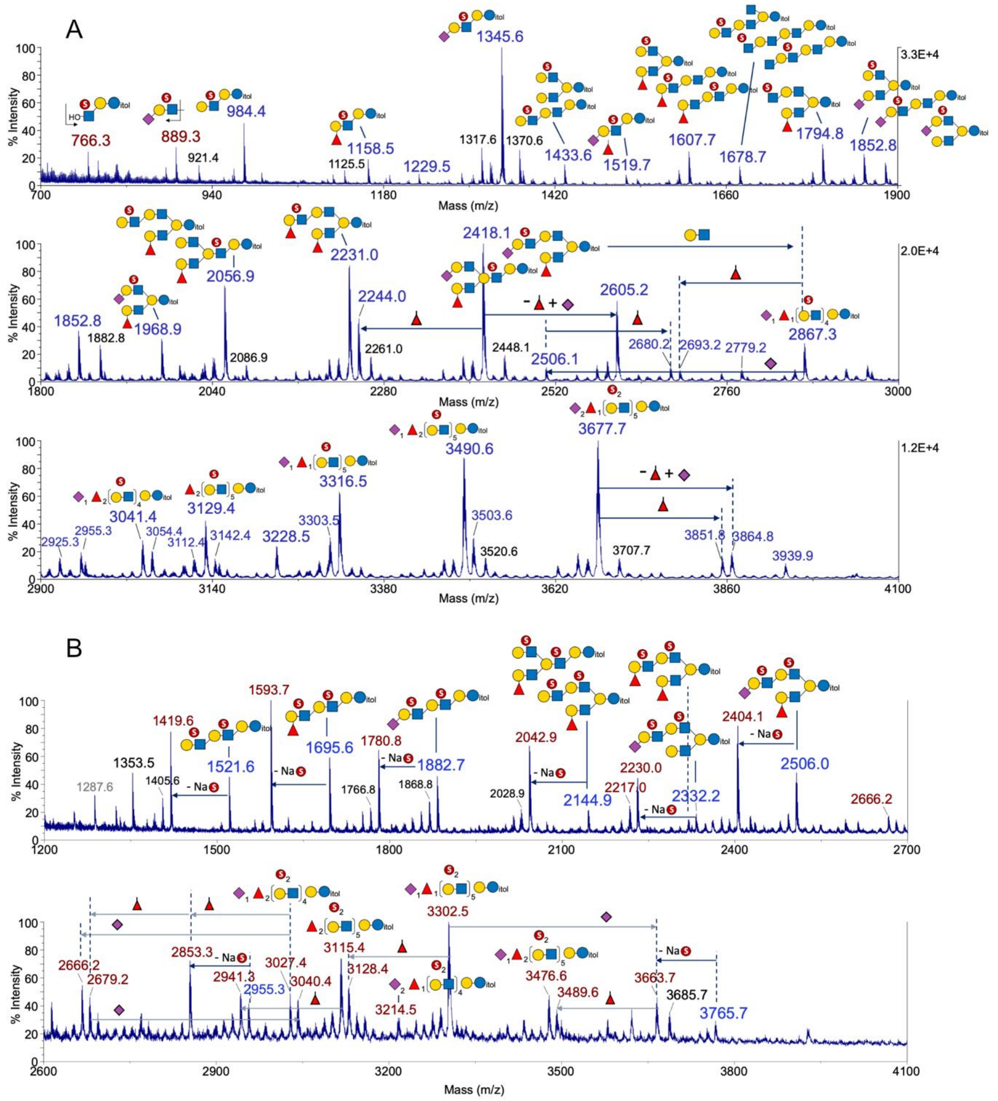

**Figure S4**. **MALDI-MS profiles of permethylated sulfated seal MO from the acidic fraction of sample JC_231115MA5**. The mono (a) and disulfated (b) MO were eluted from an Oasis MAX cartridge in two separate fractions away from the non-sulfated MO. Both were analyzed in negative mode. The monosulfated structures were detected as [M-H]$^-$, retaining the sulfate. The disulfated structures were detected as [M-2H+Na]$^-$, retaining both sulfates (annotated in blue), or as [M-SO$_3$-H]$^-$, after losing one sulfate and exposing a free OH (annotated in red). Unlike the non-sulfated MO, the smallest monosulfated components start with one fucosylated or sialylated LacNAc on the lactose core. This could then be extended by another LacNAc unit but, similar to the non-sulfated MO, the major peak series also increased by a Fuc$_1$LacNAc$_2$ unit, maintaining a similar branched pattern. Interestingly, the LC-MS$^2$ data clearly show that for structures with only one LacNAc (*m/z* 1158, 1345), the sulfate was carried on the GlcNAc. This GlcNAc-6-*O*-sulfation likely inhibited the branching of the preceding Gal. For structures with two LacNAcs (*m/z* 1433, 1519, 1607), the sulfate was carried either on the inner LacNAc of a linear diLacNAc, which inhibited the branching, or on the LacNAc of the 6-arm or a distal LacNAc of a diLAcNAc and thus not inhibiting the branching. The same sulfation pattern was similarly found in structures with three LacNAcs (*m/z* 2056, 2231, 2418), each of which could be resolved into two main isomeric forms by LC-MS$^2$ analysis (see Fig S4B for example MS$^2$ spectra). The deduced structures based on careful interpretation of the LC-MS$^2$ data acquired on signals below *m/z* 2500 are as annotated. Others not supported by productive or conclusive MS$^2$ results were annotated by glycan compositions only. Further MS$^2$ analyses of the disulfated structures likewise showed that sulfate was exclusively carried on the 6-position of GlcNAc and not found on the GlcNAc 3-linked to a 3,6-branched Gal. This apparent inhibitory effect of GlcNAc6S on the further 6-branching of Gal resulted in the major sulfated MO carrying 1, 3, and 5 LacNAc units, whereas the major nonsulfated MO carried 2, 4, 6, 8, and 10 LacNAc units instead.

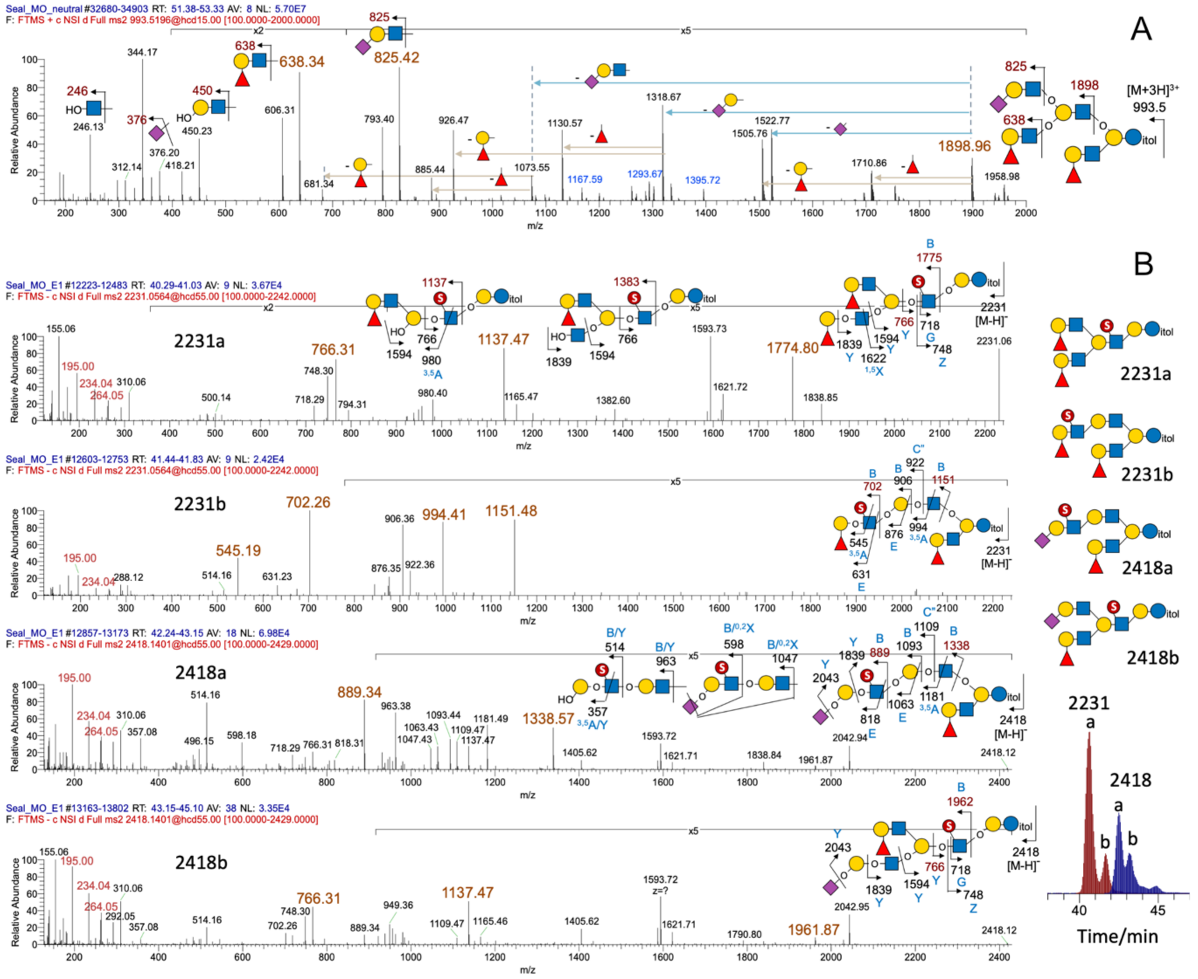

**Figure S5**. **Example HCD-MS$^2$ analyses of permethylated nonsulfated (a) and monosulfated (b) seal MO**. (a) Positive mode MS$^2$ on multiply protonated molecular ion such as the one shown here for *m/z* 993.5$^{3+}$ afforded mainly oxonium ions resulting from preferential cleavage at GlcNAc, often in combination with a series of neutral losses of nonreducing end Fuc, Fuc-Hex, and/or Neu5Ac residues. Importantly, the oxonium ions at *m/z* 638 and 825 identified terminal H1 and sialyl LacNAc units. These terminal fucosylated or sialylated LacNAc units could only be extended not by one but two LacNAc units to create the next oxonium ion at *m/z* 1898, thus establishing the branched pattern. Loss of a MeOH moiety from the oxonium ion at *m/z* 638 to create *m/z* 606 indicated a non-substituted OMe at C3 of GlcNAc, consistent with the H1, and not Lewis, structure and was further corroborated by the product dependent-MS$^3$ analyses of these MS$^2$ ions. (b) In negative mode, the low mass ions at *m/z* 195, 234, and 264 collectively identified an internal GlcNAc-6-*O*-sulfate, whereas the absence of *m/z* 153, 181, 253, 283 ruled out the alternative sulfate on terminal Gal. Liquid chromatography partially resolved each of the two structures (defined by *m/z* 2231 and 2418) into two distinct peaks (designated a and b, see the extracted ion chromatogram on the right panel). The main fragment ions were assigned as shown in the cartoon illustration, and the ion types labelled accordingly. These included the E, G, C" ions observed only in negative ion mode, in addition to the common B, Y, Z, $^{3,5}$A, $^{1,5}$X, $^{0,2}$X ions. The two isomeric forms were distinguished mainly by the ions at *m/z* 766 (accompanied by *m/z* 748, 718), which defined the presence of a sulfate on GlcNAc directly attached to the lactose core. This was further supported by the corresponding B ions at *m/z* 1775 and 1962 containing three LacNAc units. The alternative isomers with a linear diLacNAc would afford the B ions at *m/z* 1151 and 1338, respectively.

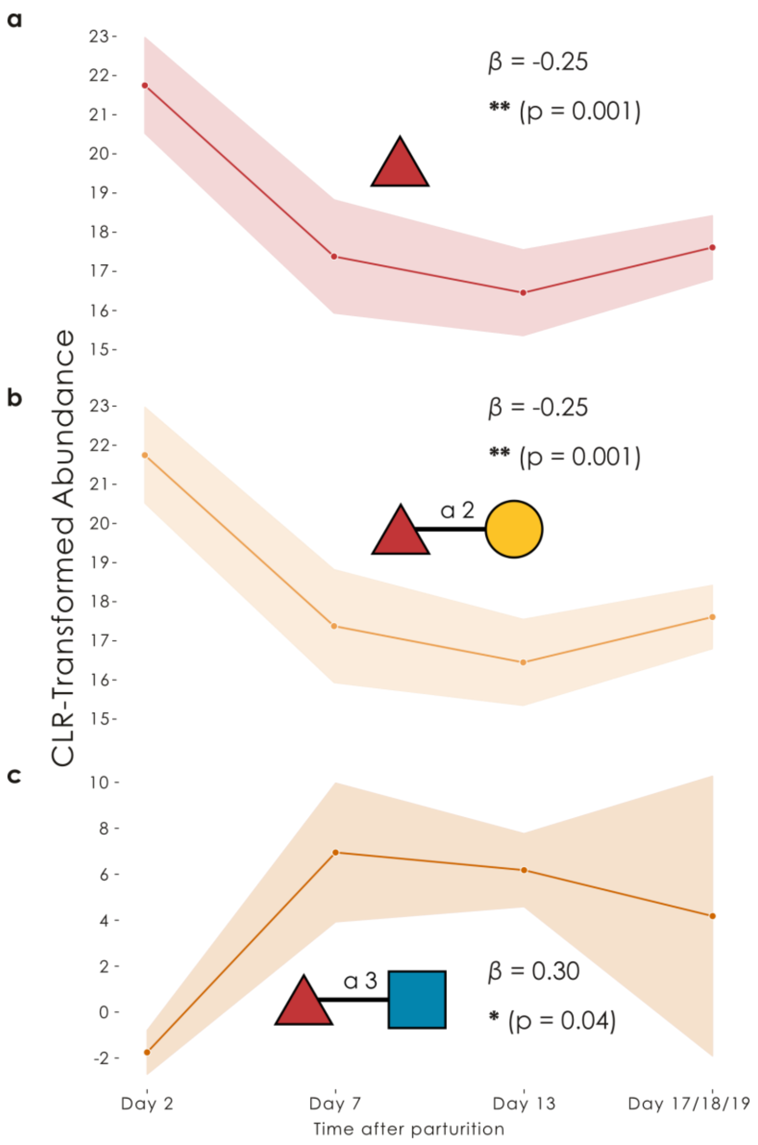

**Figure S6. Shifting fucose usage in seal milk during lactation. a-c)** For a dataset of five seals and four timepoints (N = 20), we used the *get_time_series* function of glycowork (v1.5) to analyze the expression of fucosylated glycans (a), Fucα1-2Gal-containing glycans (b), and Fucα1-3GlcNAc-containing glycans (c) throughout the lactation period. Shown are line plots of the CLR-transformed motif abundances, with a 95% confidence band, as well as the regression coefficient ($\beta$) and its significance, from fitting a degree 1 polynomial function to the time series. The fourth timepoint was collected at either day 17, 18, or 19 after giving birth, depending on the individual. *p<0.05, **p<0.01

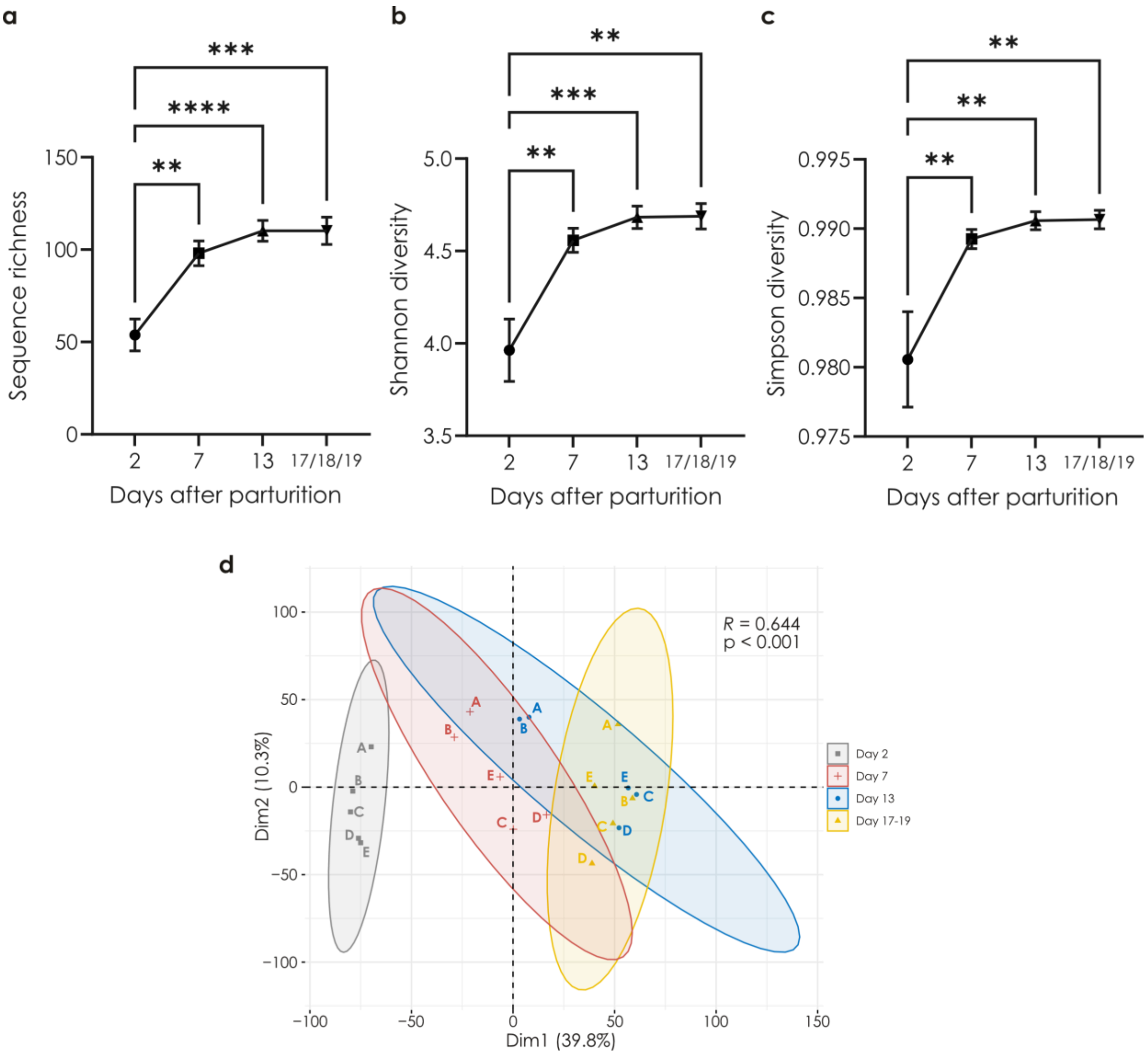

**Figure S7. The seal milk glycome is changing and becoming more diverse during lactation. a-c)** The seal milk glycome increases in diversity over time. Using the CLR-transformed seal milk glycome dataset, for each timepoint, we calculated sequence richness (a), Shannon diversity (b), and Simpson diversity (c) as three different alpha diversity measurements. Statistical comparisons were done via an ANOVA, followed by Tukey's Honestly Significant Difference post-hoc tests and a Benjamini-Hochberg correction for multiple testing. Data are shown as line plots with dots representing mean values and error bars their standard deviation. **d)** Not only glycome diversity but glycan content changes during lactation. Shown are the first two components of a PCA of the beta diversities of our CLR-transformed seal milk abundances. A directional trajectory of milk glycome change was confirmed via an ANOSIM analysis ($R = 0.644$, $p < 0.001$). Timepoints are color-coded and the percent variance explained of each component is indicated on each axis. For all panels (a-d), we used the *get_biodiversity* function from glycowork (v1.5). **p<0.01, ***p<0.001, ****p<0.0001

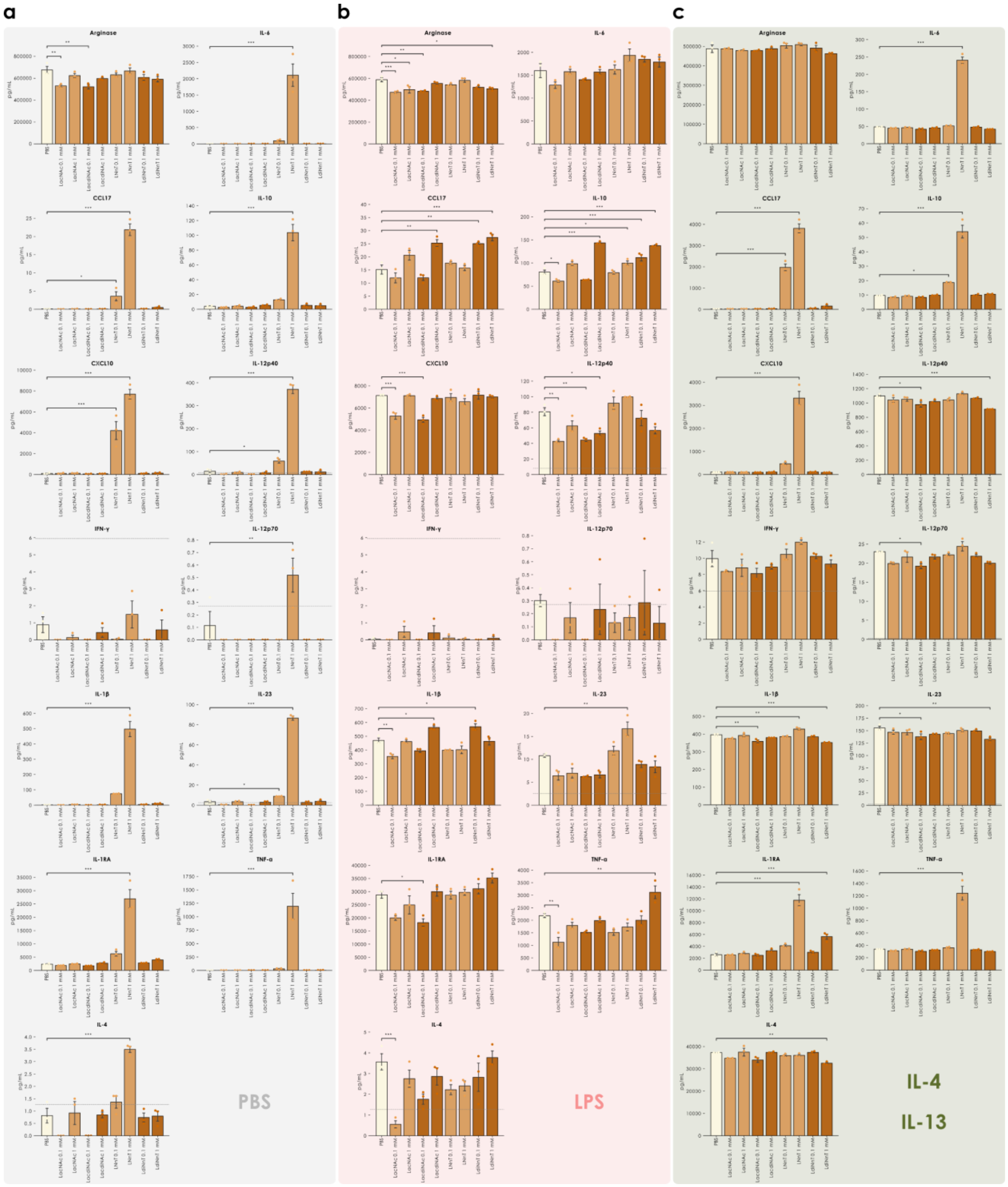

**Figure S8. Effects of LacNAc and LacdiNAc on human macrophages. a-c)** Quantification of cytokine concentrations of arginase, CCL17, CXCL10, IFN-γ, IL-1β, IL-1RA, IL-4, IL-6, IL-10, IL-12p40, IL-12p70, IL-23 and TNF-α from the culture supernatant of naïve (a), M1-polarized (b), and M2-polarized (c) macrophages in the absence or presence of varying concentrations LacNAc, LacdiNAc, LNnT, and LdiNnT. The dashed line indicates the limit of detection as determined by the standard curve of each analyte. Significant differences were established via a one-way ANOVA with post-hoc Tukey's HSD tests. ***, $p < 0.001$; **, $p < 0.01$; *, $p < 0.05$.

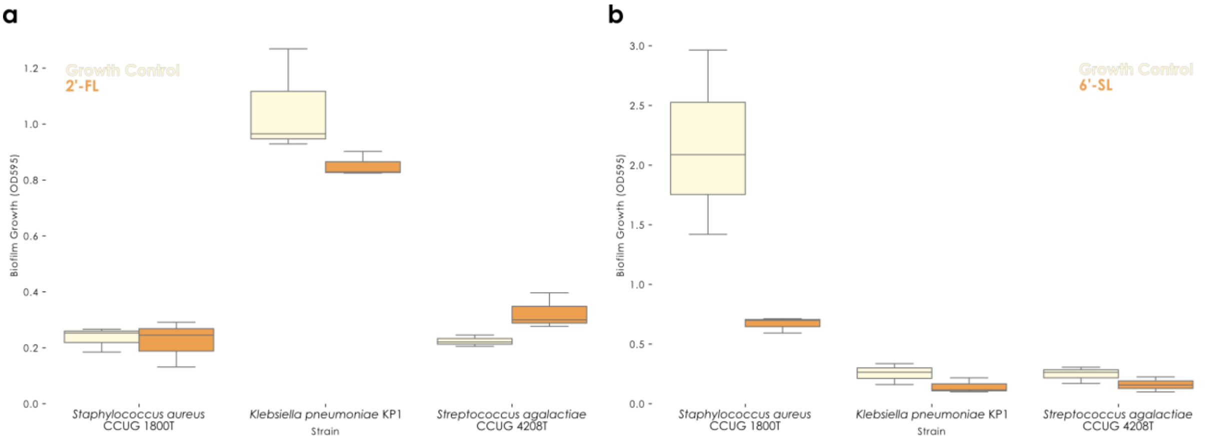

**Figure S9. Anti-biofilm properties of well-known milk oligosaccharides. a-b)** For three bacterial strains, we assessed their biofilm formation (measured as OD595 absorption, adjusted for blank medium measurements) with and without the addition of various milk oligosaccharides. “Growth control” refers to bacteria grown without milk oligosaccharides, while other conditions included 2’-FL (2’-fucosyllactose, 1 mg/mL; a) and 6’-SL (6’-sialyllactose, 1 mg/mL; b). The box plots depict mean values, with box edges indicating quartiles and whiskers indicating the remaining data distribution.

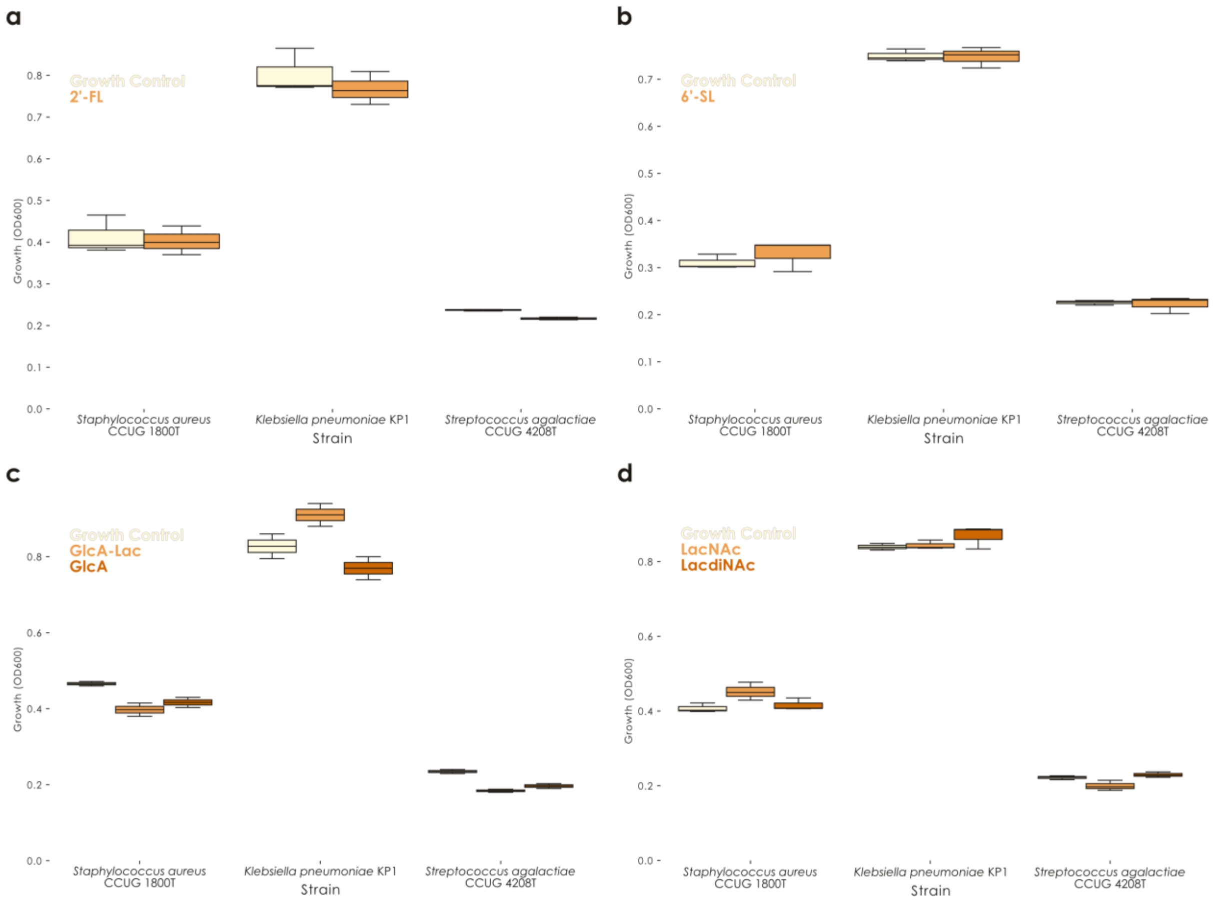

**Figure S10. The addition of milk oligosaccharides does not impact bacterial growth. a-d)** For three bacterial strains, we assessed their growth (measured as OD600 absorption, adjusted for blank medium measurements) with and without the addition of various milk oligosaccharides. "Growth control" refers to bacteria grown without milk oligosaccharides, while other conditions included 2'-FL (2'-fucosyllactose, 1 mg/mL; a), 6'-SL (6'-sialyllactose, 1 mg/mL; b), GlcA-Lac (glucoronyllactose, 1 mg/mL; c) and GlcA (glucuronic acid, 1 mg/mL; c), and LacNAc (*N*-acetyllactosamine, 1 mg/mL; d) and LacdiNAc (*N*,*N*-diacetyllactosamine, 1 mg/mL; d). The box plots depict mean values, with box edges indicating quartiles and whiskers indicating the remaining data distribution.

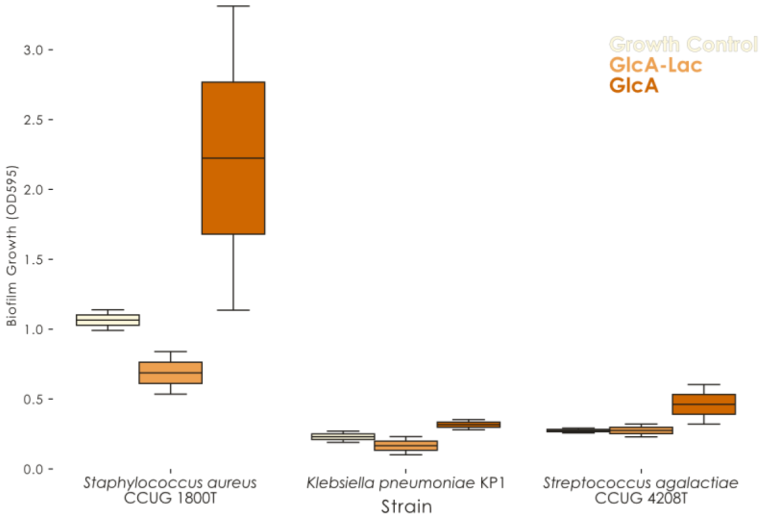

**Figure S11. Glucuronic acid-containing milk oligosaccharide show anti-biofilm potential.** For three bacterial strains, we assessed their biofilm formation (measured as OD595 absorption, adjusted for blank medium measurements) with and without the addition of various milk oligosaccharides. “Growth control” refers to bacteria grown without milk oligosaccharides, while other conditions included GlcA-Lac (glucoronyllactose, 1 mg/mL) and GlcA (glucuronic acid, 1 mg/mL). The box plots depict mean values, with box edges indicating quartiles and whiskers indicating the remaining data distribution.

### Synthetic protocols & physical data

**General methods.** Reactions were performed using commercial reagents as supplied unless stated otherwise. Anhydrous solvents were either commercially acquired or dried using standard methods and stored over molecular sieves. Thin layer chromatography was performed on silica-coated glass plates (TLC Silica Gel 60 $F_{254}$, Merck) employing UV light (254 nm), charring with 5% $H_2SO_4$ in ethanol, and/or staining with a ceric ammonium molybdate solution for visualization. Organic solutions were concentrated and/or evaporated to dry under vacuum in a water bath (<40 °C). Molecular sieves were dried at 400 °C under vacuum for 30 minutes prior to use. Amberlite IR-120H resin was washed extensively with MeOH and dried under vacuum prior to use. Medium-pressure liquid chromatography (MPLC) was performed using CombiFlash Companion instruments equipped with either RediSep normal-phase flash columns or self-packed reverse-phase C18 columns, and solvent gradients refer to sloped gradients with concentrations reported as % *v*/*v*. $^{1}H$ and $^{13}C$ NMR spectra were recorded on a Bruker Avance DMX-500 spectrometer at 500 and 125.8 MHz, respectively. Chemical shifts are expressed in ppm and referenced to either $Si(CH_3)_4$ ($\delta_H$ 0.00), residual $CHCl_3$ ($\delta_H$ 7.26 and $\delta_C$ 77.16, $CDCl_3$), residual $CHD_2OD$ ($\delta_H$ 3.31 and $\delta_C$ 49.15, $CD_3OD$), residual $CHD_2CN$ ($\delta_H$ 1.94 and $\delta_C$ 118.69, $CD_3CN$), or an acetone internal standard ($\delta_H$ 2.22 and $\delta_C$ 30.89, $D_2O$) [1]. Spectral assignments were achieved with the assistance of 2D gCOSY, 2D gTOCSY, 2D gHSQC, and 2D gHMBC. Overlapped signals in $^{1}H$ NMR spectra are reported in order of decreasing chemical shift. Low resolution electron-spray ionization mass spectrometry (ESI-MS) was performed using a Waters Micromass ZQ. High resolution mass spectrometry (HRMS) was performed using an Agilent 1100 LC equipped with a photodiode array detector and a Micromass QTOF I equipped with a 4 GHz digital-time converter. Optical rotations were measured with a Jasco P-2000 polarimeter at 20 °C. HPLC analysis was performed using an Agilent 1100 LC equipped with an EC 250/4 Nucleodur (5 μm, 4x250 mm) C18 column and detection via an evaporative light scattering detector (ELSD).

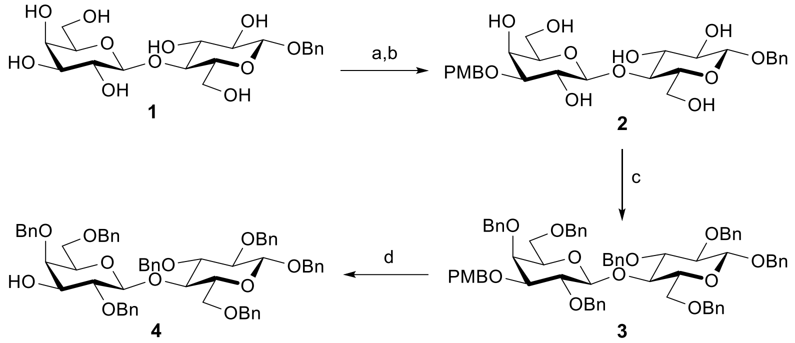

(a) $Bu_2SnO$, MeOH, reflux; (b) PMBCl, $NBu_4I$, THF, 72% (two steps); (c) BnBr, NaH, DMF, 0 °C → rt, 12 h, 92%; (d) TFA, $CH_2Cl_2$, 0 °C → rt, 20h, 78%

**Benzyl 3-*O*-(4-methoxybenzyl)-β-D-galactopyranosyl-(1→4)-β-D-glucopyranoside (2)**

Regioselective alkylation at the O-3 position was achieved with dibutyltin oxide using a method adapted from [2]. A suspension of benzyl β-D-lactoside (**1**, prepared according to literature [3]; 790 mg, 1.83 mmol) and $Bu_2SnO$ (591 mg, 2.37 mmol, 1.3 equiv.) in anhydrous MeOH (20 mL) was heated to reflux and stirred for 3 h until a homogeneous solution was formed. The solvent was removed in vacuo, and then the residue re-dissolved into anhydrous THF (20 mL). $NBu_4I$ (303 mg, 0.820 mmol, 0.45 equiv.) and 4-methoxybenzyl chloride (300 μL, 2.23 mmol, 1.2 equiv.) were added and the resulting mixture heated to reflux overnight. The mixture was concentrated and purified via MPLC on silica gel using using 0→5% MeOH – $CH_2Cl_2$ to afford the pure product as an amorphous colourless solid (**2**, 727 mg, 1.32 mmol, 72% yield). $R_f$ = 0.32 (1:9 MeOH : $CH_2Cl_2$). $[\alpha]_D^{20}$: -10 (*c* 1.0, MeOH). $^1$H-NMR ($CD_3OD$, 500 MHz): δH 7.43 – 7.39 (m, 2H, Ar), 7.38 – 7.30 (m, 4H, Ar), 7.29 – 7.24 (m, 1H, Ar), 6.90 – 6.86 (m, 2H, Ar), 4.92 (d, 1H, *J* = 11.8 Hz, PhC$H^a$H$^b$), 4.66 (d, 1H, *J* = 11.8 Hz, PhCH$^a$$H^b$), 4.69 (d, 1H, *J* = 11.3 Hz, $CH_3$OPhC$H^a$H$^b$), 4.58 (d, 1H, *J* = 11.3 Hz, $CH_3$OPhCH$^a$$H^b$), 4.39 (d, 1H, *J* = 8.0 Hz, Glc_H1), 4.38 (d, 1H, *J* = 7.8 Hz, Gal_H1), 3.98 (dd, 1H, *J* = 3.0, <1 Hz, Gal_H4), 3.94 (dd, 1H, *J* = 12.2, 2.5 Hz, Glc_H6$^a$), 3.87 (dd, 1H, *J* = 12.2, 4.3 Hz, Glc_H6$^b$), 3.80 – 3.75 (m, 1H, Gal_H6$^a$), 3.77 (s, 3H, $CH_3$O), 3.68 (dd, 1H, *J* = 11.5, 4.6 Hz, Gal_H6$^b$), 3.67 (dd, 1H, *J* = 9.6, 7.9 Hz, Gal_H2), 3.62 – 3.56 (m, 1H, Glc_H4), 3.55 – 3.49 (m, 2H, Glc_H3, Gal_H5), 3.40 (ddd, 1H, *J* = 9.6, 4.2, 2.4 Hz, Glc_H5), 3.36 (dd, 1H, *J* = 9.8, 3.2 Hz, Gal_H3), 3.34 (dd, 1H, *J* = 9.2, 7.8 Hz, Glc_H2). $^{13}$C-NMR ($CD_3OD$, 125 MHz): δC 160.94 (Ar), 139.14 (Ar), 131.80 (Ar), 130.93 (Ar), 129.44 (Ar), 129.32 (Ar), 128.86 (Ar), 114.77 (Ar), 105.13 (Gal_C1), 103.28 (Glc_C1),

81.8 (Gal_C3), 80.69 (Glc_C4), 77.06 (Gal_C5), 76.63 (Glc_C5), 76.54 (Glc_C3), 74.86 (Glc_C2), 72.36 ($CH_3OPhCH_2$), 71.93 (Gal_C2), 71.90 ($PhCH_2$), 67.12 (Gal_C4), 62.61 (Gal_C6), 62.00 (Glc_C6), 55.77 ($CH_3O$). LRMS *m/z* calc'd for $C_{27}H_{36}NaO_{12}$ $(M+Na)^+$: 575.21; found: 575.22.

**Benzyl 2,4,6-tri-*O*-benzyl-3-*O*-(4-methoxybenzyl)-β-D-galactopyranosyl-(1→4)-2,3,6-tri-*O*-benzyl-β-D-glucopyranoside (3)**

A mixture of **2** (612 mg, 1.11 mmol) and 60% sodium hydride (399 mg, 9.97 mmol, 9 equiv.) in dry DMF (18 mL) was stirred for 20 min at 0 °C, followed by the dropwise addition of benzyl bromide (1.30 mL, 11.0 mmol, 9.9 equiv.) (procedure adapted from [4]). The resulting suspension was stirred overnight at rt, and then quenched with methanol (2 mL) and concentrated under vacuum. The residue was diluted with ethyl acetate (30 mL) and washed with sat'd $NaHCO_3$ (aq) solution (20 mL), water (2 x 20 mL), dried with $Na_2SO_4$, filtered, and concentrated to dry. Purification of the residue via MPLC on silica gel using 0→20% ethyl acetate – petroleum ether yielded a colourless syrup (**3**; 1.11 g, 1.02 mmol, 92%). $R_f$ = 0.41 (1:3 ethyl acetate : hexanes). $[\alpha]_D^{20}$: +1.8 (*c* 1.0, $CHCl_3$). $^1$H-NMR ($CDCl_3$, 500 MHz): δH 7.43 – 7.13 (m, 37H, Ar), 6.89 – 6.87 (m, 2H, Ar), 5.06 (d, 1H, *J* = 10.7 Hz, $PhCH^aH^b$), 5.01 (d, 1H, *J* = 11.4 Hz, $PhCH^aH^b$), 4.99 (d, 1H, *J* = 12.0 Hz, $PhCH^aH^b$), 4.95 (d, 1H, *J* = 10.9 Hz, $PhCH^aH^b$), 4.85 (d, 1H, *J* = 11.2 Hz, $PhCH^aH^b$), 4.80 (d, 1H, *J* = 11.2 Hz, $PhCH^aH^b$), 4.79 (d, 1H, *J* = 10.9 Hz, $PhCH^aH^b$), 4.75 (d, 1H, *J* = 10.7 Hz, $PhCH^aH^b$), 4.70 (d, 1H, *J* = 12.0 Hz, $PhCH^aH^b$), 4.69 (d, 1H, *J* = 11.3 Hz, $CH_3OPhCH^aH^b$), 4.65 (d, 1H, *J* = 11.3 Hz, $CH_3OPhCH^aH^b$), 4.60 (d, 1H, *J* = 12.1 Hz, $PhCH^aH^b$), 4.59 (d, 1H, *J* = 11.4 Hz, $PhCH^aH^b$), 4.53 (d, 1H, *J* = 7.7 Hz, Glc_H1), 4.49 (d, 1H, *J* = 7.7 Hz, Gal_H1), 4.46 (d, 1H, *J* = 12.1 Hz, $PhCH^aH^b$), 4.38 (d, 1H, *J* = 11.8 Hz, $PhCH^aH^b$), 4.28 (d, 1H, *J* = 11.8 Hz, $PhCH^aH^b$), 4.00 (dd, 1H, *J* = 9.8, 8.7 Hz, Glc_H4), 3.93 (dd, 1H, *J* = 2.8, <1 Hz, Gal_H4), 3.86 (dd, 1H, *J* = 10.9, 4.4 Hz, Glc_H6$^a$), 3.83 (s, 3H, $CH_3O$), 3.80 – 3.75 (m, 2H, Glc_H6$^b$, Gal_H2), 3.62 – 3.55 (m, 2H, Glc_H3, Gal_H6$^a$), 3.52 (dd, 1H, *J* = 9.2, 7.7 Hz, Glc_H2), 3.44 (dd, 1H, *J* = 9.9, 3.0 Hz, Gal_H3), 3.42 – 3.37 (m, 3H, Gal_H5, Glc_H5, Gal_H6$^b$). $^{13}$C-NMR ($CDCl_3$, 125 MHz): δC 159.25 (Ar), 139.29 (Ar), 139.23 (Ar), 139.02 (Ar), 138.78 (Ar), 138.57 (Ar), 138.25 (Ar), 137.72 (Ar), 130.76 (Ar), 129.20 (Ar), 128.49 (Ar), 128.48 (Ar), 128.38 (Ar), 128.33 (Ar), 128.27 (Ar), 128.21 (Ar), 128.18 (Ar), 128.08 (Ar), 128.00 (Ar), 127.95 (Ar), 127.91 (Ar), 127.86 (Ar),

127.79 (Ar), 127.68 (Ar), 127.63 (Ar), 127.55 (Ar), 127.51 (Ar), 127.44 (Ar), 127.18 (Ar), 113.87 (Ar), 102.94 (Gal_C1), 102.65 (Glc_C1), 83.15 (Glc_C3), 82.47 (Gal_C3), 81.93 (Glc_C2), 81.10 (Gal_C2), 76.91 (Glc_C4), 75.50 (Ph*C*$H_2$), 75.35 (Gal_C5), 75.33 (Ph*C*$H_2$), 75.16 (Ph*C*$H_2$), 74.81 (Ph*C*$H_2$), 73.78 (Gal_C4), 73.52 (Ph*C*$H_2$), 73.22 (Ph*C*$H_2$), 73.14 (Glc_C5), 72.41 ($CH_3$OPh*C*$H_2$), 71.08 (Ph*C*$H_2$), 68.46 (Glc_C6), 68.26 (Gal_C6), 55.38 (*C*$H_3$O). LRMS *m/z* calc'd for $C_{69}H_{72}NaO_{12}$ $(M+Na)^+$: 1115.49; found: 1115.40.

**Benzyl 2,4,6-tri-*O*-benzyl-β-D-galactopyranosyl-(1→4)-2,3,6-tri-*O*-benzyl-β-D-glucopyranoside (4)**

To a solution of compound **4** (770 mg, 0.704 mmol) in $CH_2Cl_2$ (8 mL) was added trifluoroacetic acid (230 μL, 3.51 mmol, 5.0 equiv). The mixture was stirred at rt for 20 h and then diluted with $CH_2Cl_2$ (20 mL) and washed with sat'd $NaHCO_3$ (aq) solution (2 x 15 mL), water (2 x 15 mL), dried with $Na_2SO_4$, filtered, and concentrated to dry. Purification of the residue via MPLC on silica gel using 0→40% ethyl acetate – toluene yielded **4** as a colorless oil (537 mg, 0.552 mmol, 78%). $R_f$ = 0.36 (1:3 ethyl acetate : hexanes). $[\alpha]_D^{20}$: -5.6 (*c* 1.0, $CHCl_3$). $^1$H-NMR ($CDCl_3$, 500 MHz): δH 7.34 – 7.08 (m, 35H, Ar), 4.97 (d, 1H, *J* = 10.7 Hz, PhC$H^aH^b$), 4.91 (d, 1H, *J* = 12.1 Hz, PhC$H^aH^b$), 4.87 (d, 1H, *J* = 10.9 Hz, PhC$H^aH^b$), 4.76 (d, 1H, *J* = 11.5 Hz, PhC$H^aH^b$), 4.72 (d, 1H, *J* = 11.6 Hz, PhC$H^aH^b$), 4.70 (d, 1H, *J* = 10.9 Hz, PhC$H^aH^b$), 4.69 (d, 1H, *J* = 10.7 Hz, PhC$H^aH^b$), 4.63 (d, 1H, *J* = 11.5 Hz, PhC$H^aH^b$), 4.62 (d, 1H, *J* = 12.1 Hz, PhC$H^aH^b$), 4.58 (d, 1H, *J* = 11.6 Hz, PhC$H^aH^b$), 4.57 (d, 1H, *J* = 12.1 Hz, PhC$H^aH^b$), 4.45 (d, 1H, *J* = 7.7 Hz, Glc_H1), 4.41 (d, 1H, *J* = 12.1 Hz, PhC$H^aH^b$), 4.38 (d, 1H, *J* = 7.0 Hz, Gal_H1), 4.34 (d, 1H, *J* = 11.8 Hz, PhC$H^aH^b$), 4.23 (d, 1H, *J* = 11.8 Hz, PhC$H^aH^b$), 3.95 (dd, 1H, *J* = 9.9, 8.8 Hz, Glc_H4), 3.80 (dd, 1H, *J* = 2.7, <1 Hz, Gal_H4), 3.77 (dd, 1H, *J* = 10.9, 4.2 Hz, Glc_$H6^a$), 3.72 (dd, 1H, *J* = 10.9, 1.9 Hz, Glc_$H6^b$), 3.55 – 3.49 (m, 2H, Glc_H3, Gal_$H6^a$), 3.50 – 3.42 (m, 3H, Glc_H2, Gal_H2, Gal_H3), 3.41 – 3.34 (m, 2H, Gal_H5, Gal_$H6^b$), 3.33 (ddd, 1H, *J* = 9.9, 4.0, 1.7 Hz, Glc_H5), 2.04 (bs, 1H, OH). $^{13}$C-NMR ($CDCl_3$, 125 MHz): δC 139.24 (Ar), 138.86 (Ar), 138.76 (Ar), 138.59 (Ar), 138.42 (Ar), 138.17 (Ar), 137.68 (Ar), 128.58 (Ar), 128.51 (Ar), 128.50 (Ar), 128.44 (Ar), 128.40 (Ar), 128.24 (Ar), 128.22 (Ar), 128.13 (Ar), 128.06 (Ar), 128.03 (Ar), 127.88 (Ar), 127.84 (Ar), 127.82 (Ar), 127.75 (Ar), 127.70 (Ar), 127.67 (Ar), 127.65 (Ar), 127.30 (Ar), 102.80 (Gal_C1), 102.63 (Glc_C1), 83.04 (Glc_C3),

81.92, 80.77 (Gal_C2 and Glc_C2), 76.84 (Glc_C4), 76.05 (Gal_C4), 75.51 (Ph*C*$H_2$), 75.33 (Glc_C5), 75.24 (Ph*C*$H_2$), 75.17 (Ph*C*$H_2$), 75.10 (Ph*C*$H_2$), 74.24 (Gal_C3), 73.51 (Ph*C*$H_2$), 73.35 (Gal_C5), 73.31 (Ph*C*$H_2$), 71.09 (Ph*C*$H_2$), 68.42 (Glc_C6), 68.09 (Gal_C6). LRMS *m/z* calc'd for $C_{61}H_{64}NaO_{11}$ $(M+Na)^+$: 995.43; found: 995.21.

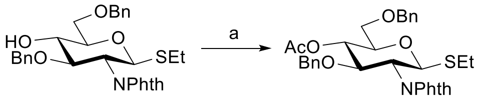

**5** **6**

(a) $Ac_2O$, $C_5H_5N$, 0 °C → rt, 2 h, 98%

**Ethyl 4-*O*-acetyl-3,6-di-*O*-benzyl-2-deoxy-2-phthalimido-1-thio-β-D-glucopyranoside (6)**

The starting material (**5**, prepared according to literature [4]; 305 mg, 0.572 mmol) was dissolved in dry pyridine (2.5 mL). The solution was cooled to 0 °C and $Ac_2O$ (2.0 mL) added, and then the reaction mixture was stirred at rt for 2 h. The reaction was quenched with ice (7 g) and the mixture was diluted with $CH_2Cl_2$ (20 mL). The organic phase was washed with 10% HCl (aq) solution (2 x 15 mL), water (2 x 15 mL), dried with $Na_2SO_4$, filtered, and concentrated to dry. Purification of the residue via MPLC on silica gel using 0→25% ethyl acetate – toluene yielded **6** as a colorless syrup (322 mg, 0.559 mmol, 98%). $R_f$ = 0.74 (1:4 ethyl acetate : toluene). $[\alpha]_D^{20}$: +68 (*c* 1.0, $CDCl_3$). $^1$H-NMR ($CDCl_3$, 500 MHz): δH 7.83 – 7.76 (m, 1H, Ar), 7.73 – 7.65 (m, 3H, Ar), 7.37 – 7.27 (m, 5H, Ar), 7.03 – 6.98 (m, 2H, Ar), 6.97 – 6.88 (m, 3H, Ar), 5.29 (d, 1H, *J* = 10.4 Hz, H-1), 5.15 (dd, 1H, *J* = 10.1, 8.9 Hz, H-4), 4.61 (d, 1H, *J* = 12.0 Hz, $PhCH^aH^b$ (*H*^a)), 4.57 (d, 1H, *J* = 12.2 Hz, $PhC\mathit{H}^aH^b$), 4.54 (d, 1H, *J* = 12.3 Hz, $PhCH^a\mathit{H}^b$), 4.48 (dd, 1H, *J* = 10.3, 8.9 Hz, H-3), 4.36 – 4.30 (m, 1H, H-2), 4.34 (d, 1H, *J* = 12.0 Hz, $PhCH^a\mathit{H}^b$), 3.79 (ddd, 1H, *J* = 10.1, 5.0, 4.0 Hz, H-5), 3.64 (dd, 1H, *J* = 10.8, 5.2 Hz, H-$6^a$), 3.61 (dd, 1H, *J* = 10.9, 4.1 Hz, H-$6^b$), 2.70 (dq, 1H, *J* = 12.6, 7.5 Hz, $SC\mathit{H}^aH^bCH_3$), 2.62 (dq, 1H, *J* = 12.5, 7.5 Hz, $SCH^a\mathit{H}^bCH_3$), 1.95 (s, 3H, OAc), 1.19 (dd, 3H, *J* = 7.4, 7.4 Hz, $SCH_2C\mathit{H}_3$). $^{13}$C-NMR ($CDCl_3$, 125 MHz): δC 169.75 (C=O), 168.23 (C=O), 167.32 (C=O), 138.03 (Ar), 137.74 (Ar), 134.08 (Ar), 133.95 (Ar), 131.67 (Ar), 128.45 (Ar), 128.22 (Ar), 127.94 (Ar), 127.89 (Ar), 127.77 (Ar), 127.56 (Ar), 123.69 (Ar), 123.38 (Ar), 81.26 (C-1), 78.08 (C-3), 77.78 (C-5), 74.17 (Ph*C*$H_2$), 73.67 (Ph*C*$H_2$), 72.65 (C-4), 69.96 (C-6), 54.70 (C-2), 24.13 (S*C*$H_2CH_3$), 20.99 (OAc),

15.03 (SCH$_2$*C*H$_3$). LRMS *m/z* calc'd for $C_{32}H_{33}NNaO_7S$ (M+Na)$^+$: 575.20; found: 575.14.

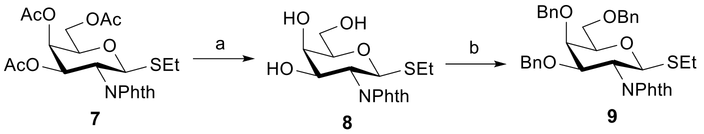

(a) NaOMe, MeOH, 0 °C, 1 h, 84 %; (b) BnBr, NaH, $Bu_4NBr$, DMF, 0 °C, 2 h, 87%.

**Ethyl 2-deoxy-2-phthalimido-1-thio-α-D-galactopyranoside (8)**

The starting material (**7**, prepared according to literature [5]; 500 mg, 1.14 mmol) was dissolved in anhydrous MeOH-$CH_2Cl_2$ 3:1 (4 mL), and then NaOMe solution was added drop-wise (0.5 M NaOMe in MeOH; to pH 10). After 1 h at 0 °C, the reaction mixture was neutralized with acidic resin (Amberlite IR-120H; to pH 6), filtered, and then evaporated to dry. The solid residue was crystallized from MeOH-ethyl acetate 1:9 to afford the pure product as a white solid (**8**; 341 mg, 0.965 mmol, 84% yield). $R_f$ = 0.56 (3:7 acetone : ethyl acetate). $[\alpha]_D^{20}$: +28 (*c* 1.0, MeOH). $^1$H-NMR (CD$_3$OD, 500 MHz): δH 7.91 – 7.80 (m, 4H, Ar), 5.27 (m/high-order coupling, 1H, H-1), 4.57 (bs, 1H, OH), 4.48 – 4.42 (m, 2H, H-3, H-2), 4.00 (d, 1H, *J* = <1 Hz, H-4), 3.81 (dd, 1H, *J* = 11.4, 6.9 Hz, H-6$^a$), 3.75 (dd, 1H, *J* = 11.4, 5.2 Hz, H-6$^b$), 3.68 (ddd, 1H, *J* = 6.9, 5.2, <1 Hz, H-5), 2.74 (dq, 1H, *J* = 12.7, 7.5 Hz, SC*H*$^a$H$^b$CH$_3$), 2.63 (dq, 1H, *J* = 12.7, 7.5 Hz, SCH$^a$*H*$^b$CH$_3$), 1.17 (dd, 3H, *J* = 7.5, 7.5 Hz, SCH$_2$C*H*$_3$). $^{13}$C-NMR (CD$_3$OD, 125 MHz): δC 170.15 (C=O), 169.83 (C=O), 135.72 (Ar), 135.66 (Ar), 133.41 (Ar), 133.14 (Ar), 124.51 (Ar), 124.18 (Ar), 83.03 (C-1), 81.31 (C-5), 70.55 (C-3), 70.29 (C-4), 62.88(C-6), 54.52 (C-2), 24.97 (S*C*H$_2$CH$_3$), 15.50 (SCH$_2$*C*H$_3$). LRMS *m/z* calc'd for $C_{16}H_{19}NNaO_6S$ (M+Na)$^+$: 376.08; found: 376.06.

**Ethyl 3,4,6-tri-*O*-benzyl-2-deoxy-2-phthalimido-1-thio-α-D-galactopyranoside (9)**

The starting material (**8**; 450 mg, 1.27 mmol), benzyl bromide (910 μL, 7.65 mmol, 6 equiv.), and tetrabutylammonium bromide (62 mg, 0.19 mmol, 0.15 equiv.) were dissolved into anhydrous DMF (7 mL). The mixture was cooled to 0 °C and NaH (60% oil dispersion; 0.23 g, 5.7 mmol, 4.5 equiv.) was added portion-wise and the mixture stirred for 1 h. The reaction was quenched via the drop-wise addition of $H_2O$ and the mixture diluted with ethyl acetate (10 mL) and saturated NaCl (aq) solution (10 mL).

The aqueous layer was removed and then re-extracted with ethyl acetate (10 mL). The combined organic phases were dried over $Na_2SO4$, filtered, and then evaporated to dry. The crude material was purified via MPLC using 0→20% ethyl acetate – hexanes to afford the pure product as a colourless syrup (**9**, 689 mg, 1.27 mmol, 87% yield). $R_f$ = 0.71 (1:4 ethyl acetate : toluene). $[\alpha]_D^{20}$: +39 (*c* 1.0, MeOH). $^1$H-NMR ($CD_3CN$, 500 MHz): δH 7.86 – 7.78 (m, 3H, Ar), 7.74 – 7.68 (m, 1H, Ar), 7.40 – 7.28 (m, 10H, Ar), 7.10 – 7.96 (m, 5H, Ar), 5.27 (d, 1H, *J* = 10.4 Hz, H-1), 4.87 (d, 1H, *J* = 11.2 Hz, PhC$H^a$$H^b$), 4.64 (d, 1H, *J* = 12.1 Hz, PhC$H^a$$H^b$), 4.60 (d, 1H, *J* = 11.2 Hz, PhC$H^a$$H^b$), 4.55 (d, 1H, *J* = 11.9 Hz, PhC$H^a$$H^b$), 4.50 (d, 1H, *J* = 11.9 Hz, PhC$H^a$$H^b$), 4.45 (dd, 1H, *J* = 10.4, 10.6 Hz, H-2), 4.36 (dd, 1H, *J* = 10.6, 2.9 Hz, H-3), 4.34 (d, 1H, *J* = 12.1 Hz, PhC$H^a$$H^b$), 4.17 (dd, 1H, *J* = 2.9, 1.1 Hz, H-4), 3.86 (ddd, 1H, *J* = 6.3, 6.3, 1.1 Hz, H-5), 3.65 – 3.61 (m, 2H, H-$6^a$, H-$6^b$), 2.62 (dq, 1H, *J* = 12.8, 7.4 Hz, SC$H^a$$H^b$$CH_3$), 2.54 (dq, 1H, *J* = 12.8, 7.4 Hz, SC$H^a$$H^b$$CH_3$), 1.11 (dd, 3H, *J* = 7.4, 7.4 Hz, $SCH_2CH_3$). $^{13}$C-NMR ($CD_3CN$, 125 MHz): δC 168.97 (C=O), 168.87 (C=O), 139.85 (Ar), 139.49 (Ar), 138.99 (Ar), 135.41 (Ar), 132.47 (Ar), 132.28 (Ar), 129.30 (Ar), 129.26 (Ar), 129.10 (Ar), 128.94 (Ar), 128.82 (Ar), 128.81 (Ar), 128.57 (Ar), 128.55 (Ar), 124.22 (Ar), 124.10 (Ar), 82.11 (C-1), 78.21 (C-5), 77.99 (C-3), 75.58 ($PhCH_2$), 74.09 (C-4), 73.79 ($PhCH_2$), 71.97 ($PhCH_2$), 70.00 (C-6), 52.75 (C-2), 24.68 ($SCH_2CH_3$), 15.51 ($SCH_2CH_3$). LRMS *m/z* calc'd for $C_{37}H_{37}NNaO_6S$ $(M+Na)^+$: 646.22; found: 646.28.

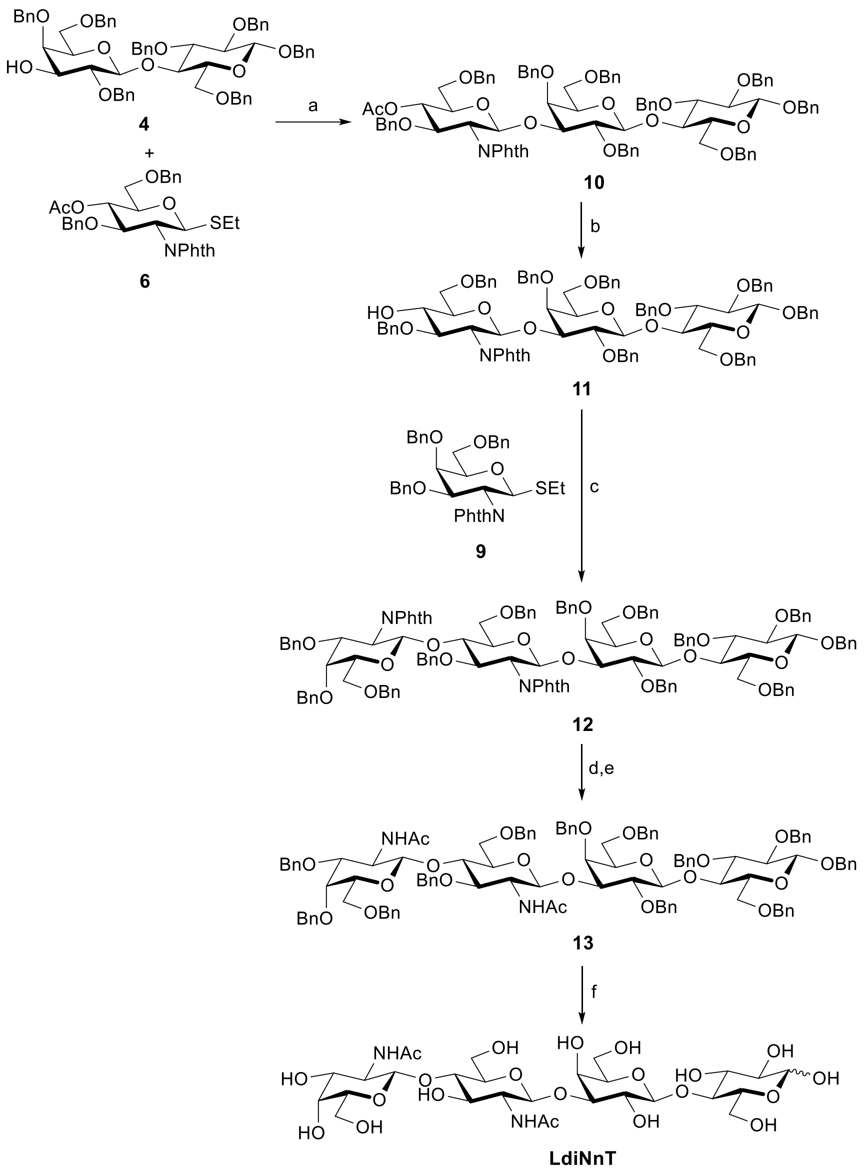

(a) NIS, TfOH, $CH_2Cl_2$, -10 °C → rt, 18 h, 96%; (b) NaOMe, MeOH, $CH_2Cl_2$, 2 h, 92%; (c) NIS, TfOH, $CH_2Cl_2$, -20 °C → rt, 4 h, 95%; (d) $N_2H_4{\cdot}H_2O$, EtOH, 80 °C, 72 h; (e) $Ac_2O$, $NaHCO_3$, MeOH, $H_2O$, rt, 3 h, 86% (2 steps); (f) $H_2$, Pd(C), 9:1 MeOH:$H_2O$, rt, 48 h, 92%.

**Benzyl 4-*O*-acetyl-3,6-di-*O*-benzyl-2-deoxy-2-phthalimido-β-D-glucopyranosyl-(1→3)-2,4,6-tri-*O*-benzyl-β-D-galactopyranosyl-(1→4)-2,3,6-tri-*O*-benzyl-β-D-glucopyranoside (10)**

The glycosyl donor (**6**; 306 mg, 0.532 mmol, 1.50 equiv.), glycosyl acceptor (**4**; 345 mg, 0.532 mmol), and crushed molecular sieves (3 Å, 300 mg) in anhydrous $CH_2Cl_2$ (6 mL) were left mixing at rt under Ar. After 30 min, the reaction flask was cooled to -10 °C, and then *N*-iodosuccinimide was added (286 mg, 1.27 mmol), followed by the drop-wise addition of TfOH (11.0 µL, 0.126 mmol). After 12 h with slow warming back to rt, the mixture was neutralized with triethylamine (to pH 7), further diluted with

$CH_2Cl_2$ (8 mL), filtered over Celite, and evaporated to dry. The crude mixture was then directly purified via MPLC on silica gel using 0→50% ethyl acetate – petroleum ether to afford the pure product as a colourless amorphous solid (**10**; 510 mg, 0.343 mmol, 96% yield). $R_f$ = 0.24 (1:3 ethyl acetate : hexanes). $[\alpha]_D^{20}$: +1.9 (*c* 1.0, $CDCl_3$). $^{1}$H-NMR ($CDCl_3$, 500 MHz): δH 7.66 – 7.35 (m, 2H, Ar), 7.34 – 7.02 (m, 40H, Ar), 6.94 – 6.91 (m, 2H, Ar), 6.89 – 6.85 (m, 2H, Ar), 6.84 – 6.79 (m, 3H, Ar), 5.35 (d, 1H, *J* = 8.3 Hz, GlcN_H1), 5.09 (dd, 1H, *J* = 10.1, 8.9 Hz, GlcN_H4), 4.97 (d, 1H, *J* = 11.4 Hz, PhC*H*$^a$H$^b$), 4.85 (d, 1H, *J* = 10.6 Hz, PhC*H*$^a$H$^b$), 4.81 (d, 1H, *J* = 12.0 Hz, PhC*H*$^a$H$^b$), 4.79 (d, 1H, *J* = 10.9 Hz, PhC*H*$^a$H$^b$), 4.64 (d, 1H, *J* = 10.9 Hz, PhCH$^a$*H*$^b$), 4.54 (d, 1H, *J* = 11.8 Hz, PhC*H*$^a$H$^b$), 4.53 – 4.48 (m, 4H, GlcN_H3, 3x PhCH$^a$*H*$^b$), 4.43 ('s', 2H, PhC*H*$^a$H$^b$, PhCH$^a$*H*$^b$), 4.42 (d, 1H, *J* = 12.1 Hz, PhC*H*$^a$H$^b$), 4.31 – 4.25 (m, 3H, GlcN_H2, PhC*H*$^a$H$^b$, PhCH$^a$*H*$^b$), 4.26 (d, 1H, *J* = 7.5 Hz, Glc_H1), 4.20 (d, 1H, *J* = 12.1 Hz, PhC*H*$^a$H$^b$), 4.18 – 4.12 (m, 3H, PhCH$^a$*H*$^b$, Gal_H1, PhCH$^a$*H*$^b$), 3.95 (dd, 1H, *J* = 3.0, <1 Hz, Gal_H4), 3.90 (d, 1H, *J* = 12.1 Hz, PhCH$^a$*H*$^b$), 3.82 – 3.75 (m, 2H, Glc_H4, GlcN_H5), 3.60 (dd, 1H, *J* = 10.6, 3.4 Hz, GlcN_H6$^a$), 3.56 (dd, 1H, *J* = 10.6, 6.0 Hz, GlcN_H6$^b$), 3.48 (dd, 1H, *J* = 9.8, 2.9 Hz, Gal_H3), 3.47 – 3.42 (m, 2H, Gal_H6$^a$, Glc_H6$^a$), 3.38 (dd, 1H, *J* = 9.8, 7.6 Hz, Gal_H2), 3.35 – 3.27 (m, 4H, Gal_H5, Glc_H2, Glc_H3, Gal_H6$^b$), 3.25 (dd, 1H, *J* = 10.9, 1.8 Hz, Glc_H6$^b$), 2.87 (ddd, 1H, *J* = 9.9, 3.9, 1.8 Hz, Glc_H5), 1.92 (s, 3H, OAc). $^{13}$C-NMR ($CDCl_3$, 125 MHz): δC 169.89 (C=O), 139.42 (Ar), 139.18 (Ar), 138.75 (Ar), 138.70 (Ar), 138.46 (Ar), 138.41 (Ar), 137.95 (Ar), 137.88 (Ar), 137.72 (Ar), 133.60 (Ar), 131.22 (Ar), 128.53 (Ar), 128.44 (Ar), 128.42 (Ar), 128.40 (Ar), 128.39 (Ar), 128.35 (Ar), 128.30 (Ar), 128.23 (Ar), 128.18 (Ar), 128.08 (Ar), 128.04 (Ar), 127.96 (Ar), 127.93 (Ar), 127.87 (Ar), 127.85 (Ar), 127.72 (Ar), 127.65 (Ar), 127.61 (Ar), 127.48 (Ar), 127.38 (Ar), 127.18 (Ar), 126.81 (Ar), 126.38 (Ar), 123.23 (Ar), 102.53 (Glc_C1), 102.49 (Gal_C1), 99.76 (GlcN_C1), 83.10 (Glc_C3), 82.01 (Gal_C3), 81.74 (Glc_C2), 78.83 (Gal_C2), 76.95 (GlcN_C3), 76.62 (Gal_C4), 76.07 (Glc_C4), 75.54 (Ph*C*$H_2$), 75.13 (Ph*C*$H_2$), 75.08 (Ph*C*$H_2$), 74.89 (Glc_C5), 74.13 (Ph*C*$H_2$), 74.00 (Ph*C*$H_2$), 73.91 (Ph*C*$H_2$), 73.43 (Ph*C*$H_2$), 73.39 (GlcN_C5), 73.26 (Gal_C5), 73.16 (Ph*C*$H_2$), 72.73 (GlcN_C4), 70.92 (Ph*C*$H_2$), 70.10 (GlcN_C6), 68.45 (Gal_C6), 67.71 (Glc_C6), 56.20 (GlcN_C2), 21.04 (OAc). LRMS *m/z* calc'd for $C_{91}H_{91}NNaO_{18}$ $(M+Na)^+$: 1508.61; found: 1508.37.

**Benzyl 3,6-di-*O*-benzyl-2-deoxy-2-phthalimido-β-D-glucopyranosyl-(1→3)-2,4,6-tri-*O*-benzyl-β-D-galactopyranosyl-(1→4)-2,3,6-tri-*O*-benzyl-β-D-glucopyranoside (11)**

The starting material (**10**; 380 mg, 0.256 mmol) was dissolved into anhydrous $CH_2Cl_2$ – MeOH 1:3 (4 mL), and then NaOMe solution was added drop-wise (1.0 M NaOMe in MeOH; to pH 10). After 2 h, the reaction mixture was neutralized with acidic resin (Amberlite IR-120H; to pH 6), filtered, and then evaporated to dry. The crude material was purified via MPLC on silica gel using 0→25% ethyl acetate – toluene to afford the pure product as a colourless syrup (**11**, 339 mg, 0.235 mmol, 92% yield). $R_f$ = 0.40 (1:4 ethyl acetate : toluene). $[\alpha]_D^{20}$: -10 (*c* 1.0, $CDCl_3$). $^1$H-NMR ($CDCl_3$, 500 MHz): δH 7.68 – 7.48 (m, 2H, Ar), 7.34 – 7.17 (m, 36H, Ar), 7.14 – 7.02 (m, 6H, Ar), 6.98 – 6.92 (m, 3H, Ar), 6.90 – 6.87 (m, 2H, Ar), 5.41 (d, 1H, *J* = 8.3 Hz, GlcN_H1), 5.00 (d, 1H, *J* = 11.4 Hz, PhC$H^a$$H^b$), 4.88 (d, 1H, *J* = 10.7 Hz, PhC$H^a$$H^b$), 4.86 (d, 1H, *J* = 12.1 Hz, PhC$H^a$$H^b$), 4.84 (d, 1H, *J* = 10.8 Hz, PhC$H^a$$H^b$), 4.73 (d, 1H, *J* = 12.2 Hz, PhC$H^a$$H^b$), 4.69 (d, 1H, *J* = 10.8 Hz, PhC$H^a$$H^b$), 4.60 (d, 1H, *J* = 11.8 Hz, PhC$H^a$$H^b$), 4.58 – 4.53 (m, 3H, 3x PhC$H^a$$H^b$), 4.52 – 4.49 (m, 2H, 2x PhC$H^a$$H^b$), 4.47 (d, 1H, *J* = 12.2 Hz, PhC$H^a$$H^b$), 4.37 (dd, 1H, *J* = 10.8, 8.4 Hz, GlcN_H3), 4.34 (d, 1H, *J* = 12.0 Hz, PhC$H^a$$H^b$), 4.32 (d, 1H, *J* = 7.4 Hz, Glc_H1), 4.28 – 4.22 (m, 4H, PhC$H^a$$H^b$, PhC$H^a$$H^b$, GlcN_H2, Gal_H1), 4.19 (d, 1H, *J* = 11.9 Hz, PhC$H^a$$H^b$), 4.00 (d, 1H, *J* = 11.9 Hz, PhC$H^a$$H^b$), 3.95 (dd, 1H, *J* = 2.9, <1 Hz, Gal_H4), 3.88 – 3.82 (m, 3H, GlcN_H4, Glc_H4, GlcN_H6$^a$), 3.80 (dd, 1H, *J* = 10.0, 4.8 Hz, GlcN_H6$^b$), 3.71 (ddd, 1H, *J* = 9.6, 4.8, 4.8 Hz, GlcN_H5), 3.54 (dd, 1H, *J* = 9.8, 2.9 Hz, Gal_H3), 3.53 – 3.47 (m, 2H, Glc_H6$^a$, Gal_H6$^a$), 3.44 (dd, 1H, *J* = 9.8, 7.6 Hz, Gal_H2), 3.40 – 3.31 (m, 5H, Gal_H5, Glc_H2, Gal_H6$^b$, Glc_H3, Glc_H6$^b$), 2.93 (ddd, 1H, *J* = 9.9, 3.9, 1.9 Hz, Glc_H5), 2.91 (bs, O*H*). $^{13}$C-NMR ($CDCl_3$, 125 MHz): δC 139.48 (Ar), 139.19 (Ar), 138.78 (Ar), 138.73 (Ar), 138.51 (Ar), 138.50 (Ar), 138.37 (Ar), 137.73 (Ar), 133.63 (Ar), 131.44 (Ar), 128.70 (Ar), 128.43 (Ar), 128.42 (Ar), 128.37 (Ar), 128.34 (Ar), 128.33 (Ar), 128.29 (Ar), 128.21 (Ar), 128.08 (Ar), 128.05 (Ar), 127.98 (Ar), 127.96 (Ar), 127.94 (Ar), 127.85 (Ar), 127.82 (Ar), 127.78 (Ar), 127.75 (Ar), 127.67 (Ar), 127.65 (Ar), 127.63 (Ar), 127.54 (Ar), 127.35 (Ar), 127.18 (Ar), 126.89 (Ar), 126.53 (Ar), 123.25 (Ar), 102.55 (2C, Glc_C1, Gal_C1), 99.80 (GlcN_C1), 83.11 (Glc_C3), 81.86 (Gal_C3), 81.79 (Glc_C2), 78.98 (Gal_C2), 78.80 (GlcN_C3), 76.70 (Gal_C4), 76.06 (Glc_C4), 75.55 (Ph$CH_2$), 75.15 (Ph$CH_2$), 75.09 (Ph$CH_2$), 74.93 (Glc_C5), 74.56

($PhCH_2$), 74.45 (GlcN_C4), 74.11 ($PhCH_2$), 74.05 ($PhCH_2$), 73.58 (GlcN_C5), 73.43 ($PhCH_2$), 73.21 (Gal_C5), 73.13 ($PhCH_2$), 71.03 (GlcN_C6), 70.96 ($PhCH_2$), 68.47 (Gal_C6), 67.83 (Glc_C6), 56.08 (GlcN_C2). LRMS *m/z* calc'd for $C_{89}H_{89}NNaO_{17}$ $(M+Na)^+$: 1466.60; found: 1467.29.

**Benzyl 3,4,6-tri-*O*-benzyl-2-deoxy-2-phthalimido-β-D-galactopyranosyl-(1→4)-3,6-di-*O*-benzyl-2-deoxy-2-phthalimido-β-D-glucopyranosyl-(1→3)-2,4,6-tri-*O*-benzyl-β-D-galactopyranosyl-(1→4)-2,3,6-tri-*O*-benzyl-β-D-glucopyranoside (12)**

The glycosyl donor (**9**; 101 mg, 0.162 mmol, 1.30 equiv.), glycosyl acceptor (**11**; 180 mg, 0.125 mmol), and crushed molecular sieves (3 Å, 250 mg) in anhydrous $CH_2Cl_2$ (5.0 mL) were left mixing at rt under Ar. After 30 minutes, the reaction flask was cooled to -20 °C, and then *N*-iodosuccinimide was added (50 mg, 0.22 mmol), followed by the drop-wise addition of TfOH (3.0 μL, 35 μmol). After 4 h with slow warming back to rt, no starting material was observed via TLC. The mixture was neutralized with triethylamine (to pH 7), further diluted with $CH_2Cl_2$ (8 mL), filtered over Celite, and evaporated to dry. The crude mixture was then directly purified via MPLC on silica gel using 0→45% ethyl acetate – petroleum ether to afford the pure product as a colourless syrup (238 mg, 0.119 mmol, 95% yield). $R_f$ = 0.60 (1:4 ethyl acetate : toluene). $[\alpha]_D^{20}$: +2.0 (*c* 1.0, $CDCl_3$). $^1H$-NMR ($CDCl_3$, 500 MHz): δH 7.94 – 9.90 (m, 1H, Ar), 7.77 – 7.72 (m, 1H, Ar), 7.71 – 7.67 (m, 2H, Ar), 7.64 – 7.44 (m, 2H, Ar), 7.33 – 7.15 (m, 42H, Ar), 7.14 – 7.00 (m, 13H, Ar), 6.94 – 6.91 (m, 2H, Ar), 6.86 – 6.76 (m, 5H, Ar), 5.31 (d, 1H, *J* = 8.4 Hz, GalN_H1), 5.24 (d, 1H, *J* = 8.3 Hz, GlcN_H1), 4.99 (d, 1H, *J* = 11.5 Hz, $PhCH^aH^b$), 4.94 (d, 1H, *J* = 11.6 Hz, $PhCH^aH^b$), 4.92 (d, 1H, *J* = 12.1 Hz, $PhCH^aH^b$), 4.87 – 4.80 (m, 3H, 3x $PhCH^aH^b$), 4.73 (dd, 1H, *J* = 11.1, 8.4 Hz GalN_H2), 4.67 (d, 1H, *J* = 10.9 Hz, $PhCH^aH^b$), 4.59 (d, 1H, *J* = 12.2 Hz, $PhCH^aH^b$), 4.55 – 4.50 (m, 3H, 3x $PhCH^aH^b$), 4.44 – 4.36 (m, 4H, 2x $PhCH^aH^b$, 2x $PhCH^aH^b$), 4.36 – 4.25 (m, 8H, GalN_H3, $PhCH^aH^b$, GlcN_H3, 2x $PhCH^aH^b$, Glc_H1, $PhCH^aH^b$, $PhCH^aH^b$), 4.24 – 4.16 (m, 3H, GlcN_H2, $PhCH^aH^b$, $PhCH^aH^b$), 4.15 (d, 1H, *J* = 7.0 Hz, Gal_H1), 4.14 – 4.09 (m, 2H, GlcN_H4, $PhCH^aH^b$), 4.03 (d, 1H, *J* = 2.8 Hz, GalN_H4), 3.96 (d, 1H, *J* = 12.0 Hz, $PhCH^aH^b$), 3.88 (dd, 1H, *J* = 2.5, <1 Hz, Gal_H4), 3.82 – 3.76 (m, 1H, Glc_H4), 3.60 – 3.56 (m, 1H, GalN_H5), 3.54 – 3.48 (m, 2H, GalN_H6$^a$, GlcN_H6$^a$), 3.47 – 3.35 (m, 7H, Glc_H6$^a$, GalN_H6$^b$, GlcN_H6$^b$, GlcN_H5, Gal_H6$^a$, Gal_H3, Gal_H2), 3.35 – 3.30 (m, 2H, Glc_H2, Glc_H3), 3.30 – 3.24 (m, 3H,

Gal_H5, Gal_H6[b], Glc_H6[b]), 2.90 (ddd, 1H, *J* = 9.9, 4.0, 1.9 Hz, Glc_H5). $^{13}$C-NMR ($CDCl_3$, 125 MHz): δC 169.01 (C=O), 167.94 (C=O), 139.52 (Ar), 139.18 (Ar), 139.03 (Ar), 138.99 (Ar), 138.78 (Ar), 138.66 (Ar), 138.52 (Ar), 138.46 (Ar), 138.45 (Ar), 138.20 (Ar), 137.91 (Ar), 137.73 (Ar), 134.16 (Ar), 133.97 (Ar), 133.50 (Ar), 132.02 (Ar), 131.83 (Ar), 131.39 (Ar), 128.53 (Ar), 128.42 (Ar), 128.38 (Ar), 128.36 (Ar), 128.30 (Ar), 128.28 (Ar), 128.18 (Ar), 128.13 (Ar), 128.02 (Ar), 127.99 (Ar), 127.94 (Ar), 127.89 (Ar), 127.88 (Ar), 127.86 (Ar), 127.84, 127.82 (Ar), 127.77 (Ar), 127.75 (Ar), 127.72 (Ar), 127.63 (Ar), 127.59 (Ar), 127.54 (Ar), 127.46 (Ar), 127.44 (Ar), 127.20 (Ar), 127.13 (Ar), 126.80 (Ar), 126.77 (Ar), 126.53 (Ar), 123.77 (Ar), 123.16 (Ar), 123.12 (Ar), 102.54 (Glc_C1), 102.49 (Gal_C1), 99.76 (GlcN_C1), 97.85 (GalN_C1), 83.09 (Glc_C3), 82.09 (Gal_C3), 81.76 (Glc_C2), 78.87 (Gal_C2), 77.08 (GlcN_C3), 76.73 (Gal_C4), 76.58 (GalN_C3), 76.33 (GlcN_C4), 76.11 (Glc_C4), 75.52 (Ph*C*$H_2$), 75.13 (Ph*C*$H_2$), 75.11 (Ph*C*$H_2$), 74.87 (Glc_C5), 74.63 (Ph*C*$H_2$), 74.57 (GlcN_C5), 74.56 (Ph*C*$H_2$), 74.12 (Ph*C*$H_2$), 73.57 (Ph*C*$H_2$), 73.37 (GalN_C5), 73.35 (Ph*C*$H_2$), 73.25 (Gal_C5), 73.03 (Ph*C*$H_2$), 72.98 (Ph*C*$H_2$), 72.18 (GalN_C4), 71.52 (Ph*C*$H_2$), 70.93 (Ph*C*$H_2$), 68.65 (GlcN_C6), 68.49 (Gal_C6), 68.27 (GalN_C6), 67.77 (Glc_C6), 56.52 (GlcN_C2), 53.88 (GalN_C2). LRMS *m/z* calc'd for $C_{124}H_{120}N_2NaO_{23}$ $(M+Na)^+$: 2027.87; found: 2028.30.

**Benzyl 3,4,6-tri-*O*-benzyl-2-acetamido-2-deoxy-β-D-galactopyranosyl-(1→4)-2-acetamido-3,6-di-*O*-benzyl-2-deoxy-β-D-glucopyranosyl-(1→3)-2,4,6-tri-*O*-benzyl-β-D-galactopyranosyl-(1→4)-2,3,6-tri-*O*-benzyl-β-D-glucopyranoside (13)**

The starting material (**12**; 192 mg, 94.6 μmol) and $N_2H_4{\cdot}H_2O$ (65% *v/v*, 45 μL, 1.0 mmol, 10 equiv.) in EtOH (4.0 mL) were left mixing at 80 °C. After 24 h, LRMS of the mixture indicated unreacted starting material and therefore an additional 20 equiv. $N_2H_4{\cdot}H_2O$ (65% *v/v*, 90 μL, 2.0 mmol) were added to the mixture. After 48 hours at 80 °C, the mixture was evaporated to dry to afford the diamine intermediate: LRMS *m/z* calc'd for $C_{108}H_{117}N_2O_{19}$ $(M+H)^+$: 1745.83; found: 1746.17. The crude product was redissolved into MeOH (4.0 mL), and then $NaHCO_3$ (318 mg, 3.78 mmol, 40 equiv.) and $Ac_2O$ added (0.22 mL, 2.4 mmol, 25 equiv.). After 3 hours, the solution was evaporated to dry and the crude mixture purified via MPLC on silica gel using 0→50% ethyl acetate – toluene to afford the product as a colourless syrup (149 mg, 81.4 μmol, 86% yield over 2 steps). $R_f$ = 0.49 (1:1 ethyl acetate : toluene). $[\alpha]_D^{20}$: -15 (*c* 1.0,

MeOH). $^{1}$H-NMR ($CD_3CN$, 500 MHz): δH 7.40 – 7.10 (m, 60H, Ar), 6.51 (d, 1H, *J* = 9.5 Hz, NH), 6.20 (d, 1H, *J* = 9.5 Hz, NH), 4.95 (d, 1H, *J* = 11.2 Hz, PhC$H^aH^b$), 4.88 (d, 1H, *J* = 10.6 Hz, PhC$H^aH^b$), 4.85 – 4.76 (m, 4H, 4x PhC$H^aH^b$), 4.76 – 4.71 (m, 2H, 2x PhC$H^aH^b$), 4.69 (d, 1H, *J* = 7.7 Hz, GlcNAc_H1), 4.67 – 4.60 (m, 3H, PhC$H^aH^b$, PhC$H^aH^b$, PhC$H^aH^b$), 4.58 – 4.50 (m, 6H, 6x PhC$H^aH^b$), 4.48 (d, 1H, *J* = 11.3 Hz, PhC$H^aH^b$), 4.47 (d, 1H, *J* = 8.5 Hz, GalNAc_H1), 4.46 – 4.39 (m, 4H, PhC$H^aH^b$, Glc_H1, PhC$H^aH^b$, PhC$H^aH^b$), 4.38 (d, 1H, *J* = 7.7 Hz, Gal_H1), 4.33 (d, 1H, *J* = 12.1 Hz, PhC$H^aH^b$), 4.30 (d, 1H, *J* = 11.9 Hz, PhC$H^aH^b$), 4.25 (d, 1H, *J* = 12.0 Hz, PhC$H^aH^b$), 4.05 – 3.93 (m, 5H, GalNAc_H2, GalNAc_H4, Gal_H4, GlcNAc_H5, GlcNAc_H2), 3.84 – 3.79 (m, 1H, Glc_H4), 3.77 – 3.69 (m, 3H, GlcNAc_H6$^a$, GlcNAc_H6$^b$, Glc_H6$^a$), 3.64 – 3.53 (m, 5H, Gal_H3, GlcNAc_H4, GalNAc_H3, Gal_H6$^a$, Glc_H6$^b$), 3.53 – 3.40 (m, 7H, Gal_H5, Gal_H2, GlcNAc_H3, Gal_H6$^b$, GalNAc_H6$^a$, GalNAc_H5, Glc_H3), 3.37 (dd, 1H, *J* = 8.5, 5.2 Hz, GalNAc_H6$^b$), 3.30 (ddd, 1H, *J* = 9.8, 1.7, 1.7 Hz, Glc_H5), 3.23 (dd, 1H, *J* = 9.2, 7.8 Hz, Glc_H2), 1.88 (s, 3H, NHAc), 1.61 (s, 3H, NHAc). $^{13}$C-NMR ($CD_3CN$, 125 MHz): δC 171.33 (C=O), 170.28 (C=O), 140.39 (Ar), 140.25 (Ar), 140.05 (Ar), 140.04 (Ar), 140.00 (Ar), 139.92 (Ar), 139.66 (Ar), 139.63 (Ar), 139.50 (Ar), 138.86 (Ar), 129.34 (Ar), 129.30 (Ar), 129.24 (Ar), 129.22 (Ar), 129.18 (Ar), 129.16 (Ar), 129.10 (Ar), 129.00 (Ar), 128.96 (Ar), 128.88 (Ar), 128.83 (Ar), 128.79 (Ar), 128.76 (Ar), 128.75 (Ar), 128.71 (Ar), 128.67 (Ar), 128.66 (Ar), 128.61 (Ar), 128.53 (Ar), 128.50 (Ar), 128.47 (Ar), 128.45 (Ar), 128.39 (Ar), 128.37 (Ar), 128.26 (Ar), 128.18 (Ar), 128.10 (Ar), 128.03 (Ar), 104.04 (GlcNAc_C1), 103.12 (Glc_C1), 103.03 (Gal_C1), 101.26 (GalNAc_C1), 83.54 (Gal_C3), 83.46 (Glc_C3), 82.48 (Glc_C2), 81.44 (GlcNAc_C3), 80.40 (GalNAc_C3), 80.01 (Gal_C2), 77.83 (Gal_C4), 76.84 (Glc_C4), 75.84 (GlcNAc_C4), 75.56 (Ph$CH_2$), 75.54 (Ph$CH_2$), 75.50 (Ph$CH_2$), 75.49 (Ph$CH_2$), 75.36 (Glc_C5), 75.33 (GlcNAc_C5), 75.30 (Ph$CH_2$), 74.23 (Gal_C5), 74.17 (GalNAc_C5), 73.87 (2C, 2x Ph$CH_2$), 73.82 (Ph$CH_2$), 73.79 (Ph$CH_2$), 73.78 (GalNAc_C4), 73.52 (Ph$CH_2$), 72.67 (Ph$CH_2$), 71.50 (Ph$CH_2$), 70.19 (GlcNAc_C6), 69.62 (Gal_C6), 69.50 (GalNAc_C6), 68.89 (Glc_C6), 55.16 (GlcNAc_C2), 53.16 (GalNAc_C2), 23.62 (NHAc), 23.21 (NHAc). LRMS *m/z* calc'd for $C_{112}H_{120}N_2NaO_{21}$ $(M+Na)^+$: 1851.83; found: 1852.60.

**2-Acetamido-2-deoxy-β-D-galactopyranosyl-(1→4)-2-acetamido-2-deoxy-β-D-glucopyranosyl-(1→3)-β-D-galactopyranosyl-(1→4)-D-glucopyranose (LdiNnT)**

The starting material **13** (56 mg, 31 µmol) and palladium on activated carbon (Pd/C 10% *w/w*, 55 mg) were suspended in 9:1 methanol–$H_2O$ (3.0 mL), the atmosphere evacuated, and the flask flushed with $H_2$(g) (via balloon). After 24 h at rt, LRMS of the mixture indicated complete deprotection of starting material. The catalyst was filtered through Celite washing with $H_2O$ (3 mL). The filtrate was evaporated to dry, and the crude material purified by passing through a C-18 reverse phase cartridge, to afford the pure product as a white solid in a 0.5 : 1 α/β anomeric mixture, as determined by integration of the corresponding NMR signals, (21.1 mg, 28.2 μmol, 92% yield). $R_f$ = 0.21 (7:2:2 *n*-propanol : ethanol : $H_2O$). $[\alpha]_D^{20}$: +27.8 (*c* 1.0, $H_2O$). $^1$H-NMR ($D_2O$, 500 MHz) δH 5.21 (d, *J* = 3.8 Hz, Glc_H1α), 4.68 (d, *J* = 8.2 Hz, GlcNAc_H1α), 4.68 (d, *J* = 8.2 Hz, GlcNAc_H1β), 4.65 (d, *J* = 8.0 Hz, Glc_H1β), 4.52 (d, *J* = 8.5 Hz, GalNAc_H1α/β), 4.43 (d, *J* = 7.9 Hz, Gal_H1α/β), 4.13 (dd, *J* = 3.4, <1 Hz, Gal_H4α/β), 3.96 – 3.90 (m, Glc_H6$^a$β, Glc_H5α, GalNAc_H4α/β, GalNAc_H2α/β), 3.88 (dd, *J* = 12.2, 2.3 Hz, Glc_H6$^a$α), 3.85 – 3.69 (m, GlcNAc_H6$^a$α/β, Glc_H3α, Gal_H6$^a$α/β, Glc_H6$^b$α, GalNAc_H6$^a$α/β, Glc_H6$^b$β, GlcNAc_H2α/β, Gal_H6$^b$α/β, GalNAc_H6$^b$α/β, GalNAc_H3α/β, GlcNAc_H3α/β, GalNAc_H5α/β, Gal_H3α/β, Gal_H5α/β), 3.67 – 3.55 (m, GlcNAc_H6$^b$α/β, GlcNAc_H4α/β, Glc_H3β, Glc_H4α/β, Glc_H5β, Gal_H2α/β, Glc_H2α), 3.50 (ddd, *J* = 9.7, 4.9, 2.2 Hz, GlcNAc_H5α/β), 3.30 – 3.24 (m, 1H, Glc_H2β), 2.06 (s, GalNAc_Acα/β), 2.02 (s, GlcNAc_Acα/β). $^{13}$C-NMR ($D_2O$, 125 MHz): δC 175.58 (GlcNAc_C=Oα/β), 175.44 (GalNAc_C=Oα/β), 103.58 (Gal_C1β), 103.55 (Gal_C1α), 103.37 (GlcNAc_C1α/β), 102.38 (GalNAc_C1α/β), 96.40 (Glc_C1β), 92.47 (Glc_C1α), 82.71 (Gal_C3β), 82.70 (Gal_C3α), 79.57 (GlcNAc_C4α/β), 79.04 (Glc_C4α), 78.93 (Glc_C4β), 75.98 (GalNAc_C5α/β), 75.52, 75.46 (Gal_C5α/β and Glc_C5β), 75.01 (Glc_C3β), 74.98 (GlcNAc_C5α/β), 74.45 (Glc_C2β), 72.94 (GlcNAc_C3α/β), 72.06 (Glc_C3α), 71.80 (Glc_C2α), 71.33 (GalNAc_C3α/β), 70.78 (Glc_C5α), 70.65 (Gal_C2α), 70.63 (Gal_C2β), 69.01 (Gal_C4α), 68.98 (Gal_C4β), 68.25 (GalNAc_C4α/β), 61.61 (Gal_C6α/β and GalNAc_C6α/β), 60.74 (Glc_C6β), 60.62 (GlcNAc_C6α/β and Glc_C6α), 55.61 (GlcNAc_C2α/β), 53.20 (GalNAc_C2α/β), 22.84 (GalNAc_Acα/β), 22.83 (GlcNAc_Acα/β). ESI-HRMS *m/z* calc'd for $C_{28}H_{48}N_2NaO_{21}$ $(M+Na)^+$: 771.2647; found: 771.2647. HPLC purity analysis: 99.3%, $R_t$ 7.35 minutes, EC 250/4 Nucleodur C18 column.

<u>References</u>

[1] G. R. Fulmer *et al. Organometallics* **2010**, *29*, 9, 2176 – 2179, doi: 10.1021/om100106e.

[2] P. Morrone-Pozzuto, M. L. Uhrig, and R. Agusti, *J. Org. Chem.* **2022**, *87*, 20, 13455 – 13468, doi: 10.1021/acs.joc.2c01059.

[3] K. Matsuoka *et al.*, *Carbohydr. Polym.* **2007**, *69*, 2, 326 – 335, doi: 10.1016/j.carbpol.2006.10.011.

[4] G. Minzer and R. Hevey, *ChemistryOpen* **2022**, *12*, e20220134, doi: 10.1002/open.202200134.

[5] J. Veselý, M. Ledvina, J. Jindřich, D. Šaman, and T. Trnka, *Collect. Czechoslov. Chem. Commun.* **2003**, *68*, 7, 1264 – 1274, doi: 10.1135/cccc20031264.

### NMR spectra for compound 2

$^1H$ NMR in $CD_3OD$ (500 MHz)

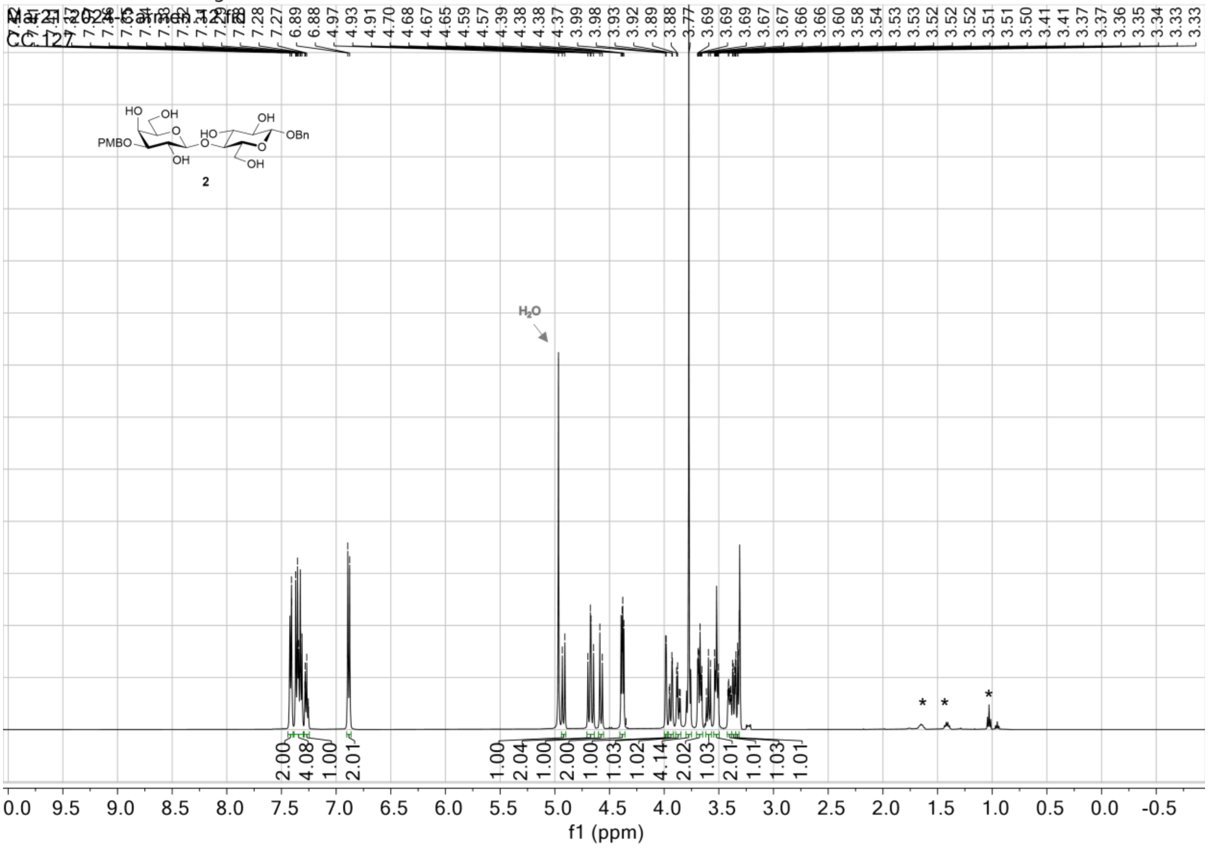

$^{13}$C NMR (APT) in $CD_3OD$ (125 MHz)

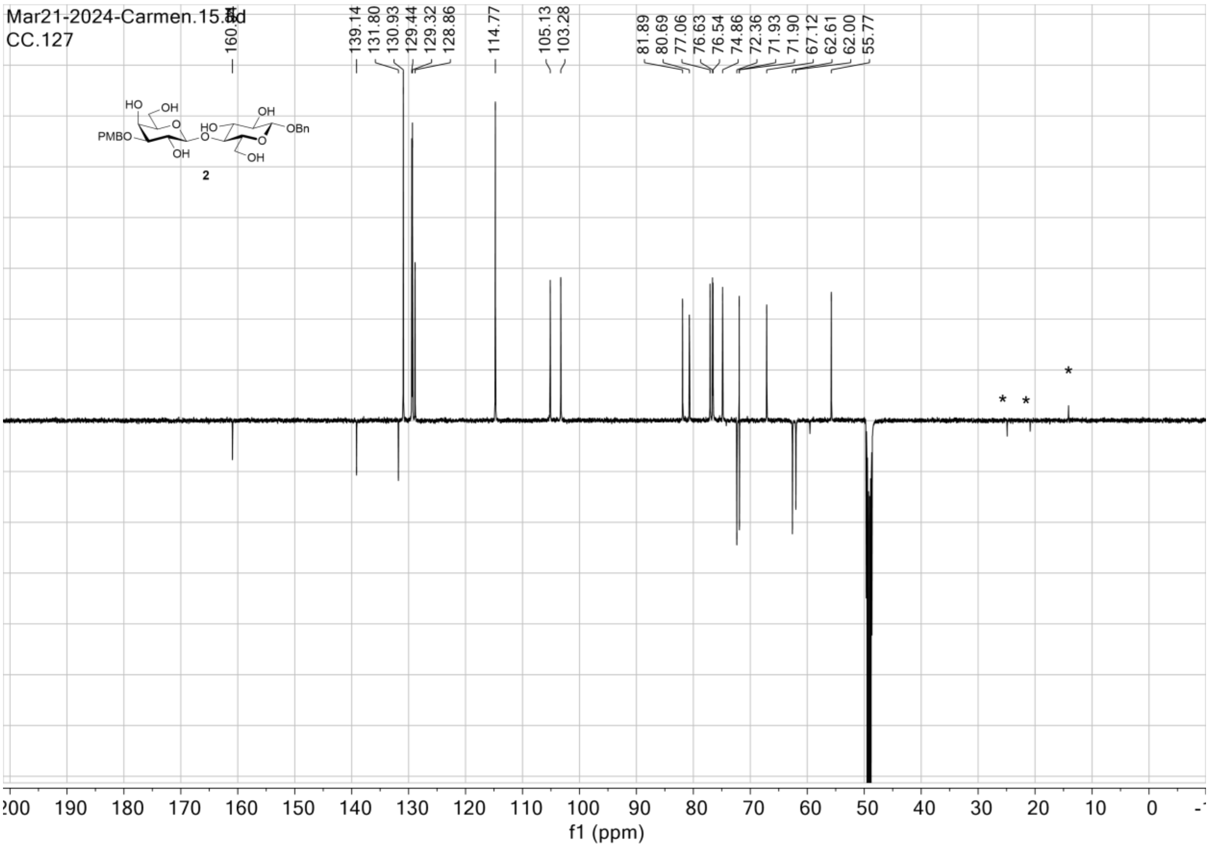

$^{1}$H\_$^{1}$H gCOSY in $CD_3OD$

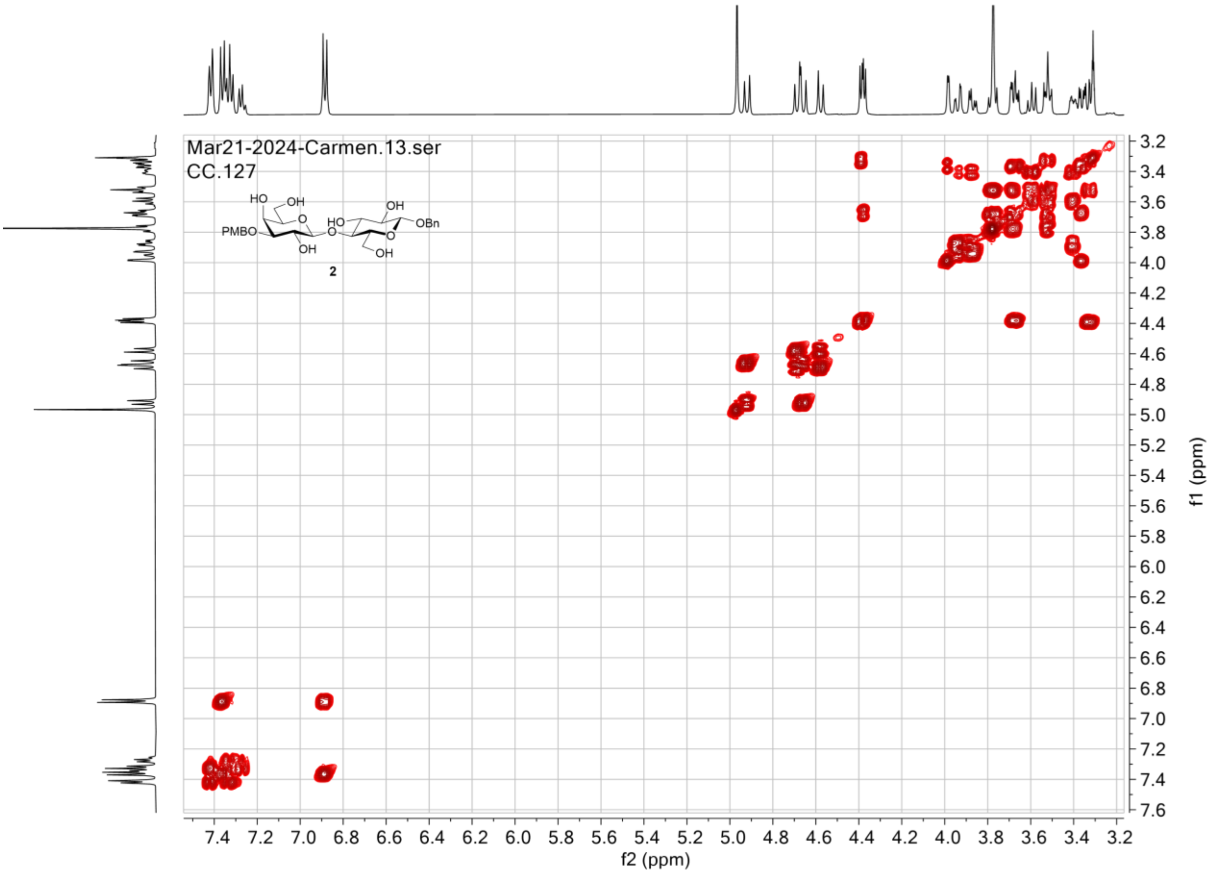

$^{1}H\_^{13}C$ gHSQC in $CD_3OD$

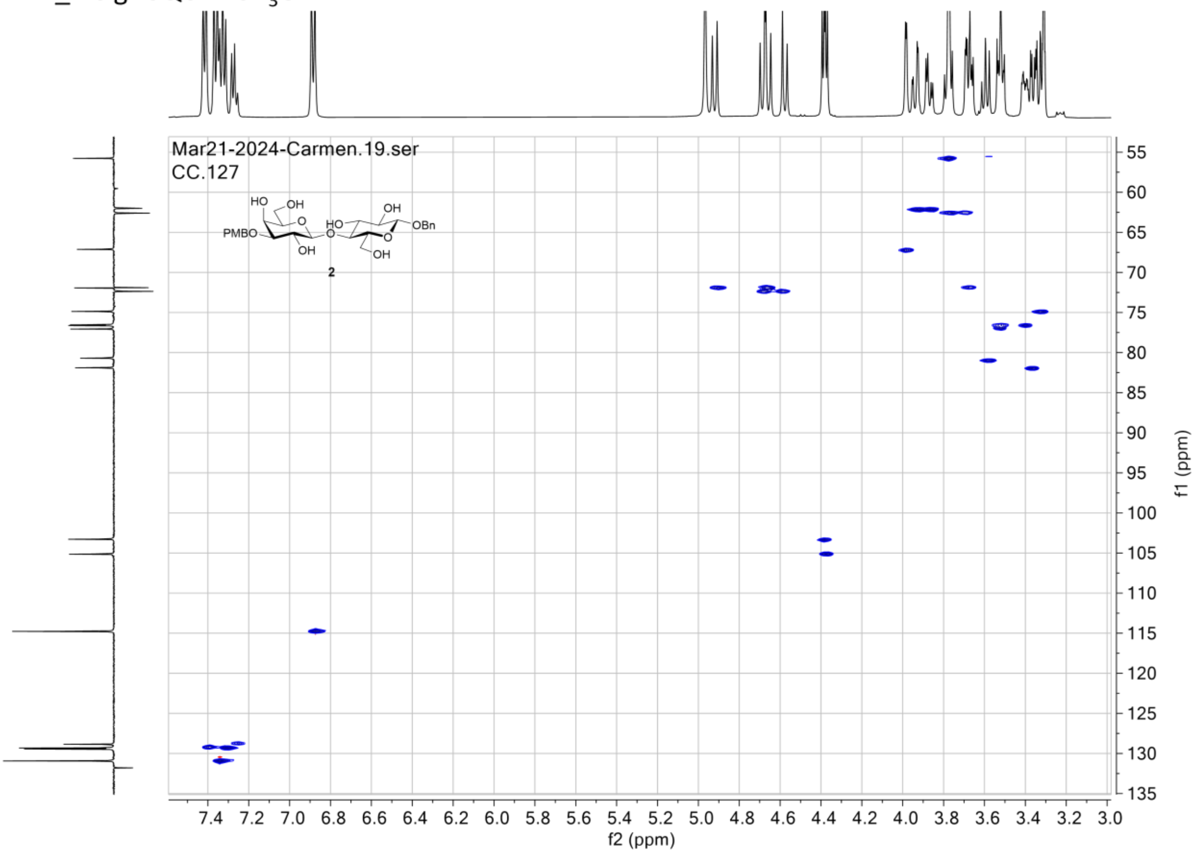

### NMR spectra for compound 3

$^1H$ NMR in $CDCl_3$ (500 MHz)

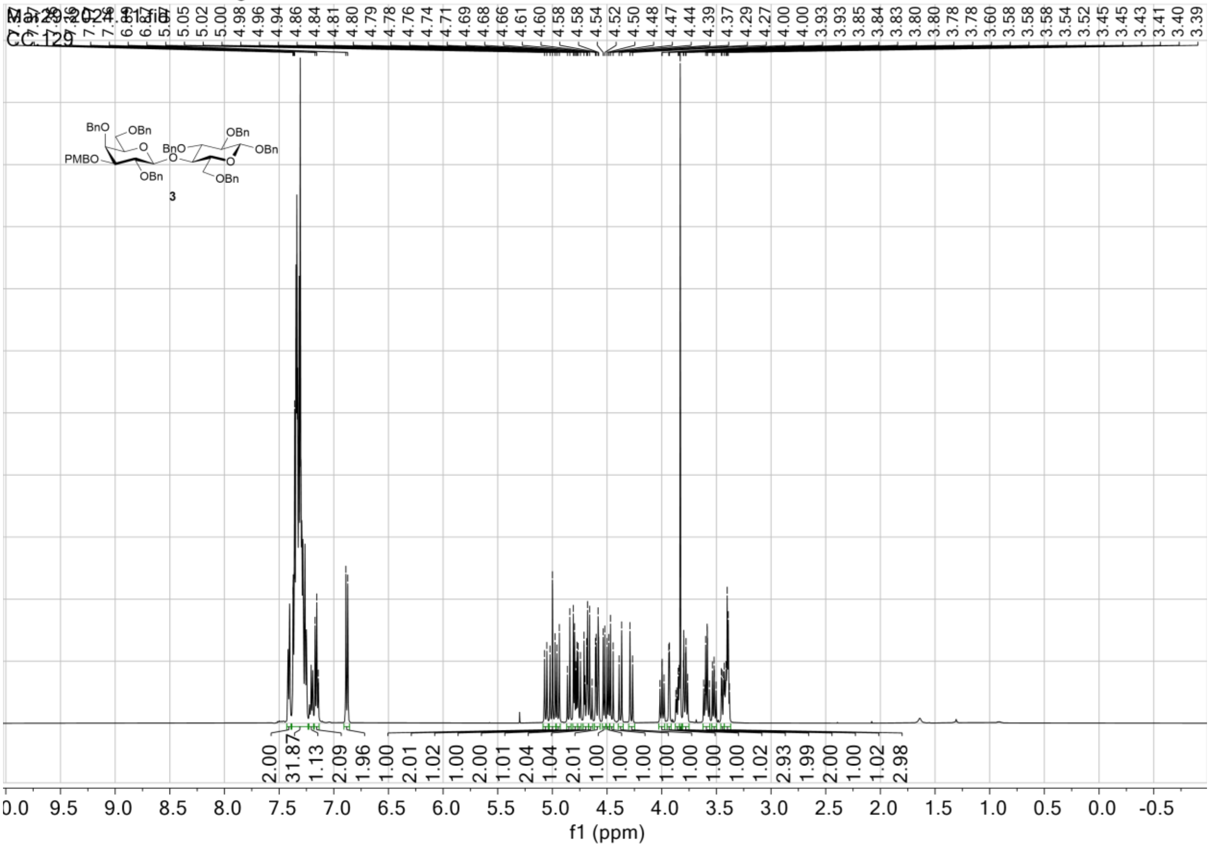

$^{13}C$ NMR (APT) in $CDCl_3$ (125 MHz)

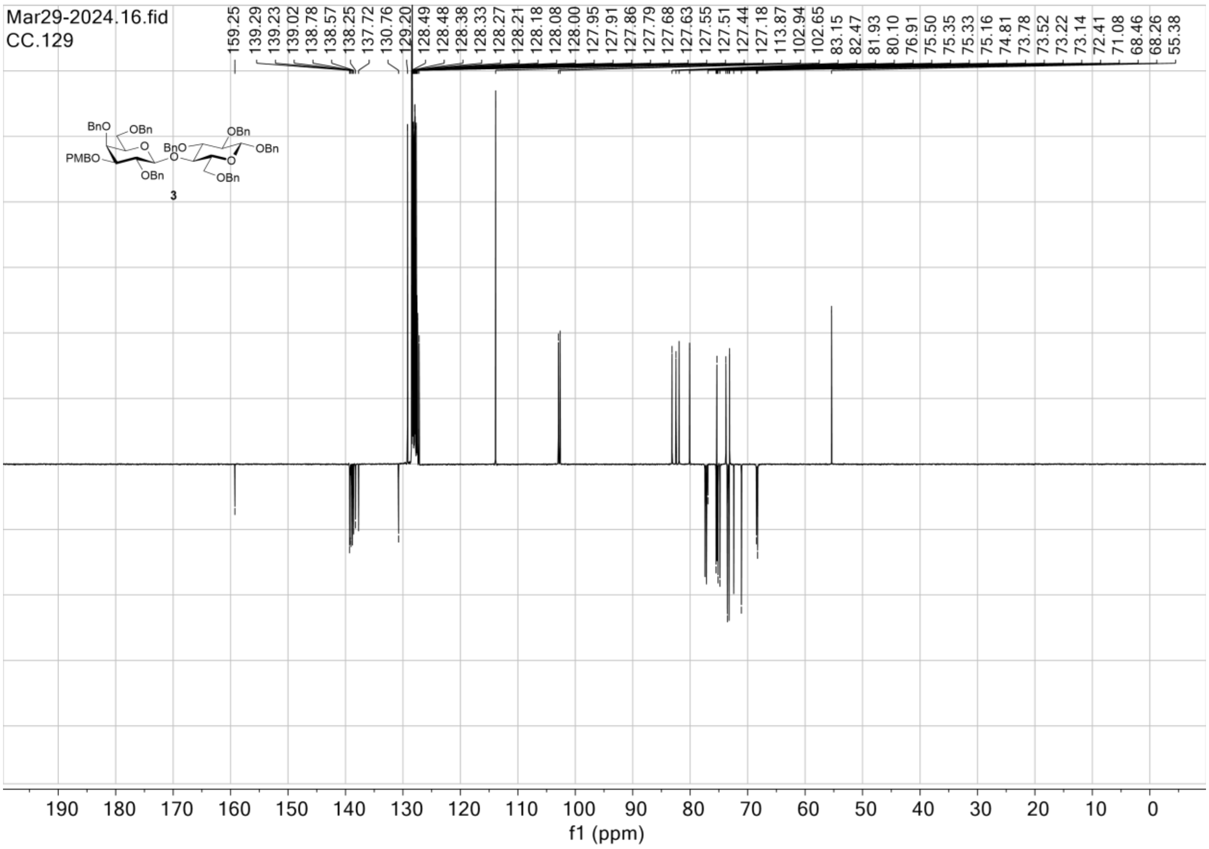

$^1H\_^1H$ gCOSY in $CDCl_3$

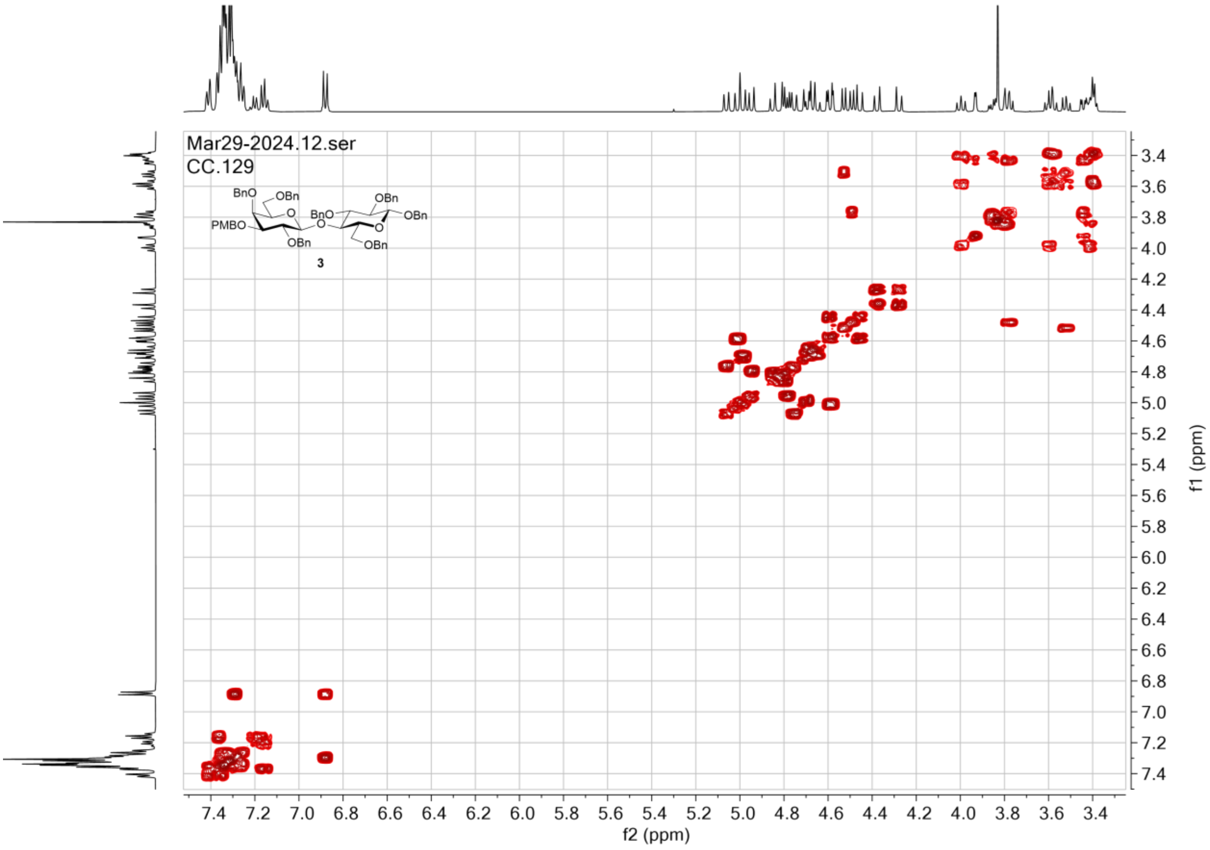

$^{1}H\_^{13}C$ gHSQC in $CDCl_3$

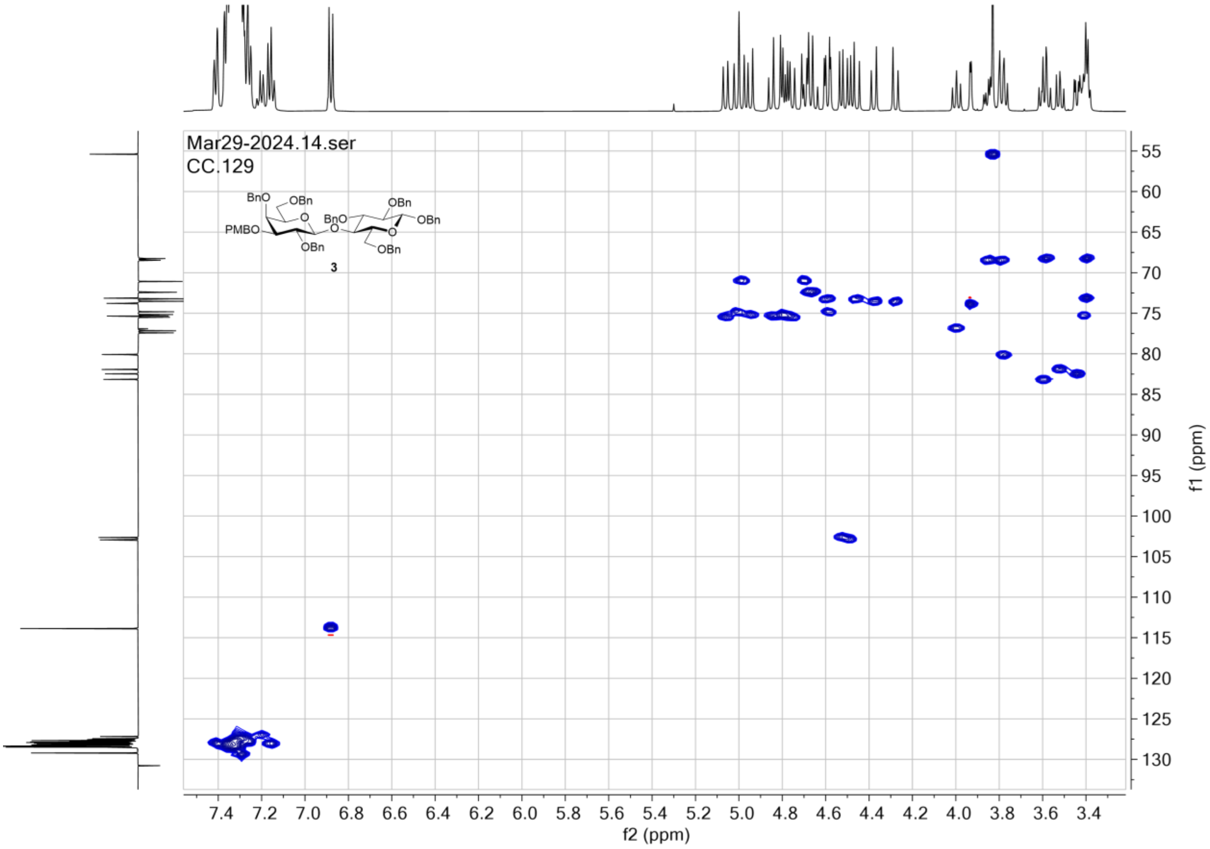

### NMR spectra for compound 4

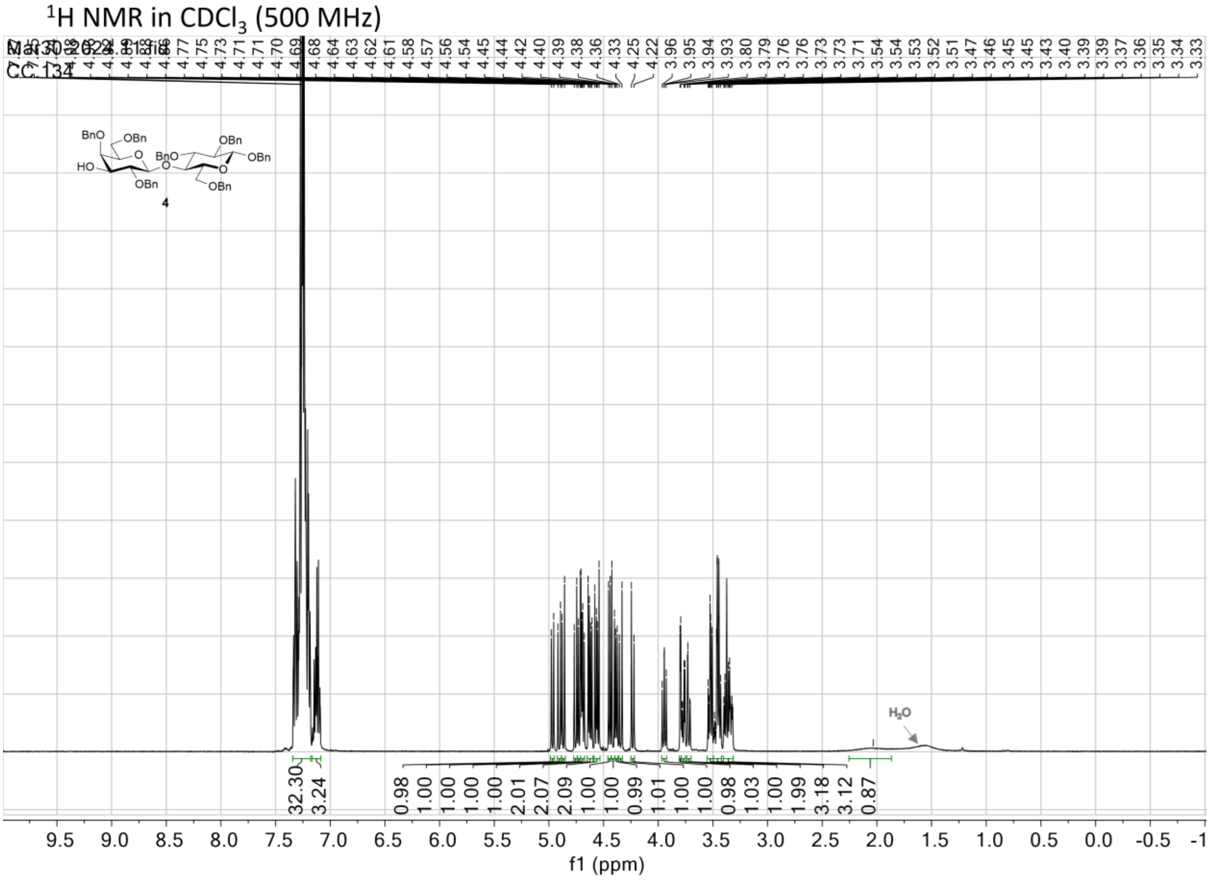

$^{13}C$ NMR (APT) in $CDCl_3$ (125 MHz)

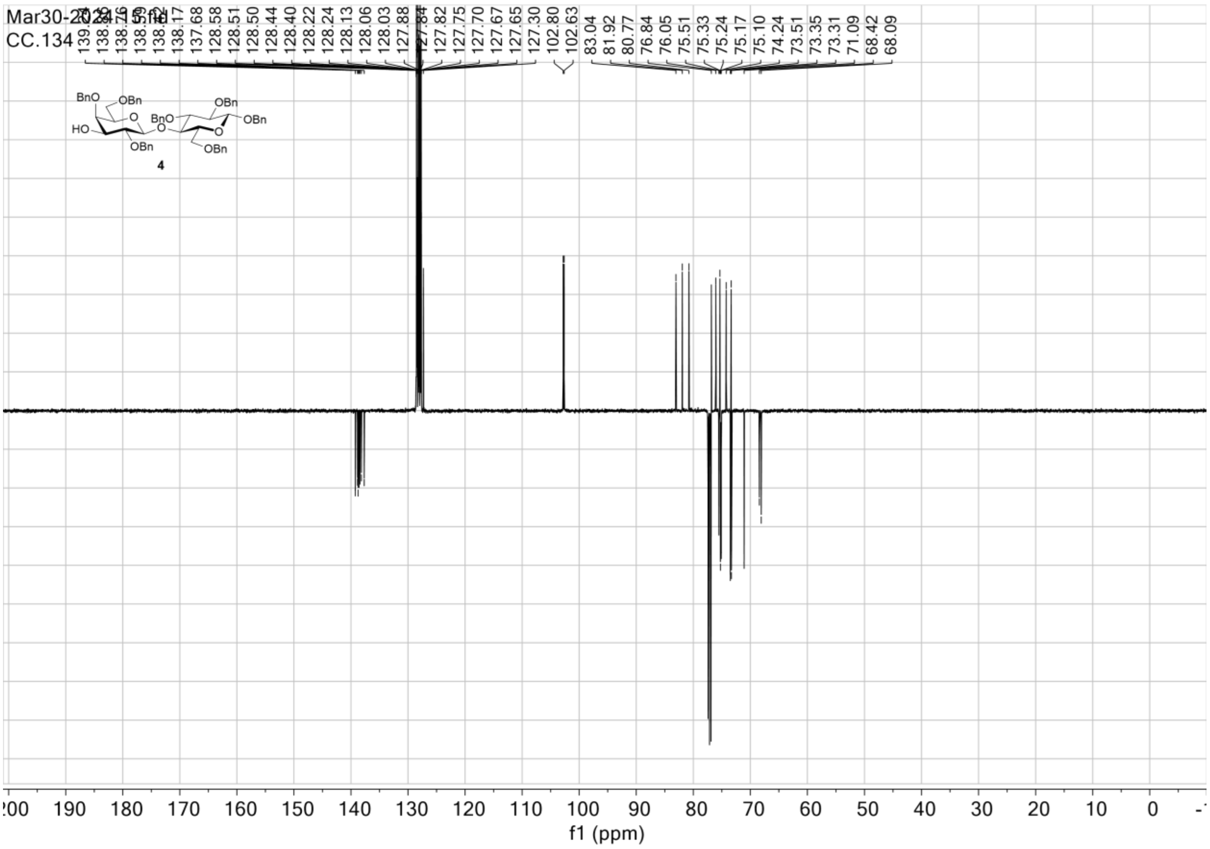

$^1H$_$^1H$ gCOSY in $CDCl_3$

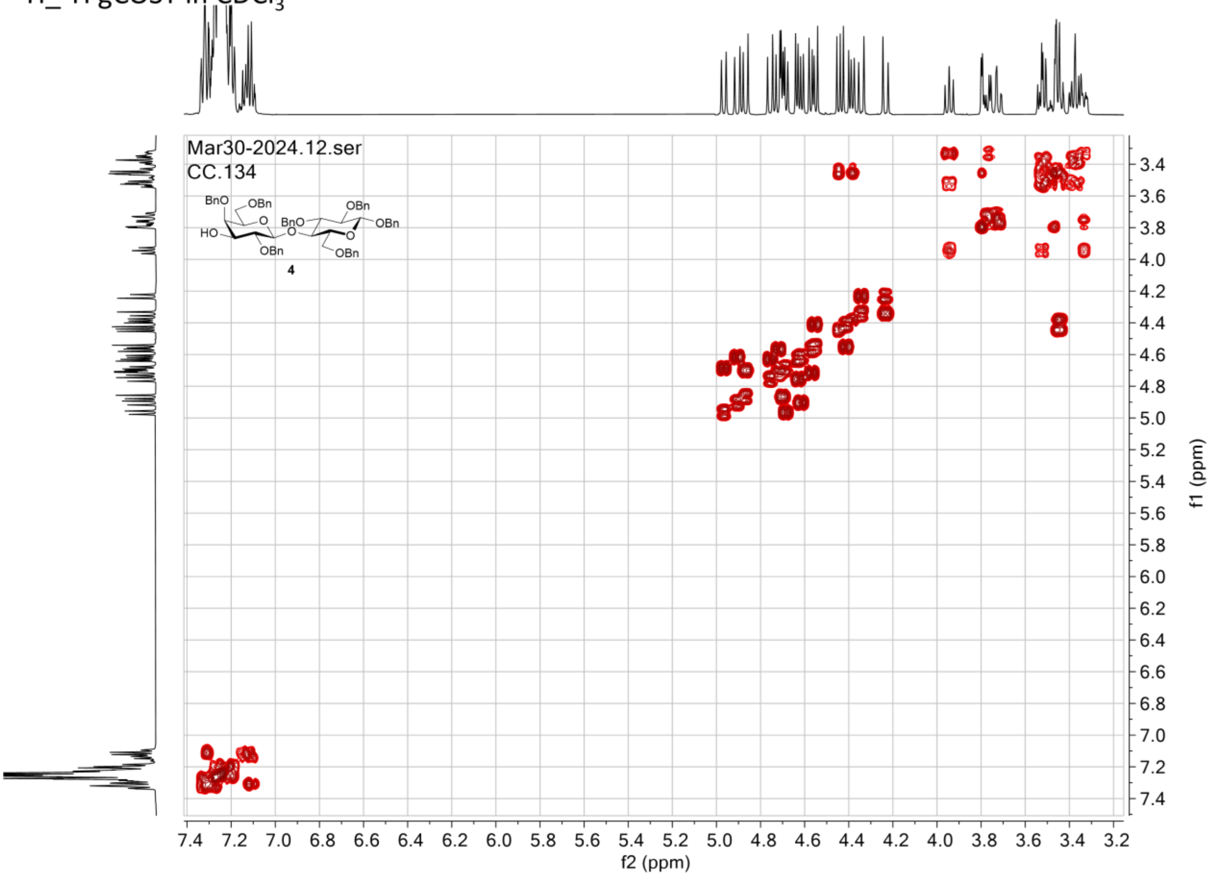

$^{1}H\_^{13}C$ gHSQC in $CDCl_3$

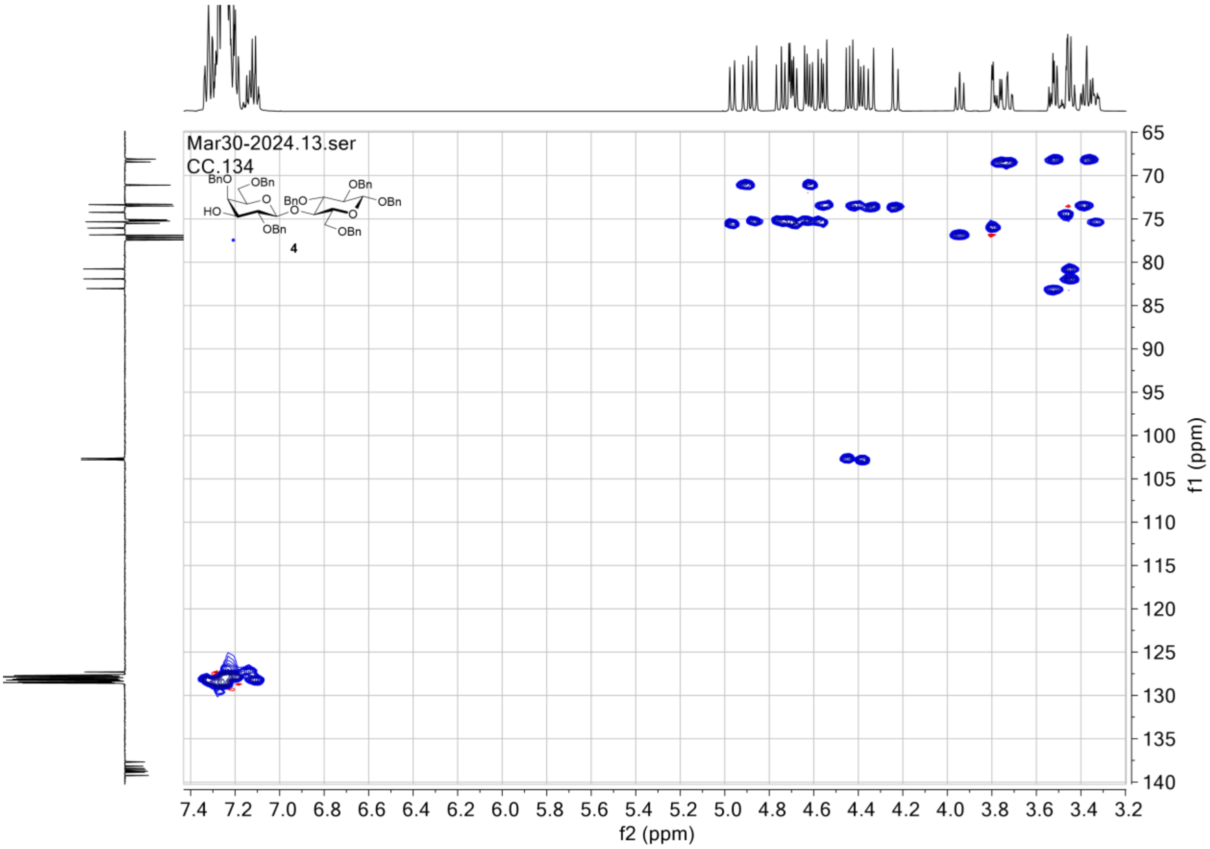

### NMR spectra for compound 6

$^1$H NMR in $CDCl_3$ (500 MHz)

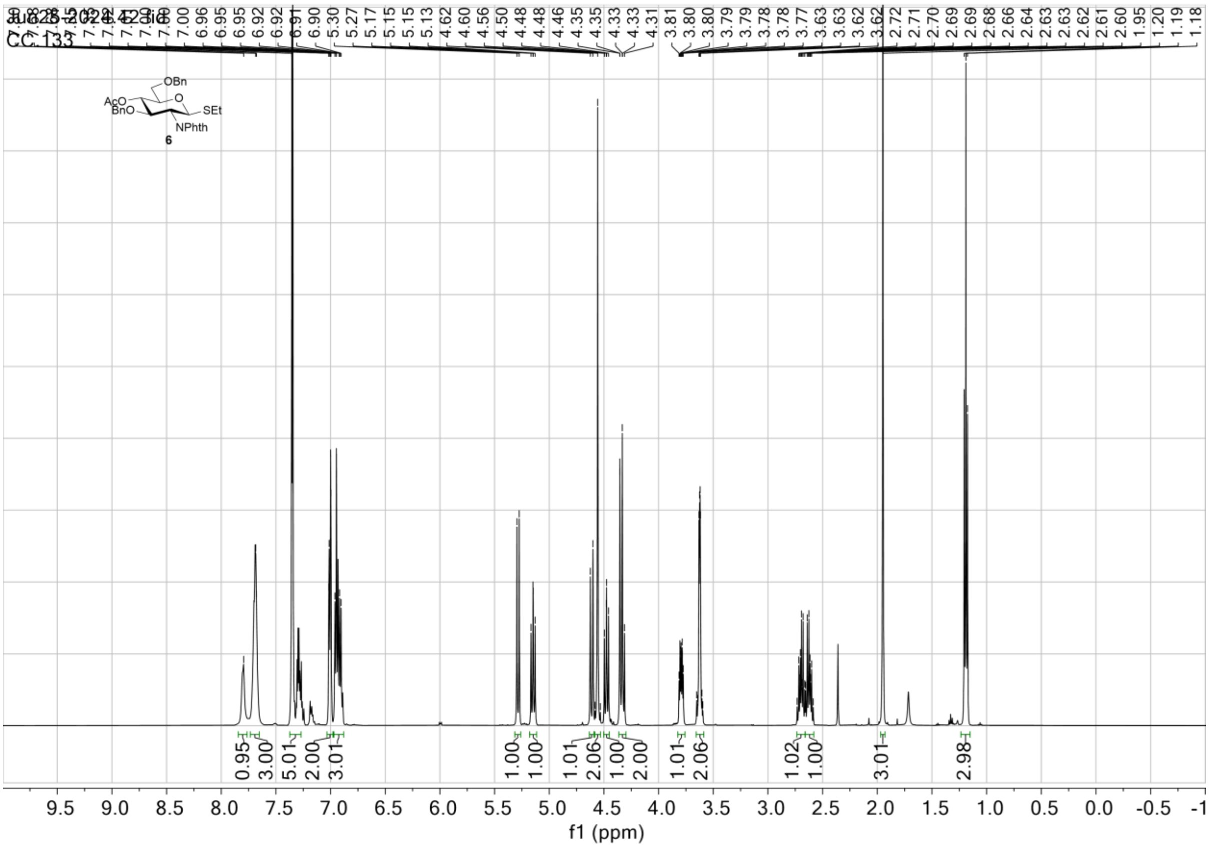

$^{13}C$ NMR (APT) in $CDCl_3$ (125 MHz)

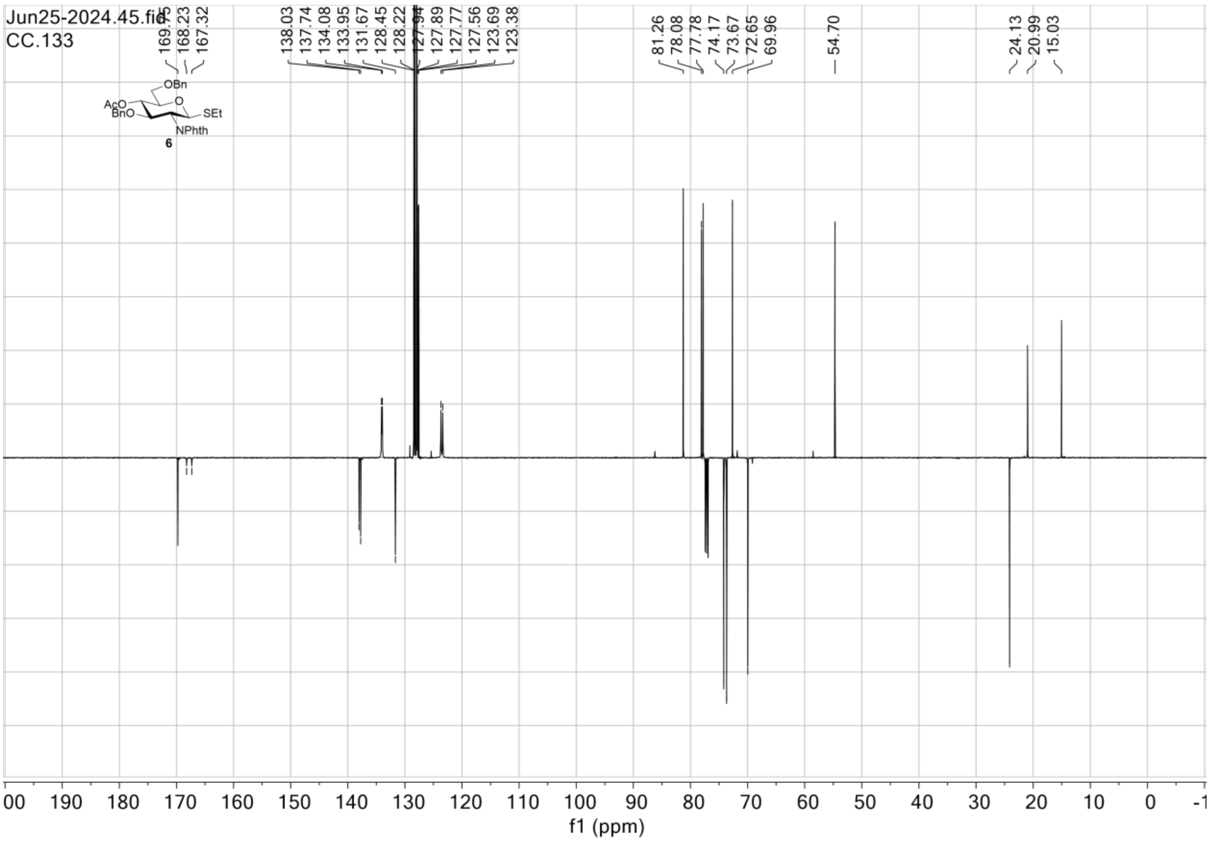

$^1H\_^1H$ gCOSY in $CDCl_3$

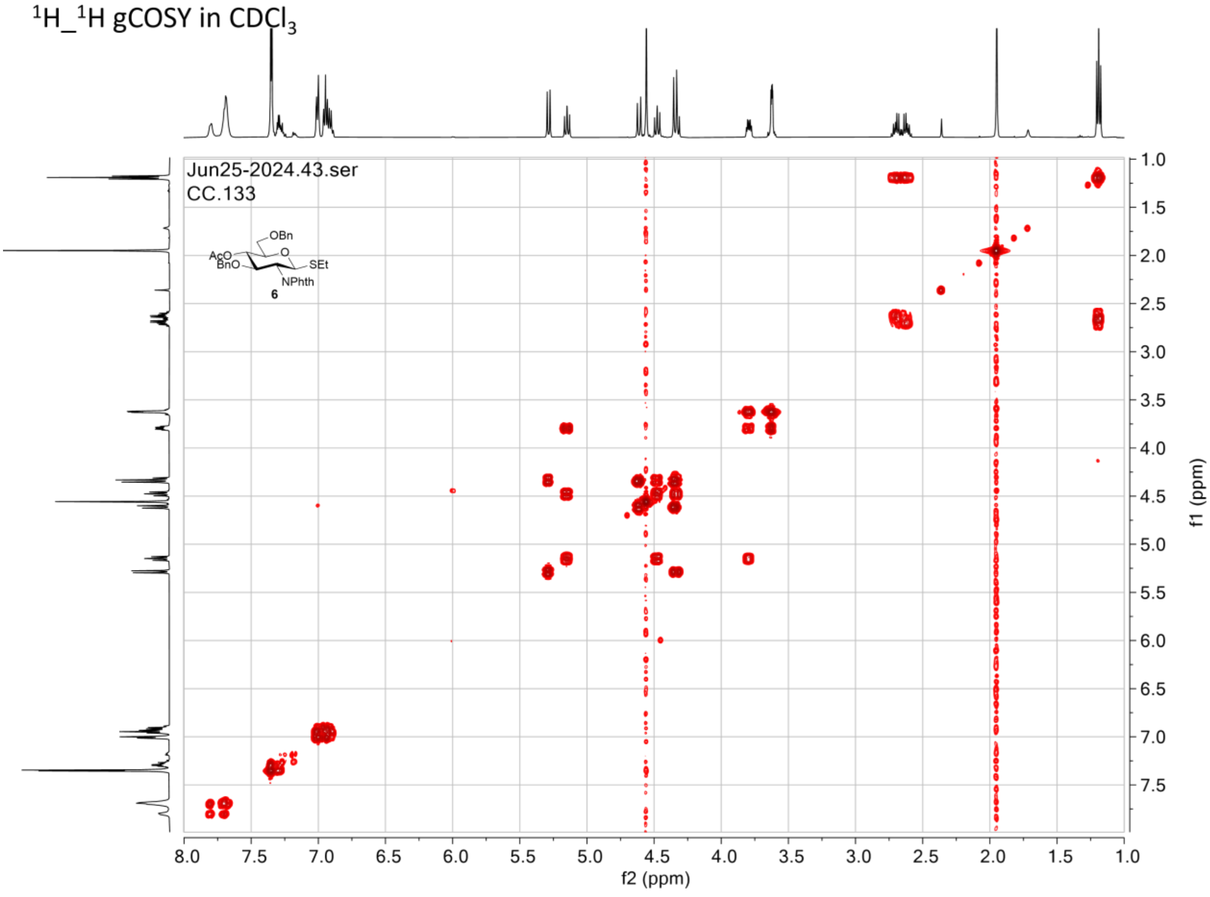

$^{1}H\_^{13}C$ gHSQC in $CDCl_3$

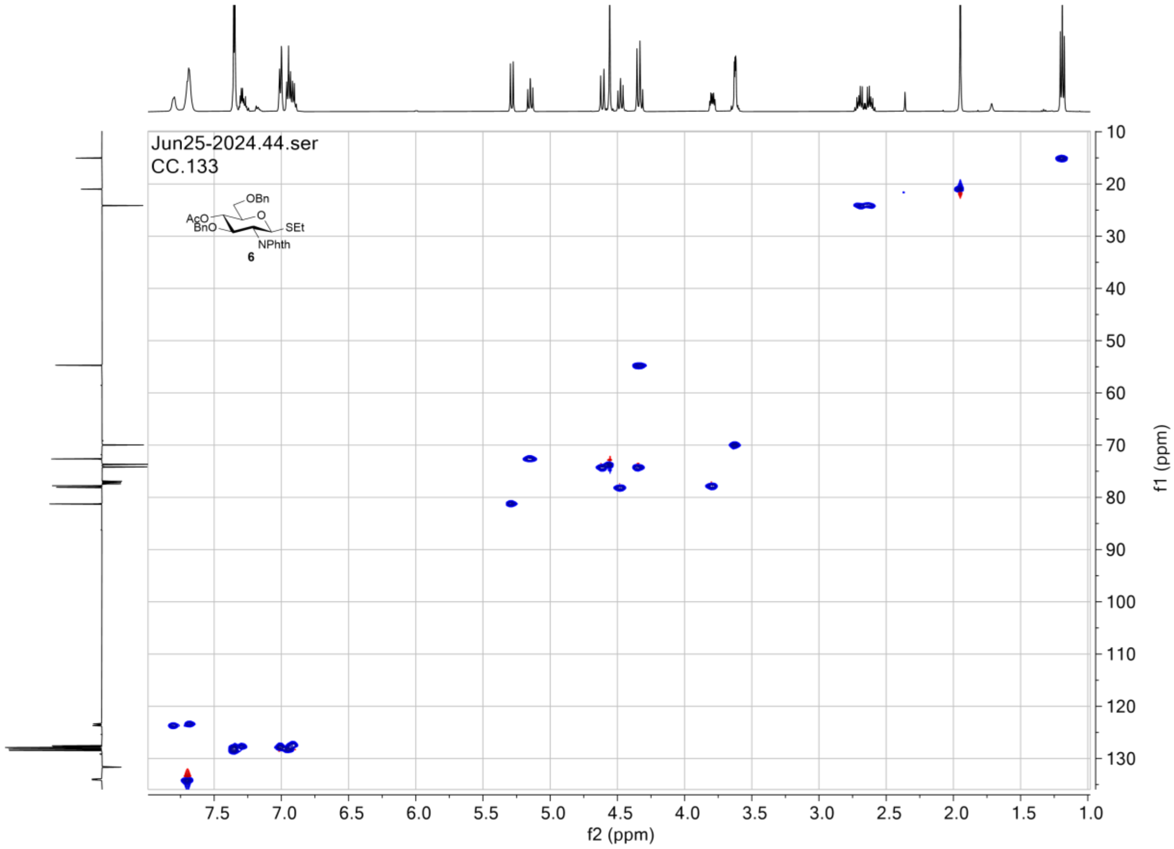

### NMR spectra for compound 8

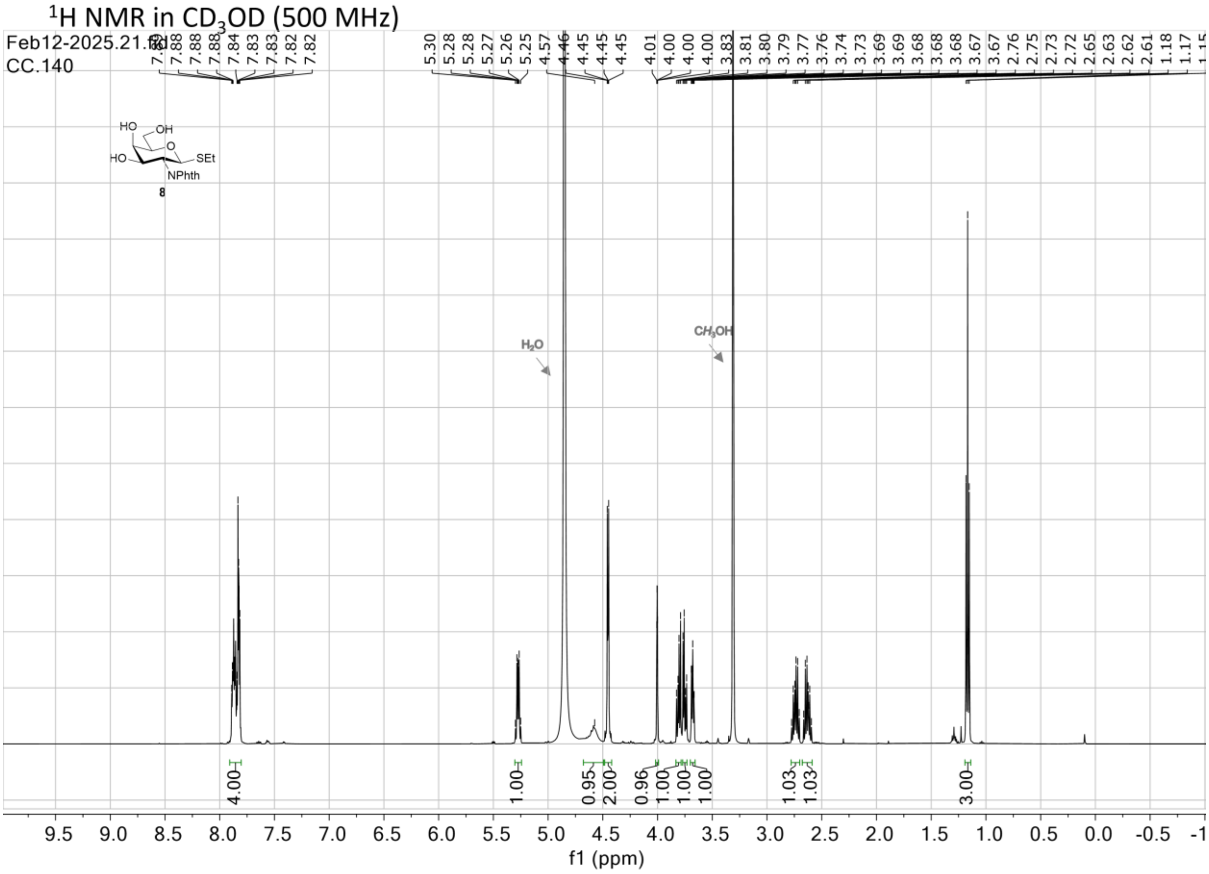

$^{13}C$ NMR (APT) in $CD_3OD$ (125 MHz)

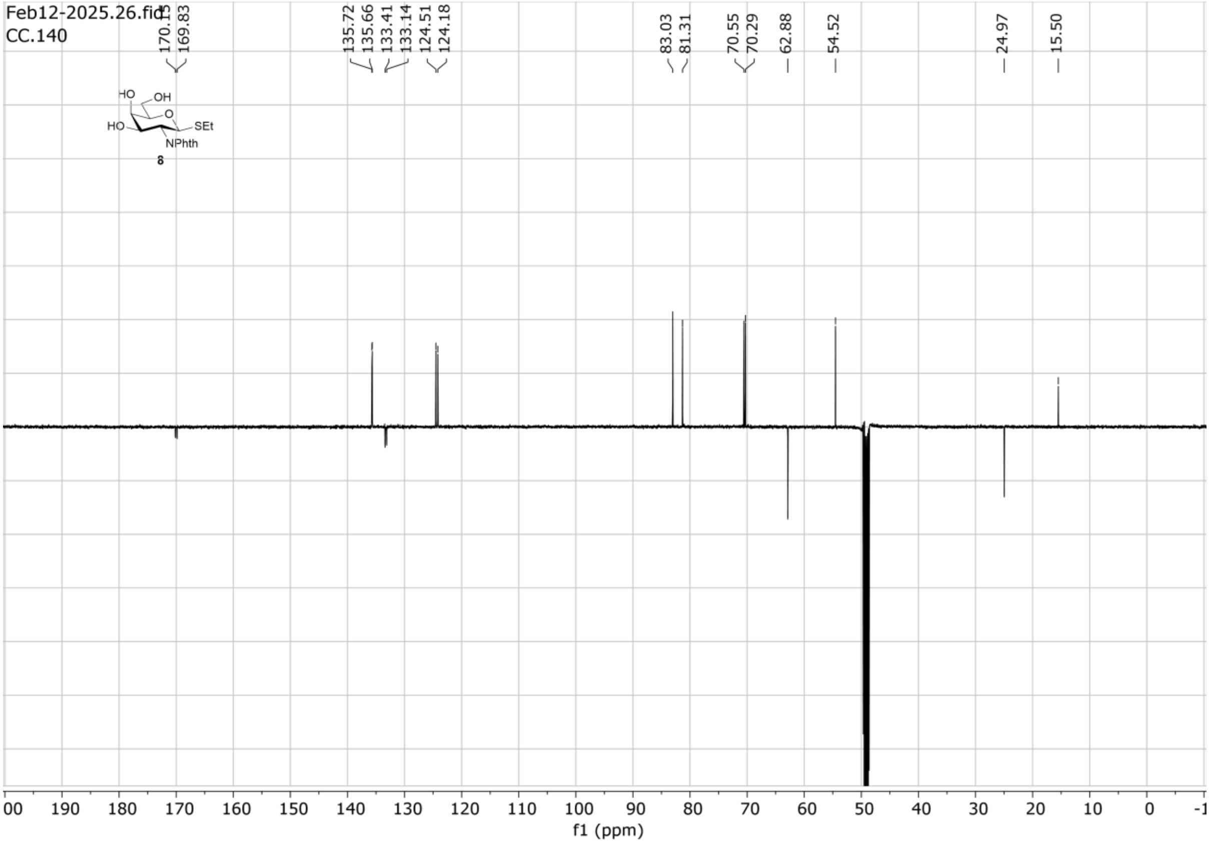

$^1H\_^1H$ gCOSY in $CD_3OD$

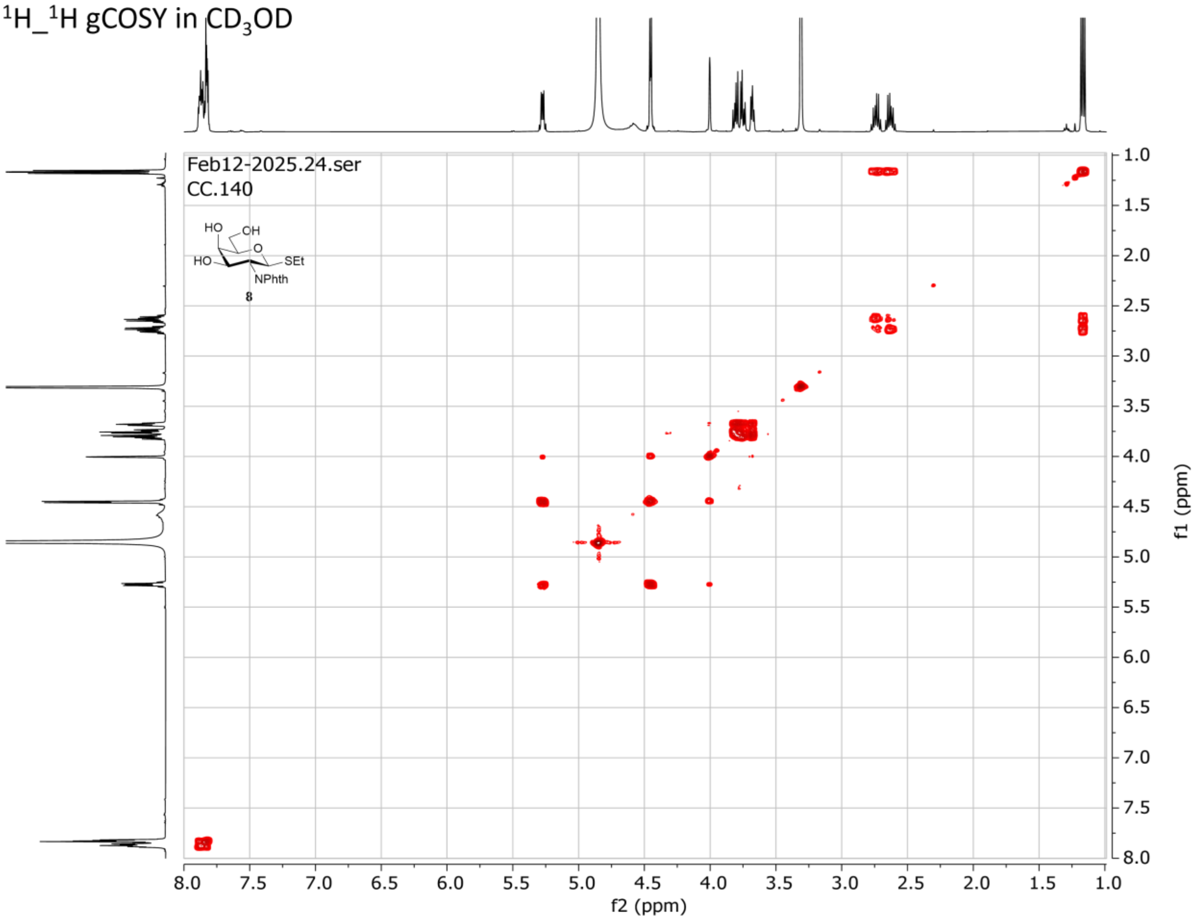

$^1H\_^{13}C$ gHSQC in $CD_3OD$

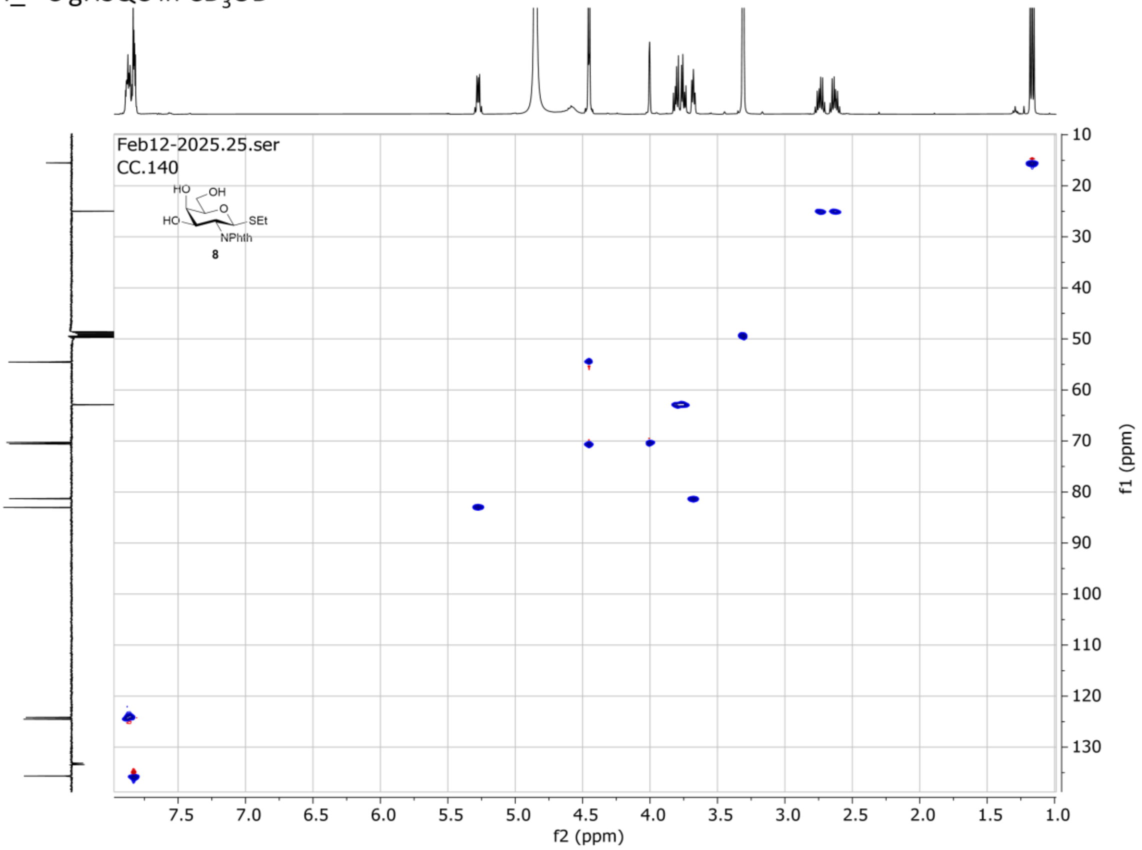

### NMR spectra for compound 9

$^1$H NMR in $CD_3CN$ (500 MHz)

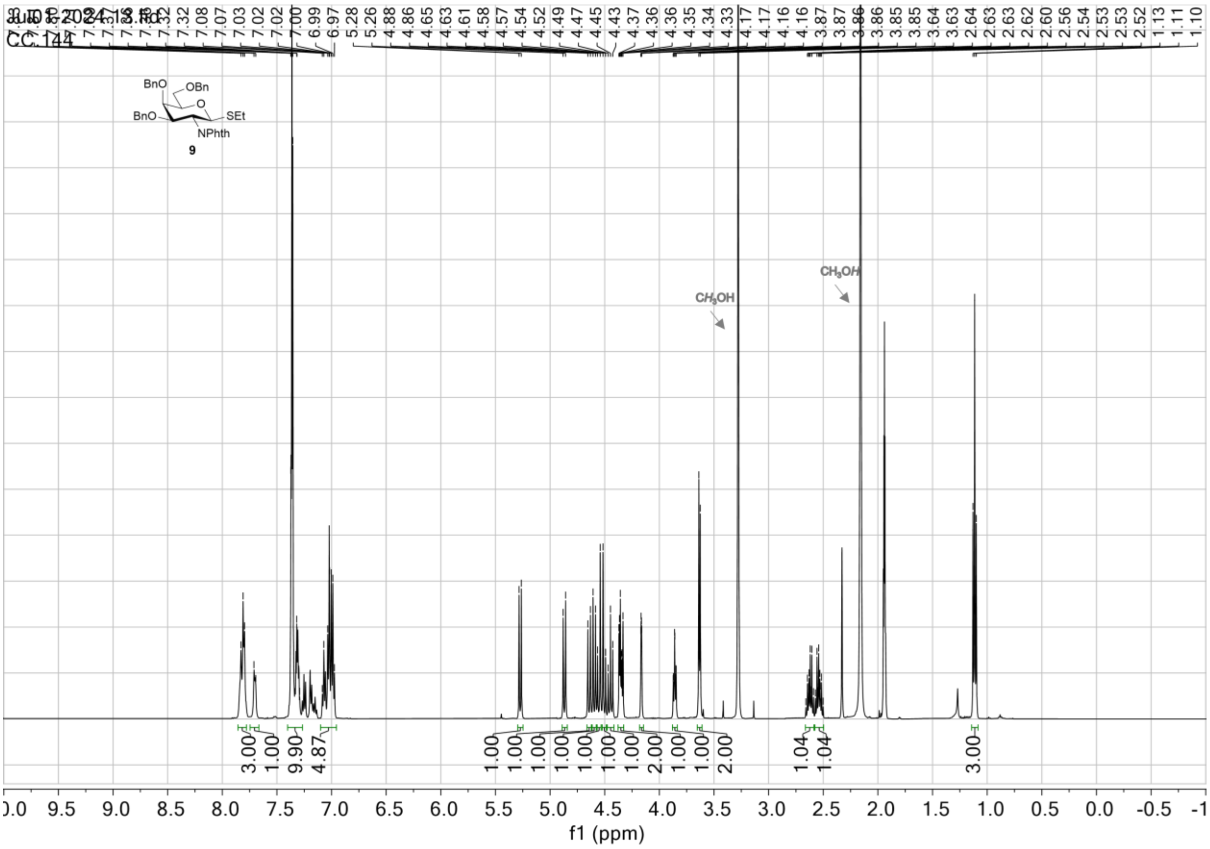

$^{13}C$ NMR (APT) in $CD_3CN$ (125 MHz)

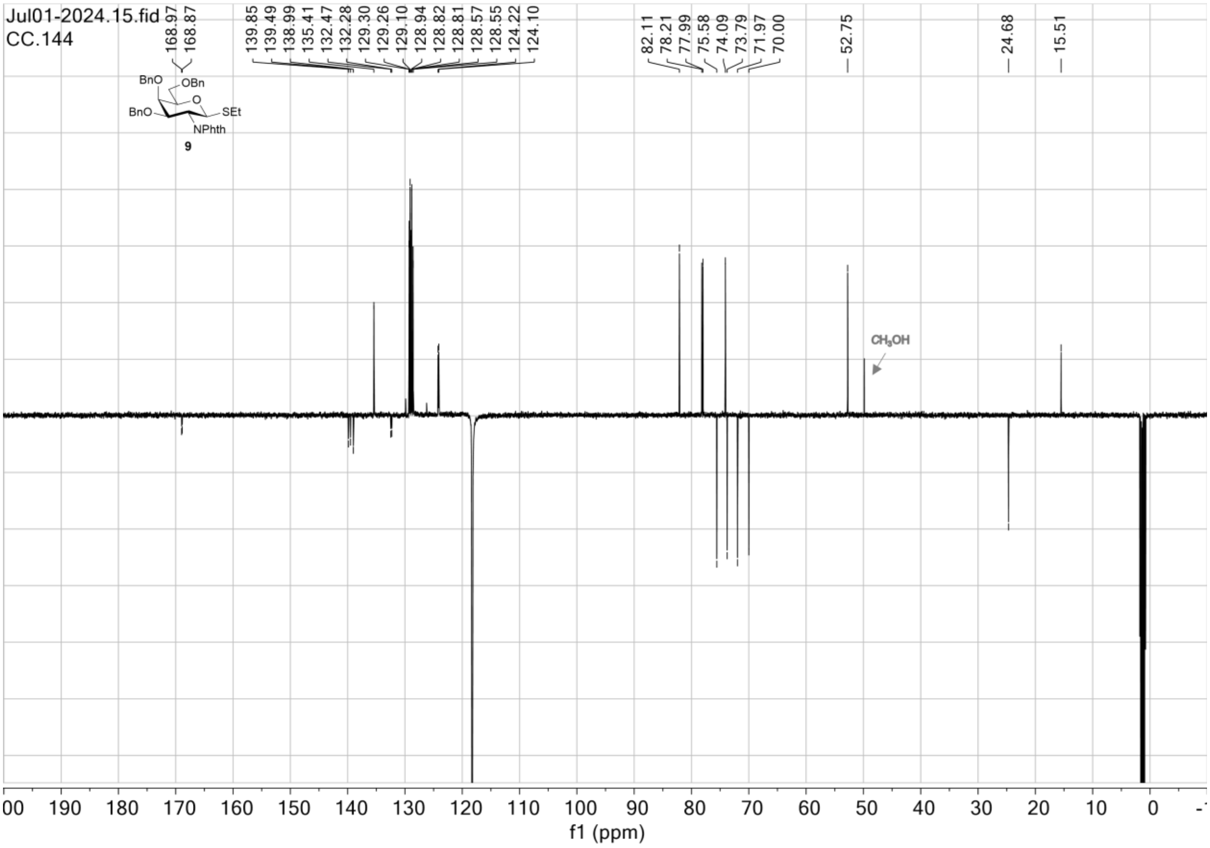

$^1H\_^1H$ gCOSY in $CD_3CN$

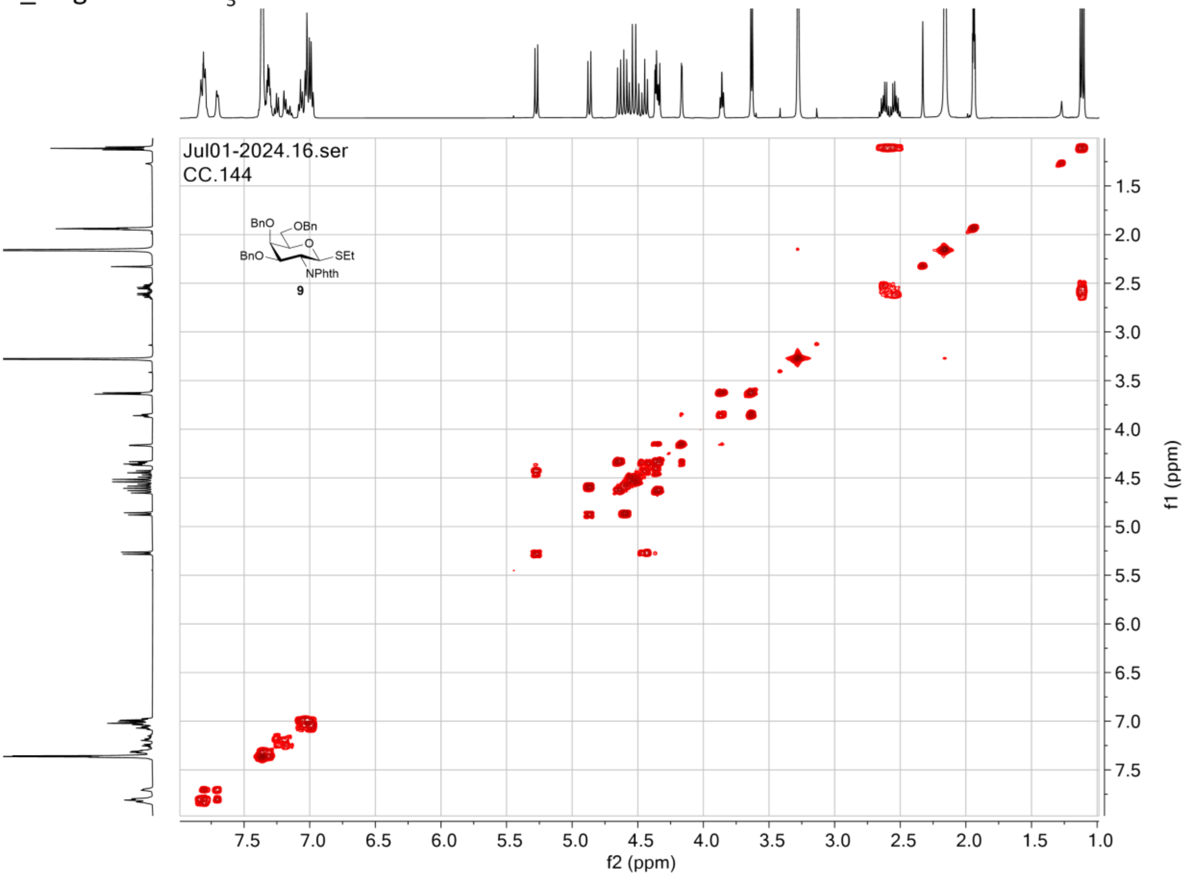

$^1H\_^{13}C$ gHSQC in $CD_3CN$

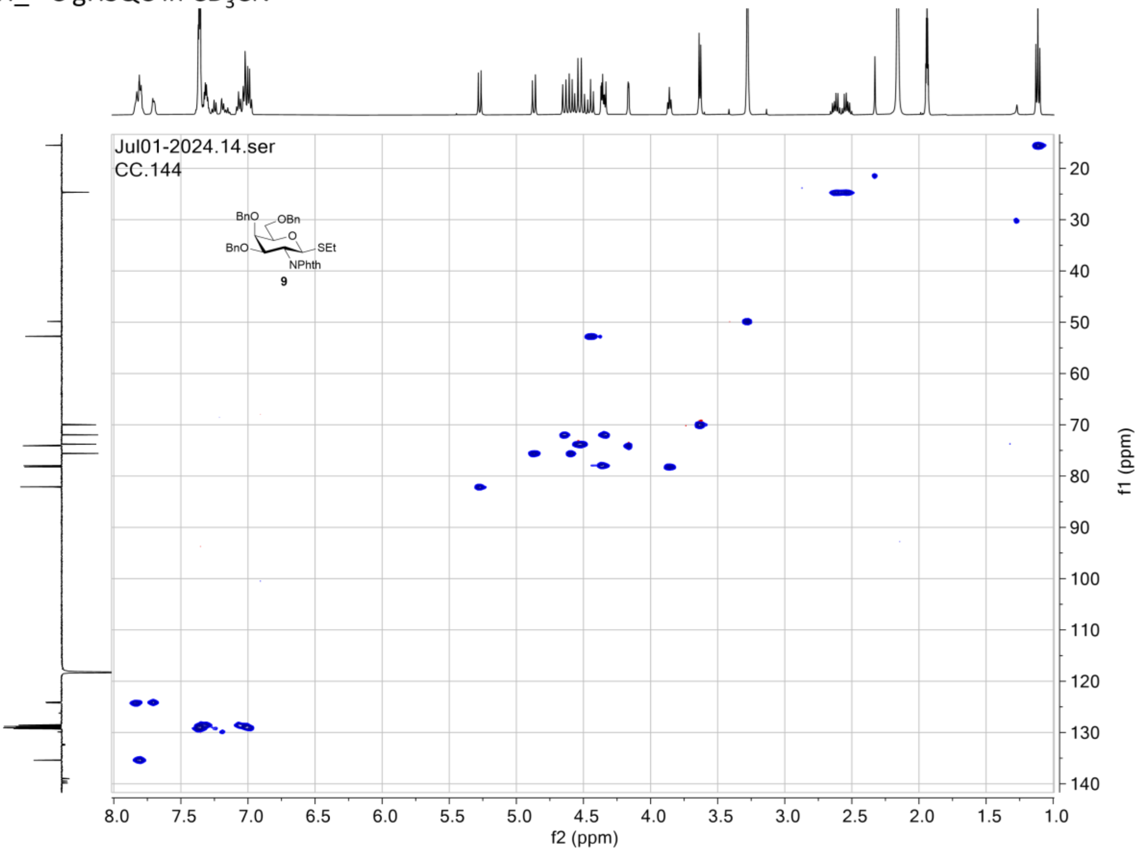

### NMR spectra for compound 10

$^{1}$H NMR in $CDCl_3$ (500 MHz)

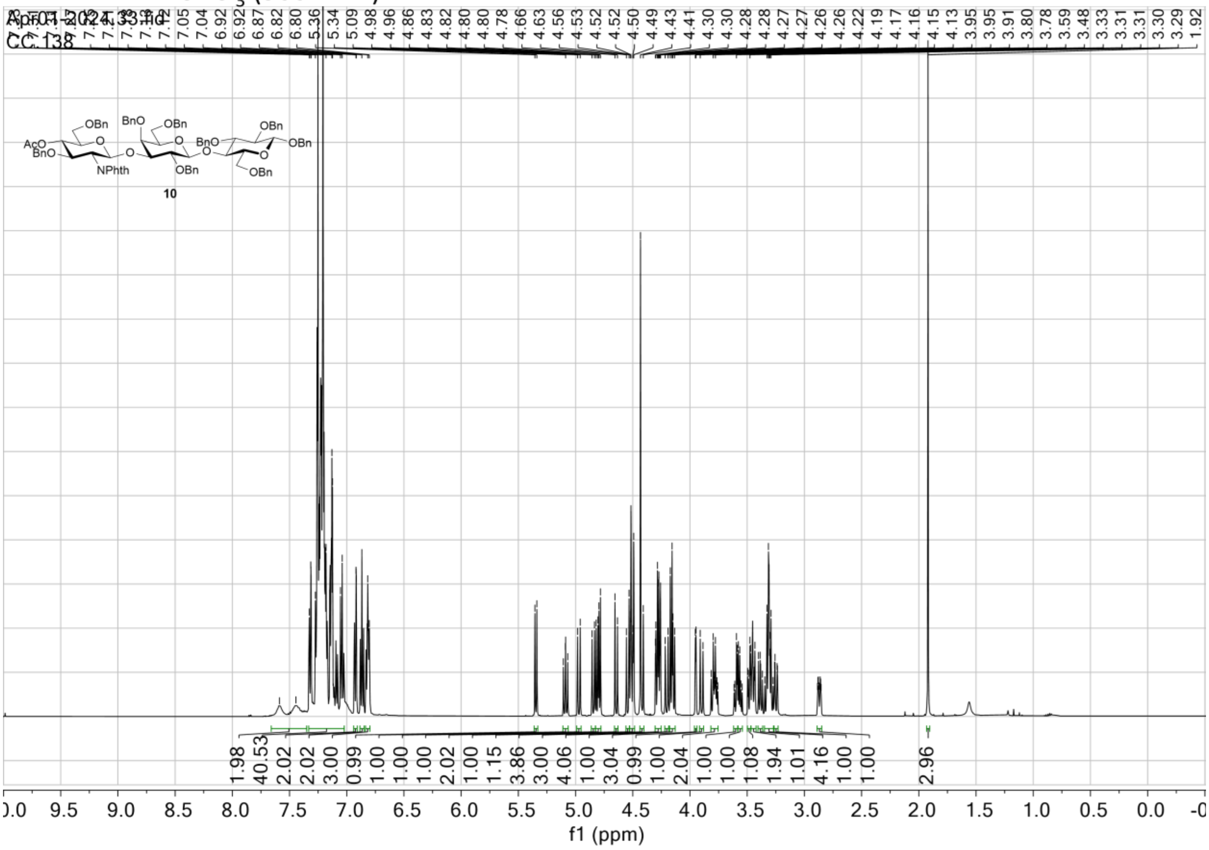

$^{13}C$ NMR (APT) in $CDCl_3$ (125 MHz)

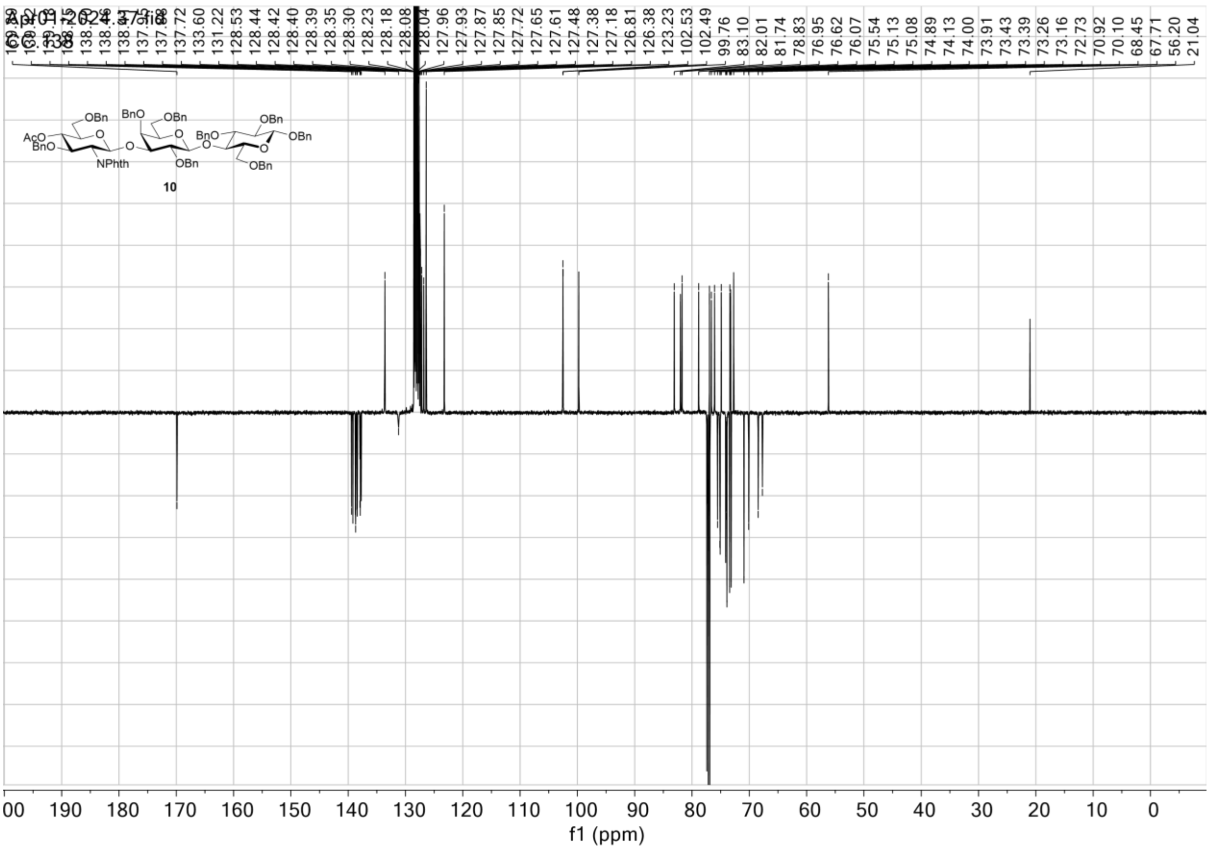

$^1H\_^1H$ gCOSY in $CDCl_3$

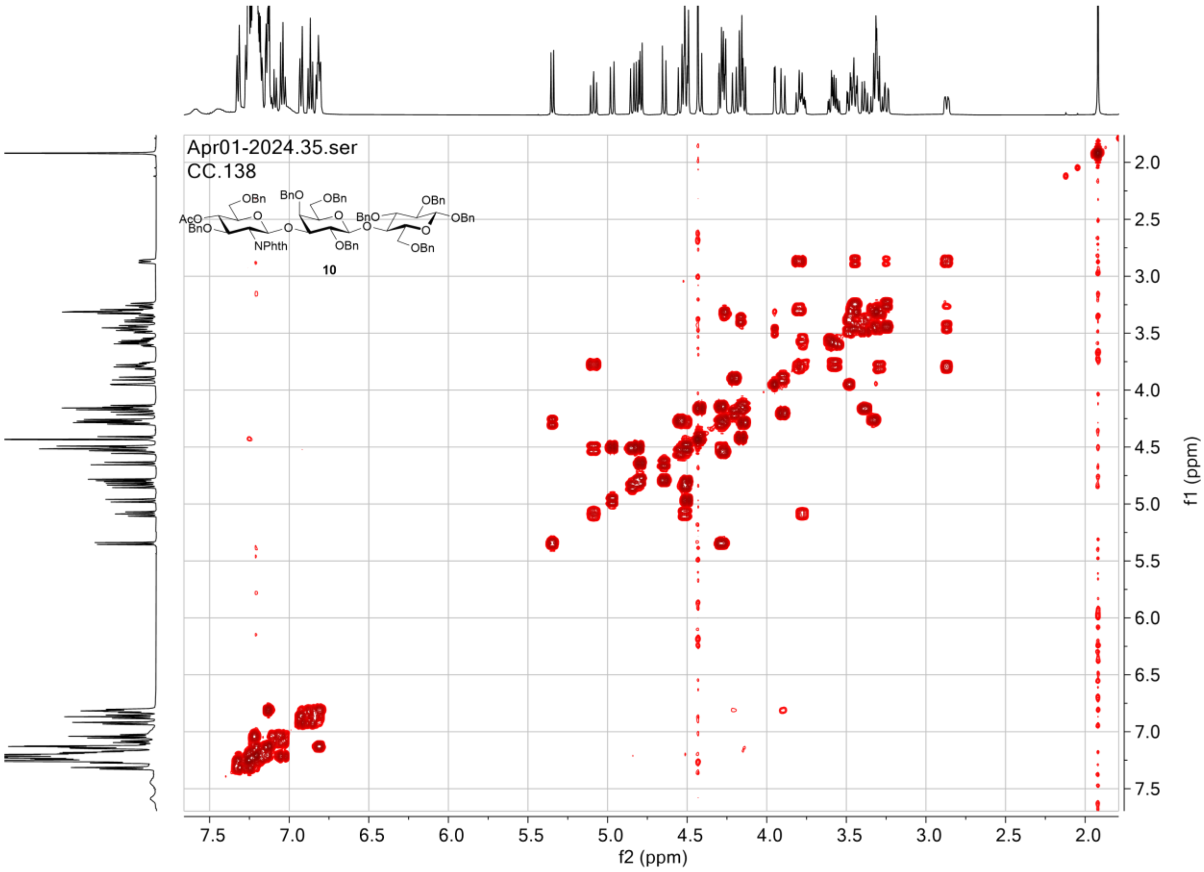

$^{1}H\_^{13}C$ gHSQC in $CDCl_3$

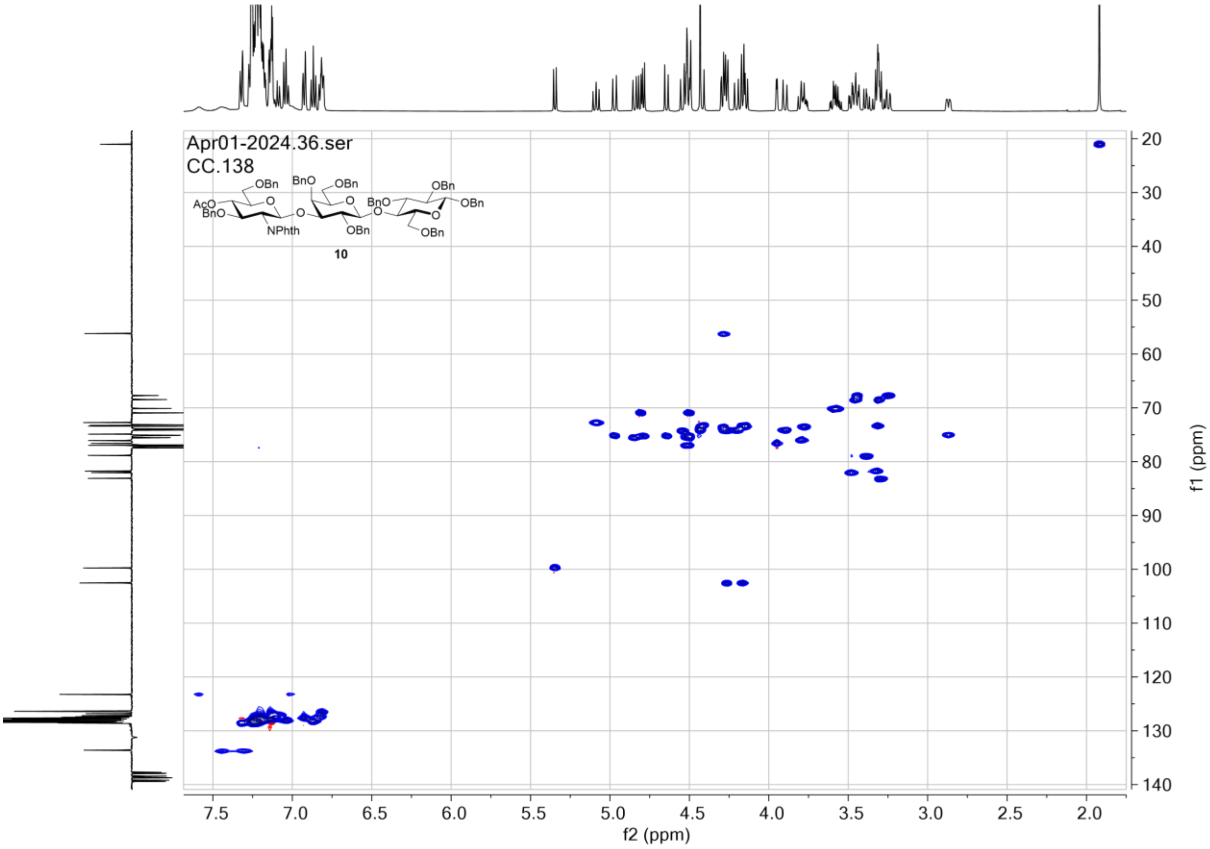

### NMR spectra for compound 11

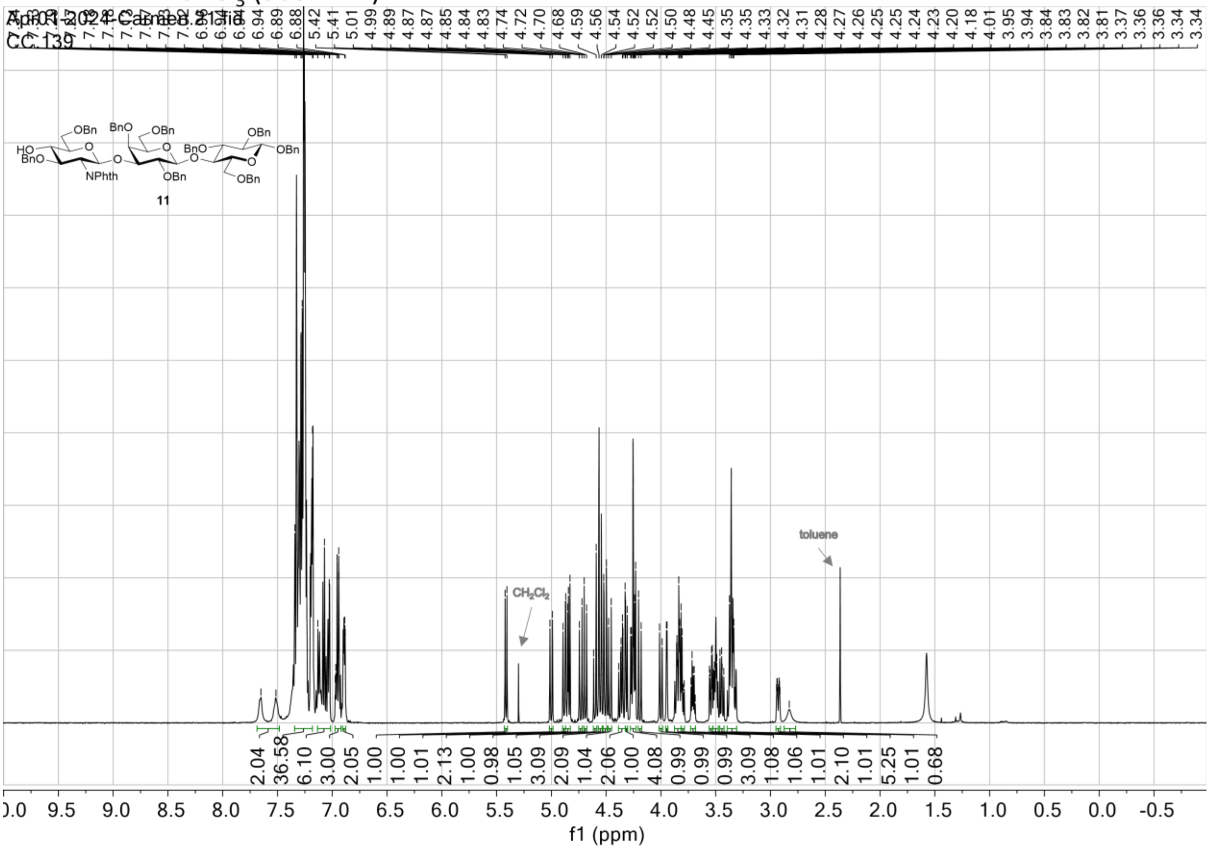

$^{13}C$ NMR (APT) in $CDCl_3$ (125 MHz)

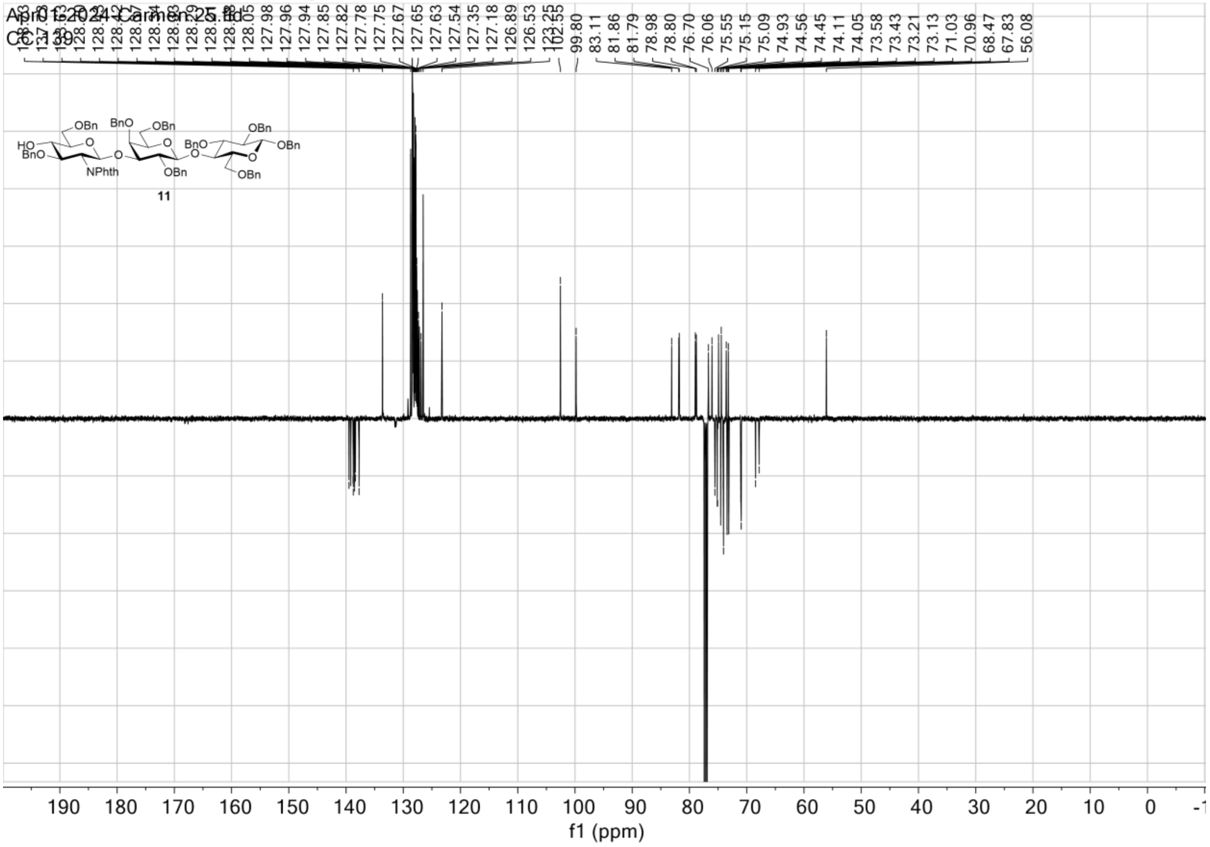

$^1H\_^1H$ gCOSY in $CDCl_3$

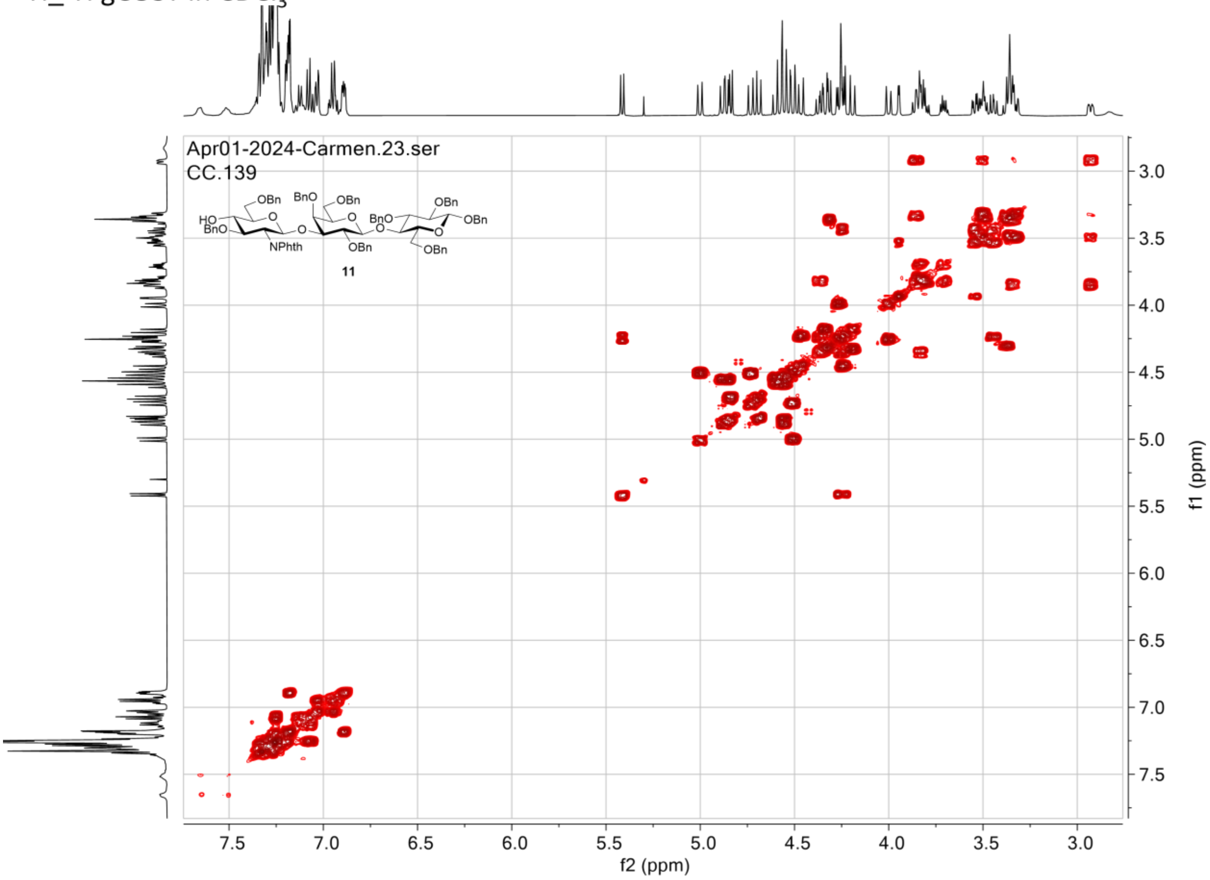

$^{1}H\_^{13}C$ gHSQC in $CDCl_3$

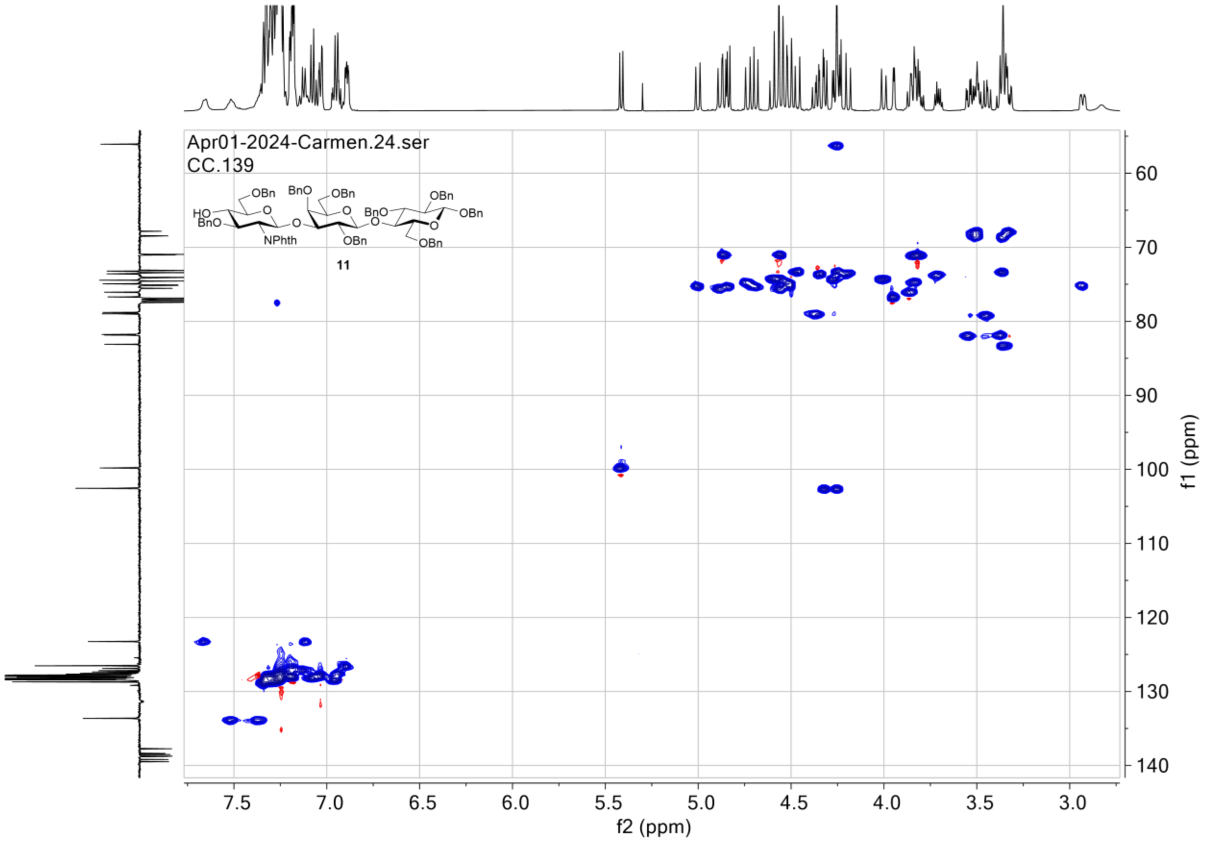

### NMR spectra for compound 12

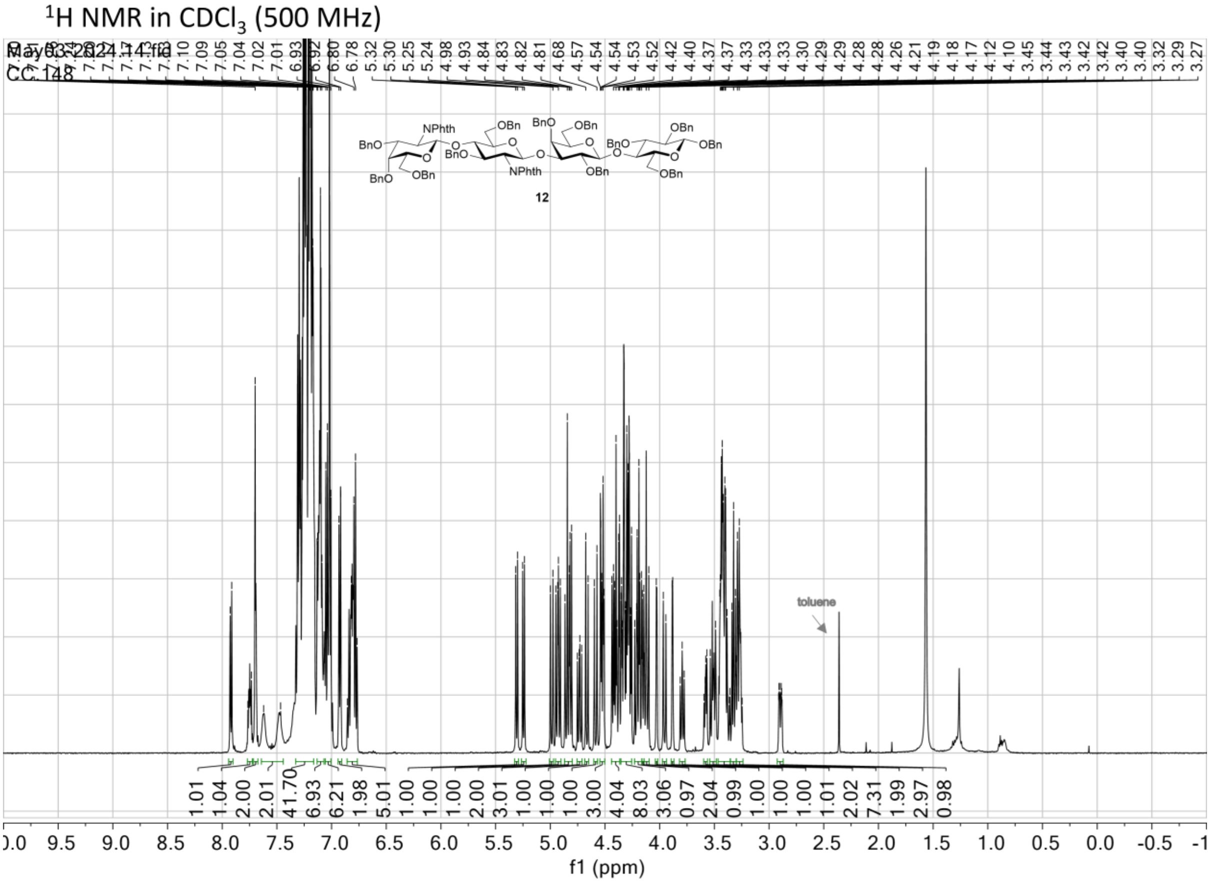

$^{13}C$ NMR (APT) in $CDCl_3$ (125 MHz)

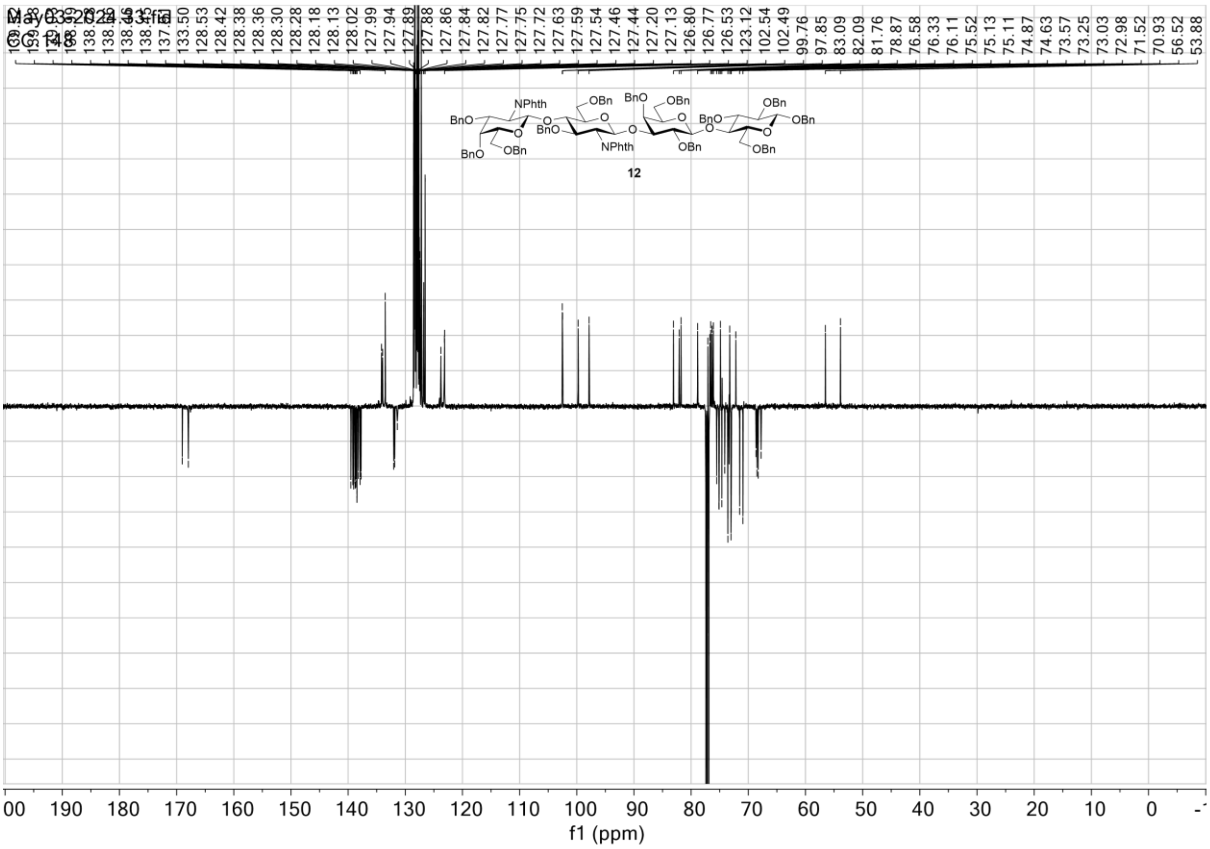

$^1H\_^1H$ gCOSY in $CDCl_3$

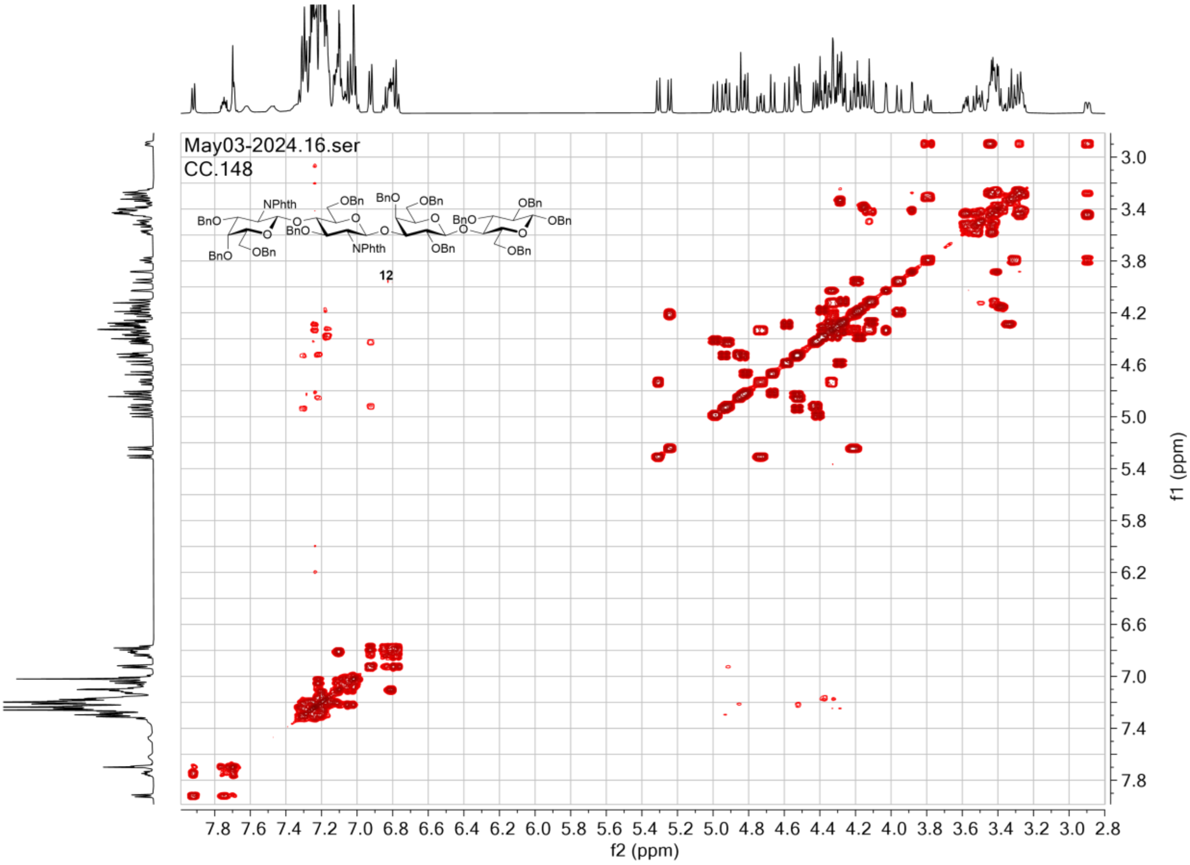

$^{1}H\_^{13}C$ gHSQC in $CDCl_3$

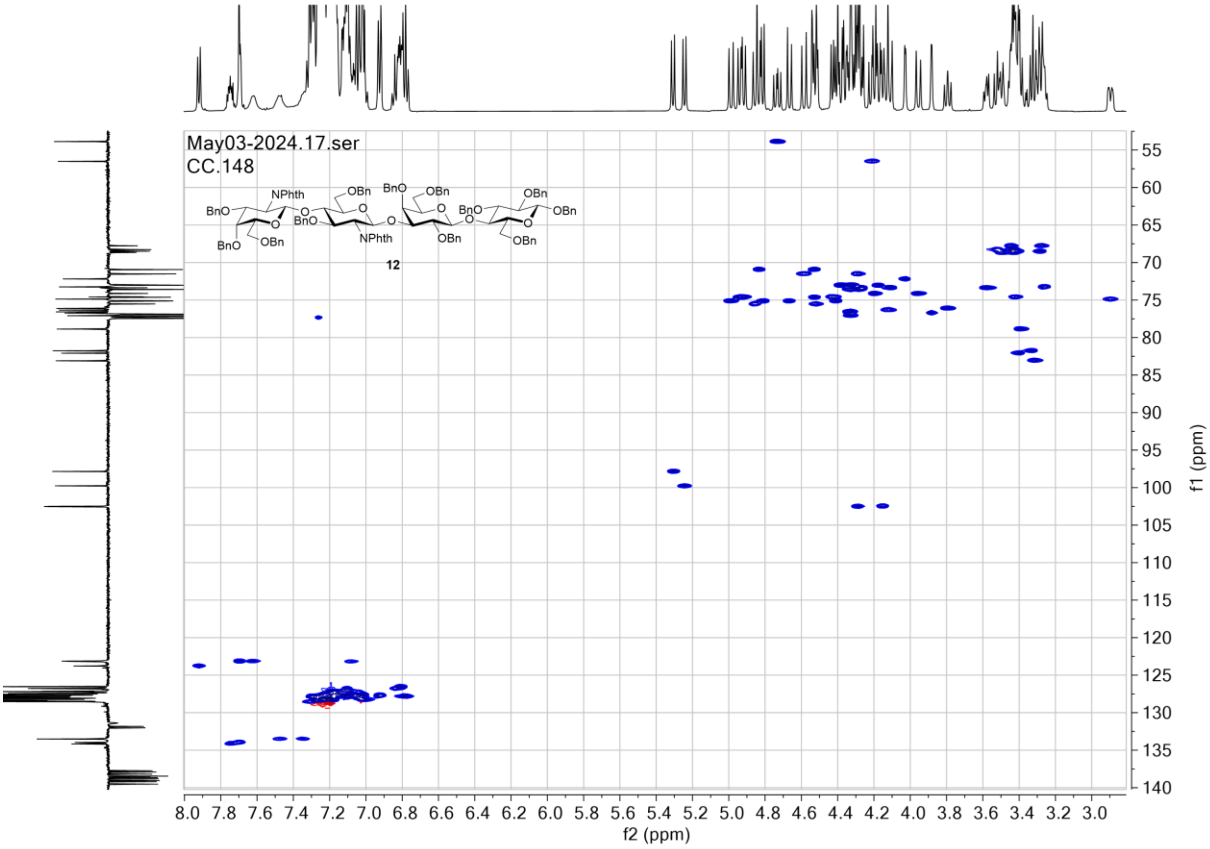

### NMR spectra for compound 13

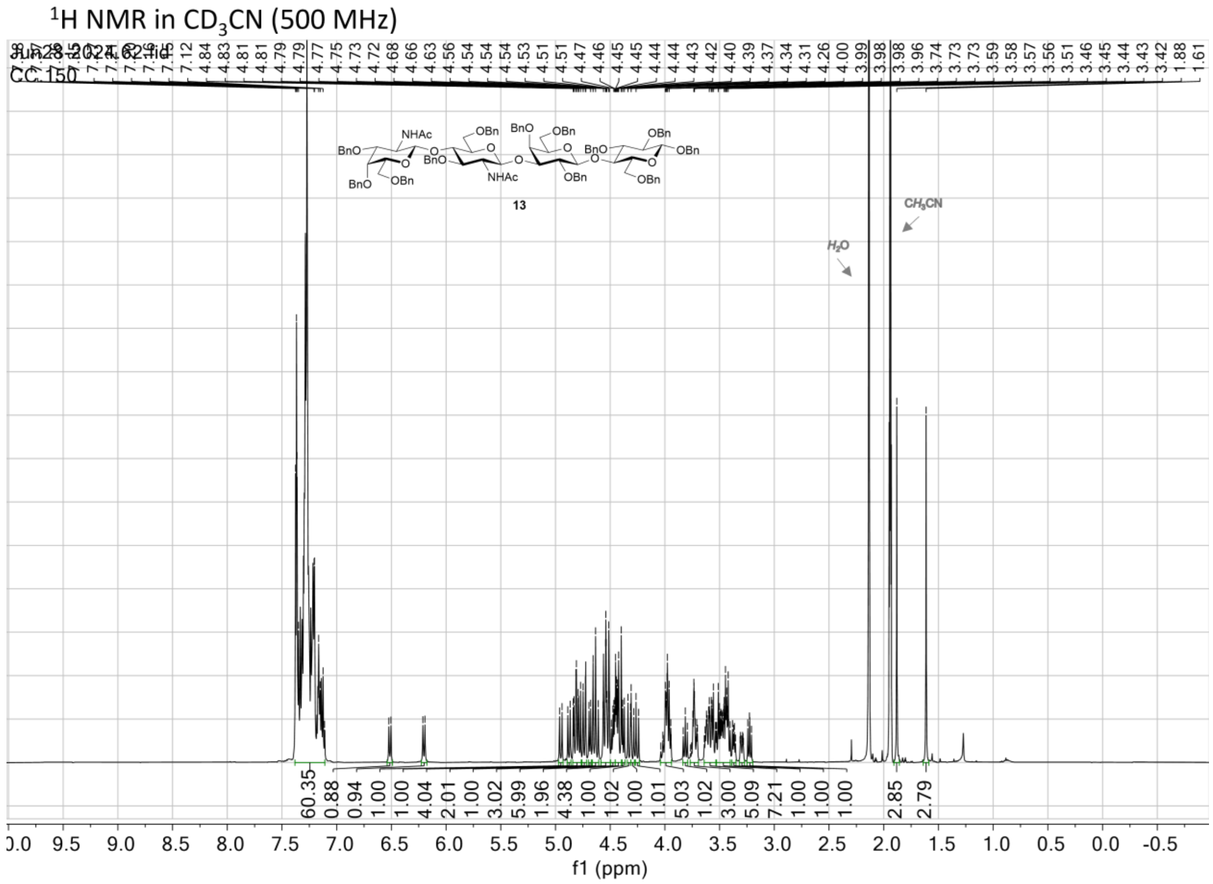

$^{1}H\_^{1}H$ gCOSY in $CD_3CN$

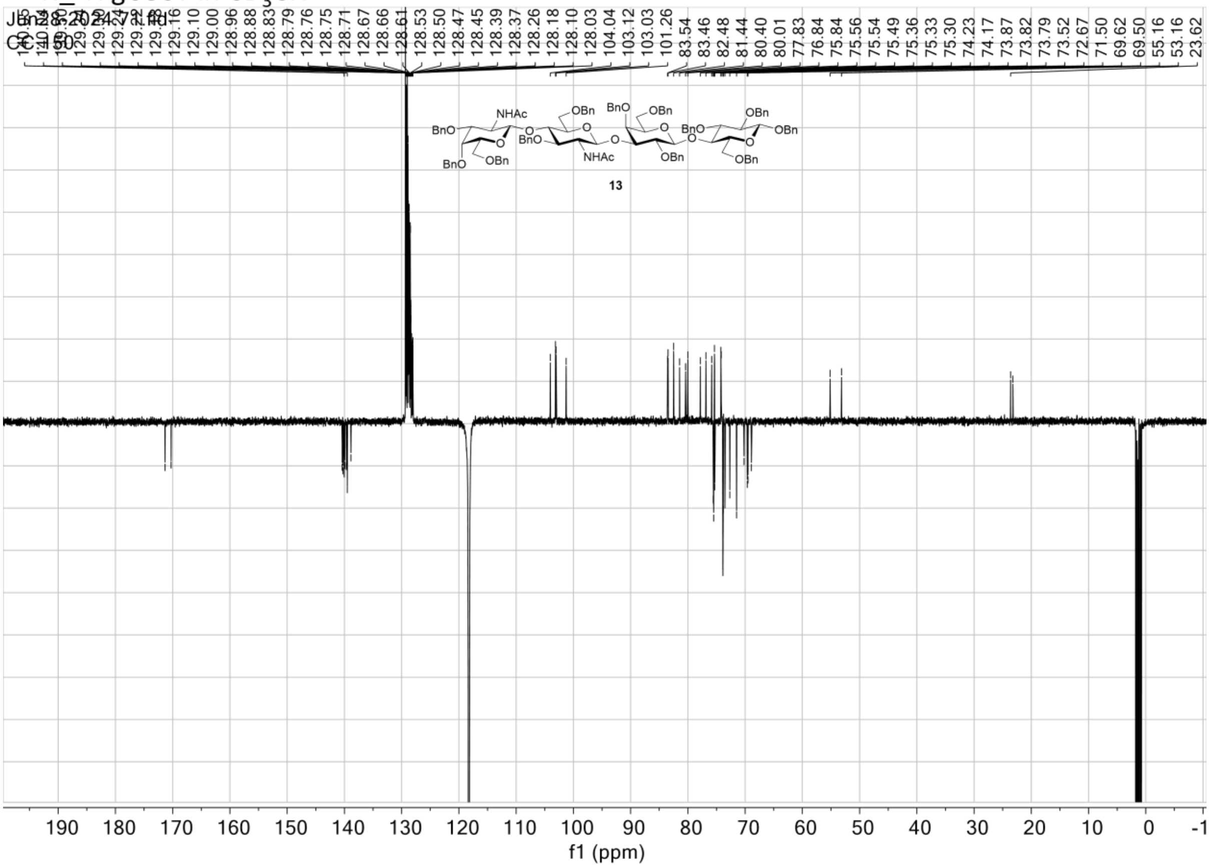

$^{1}H\_^{13}C$ gHSQC in $CD_3CN$

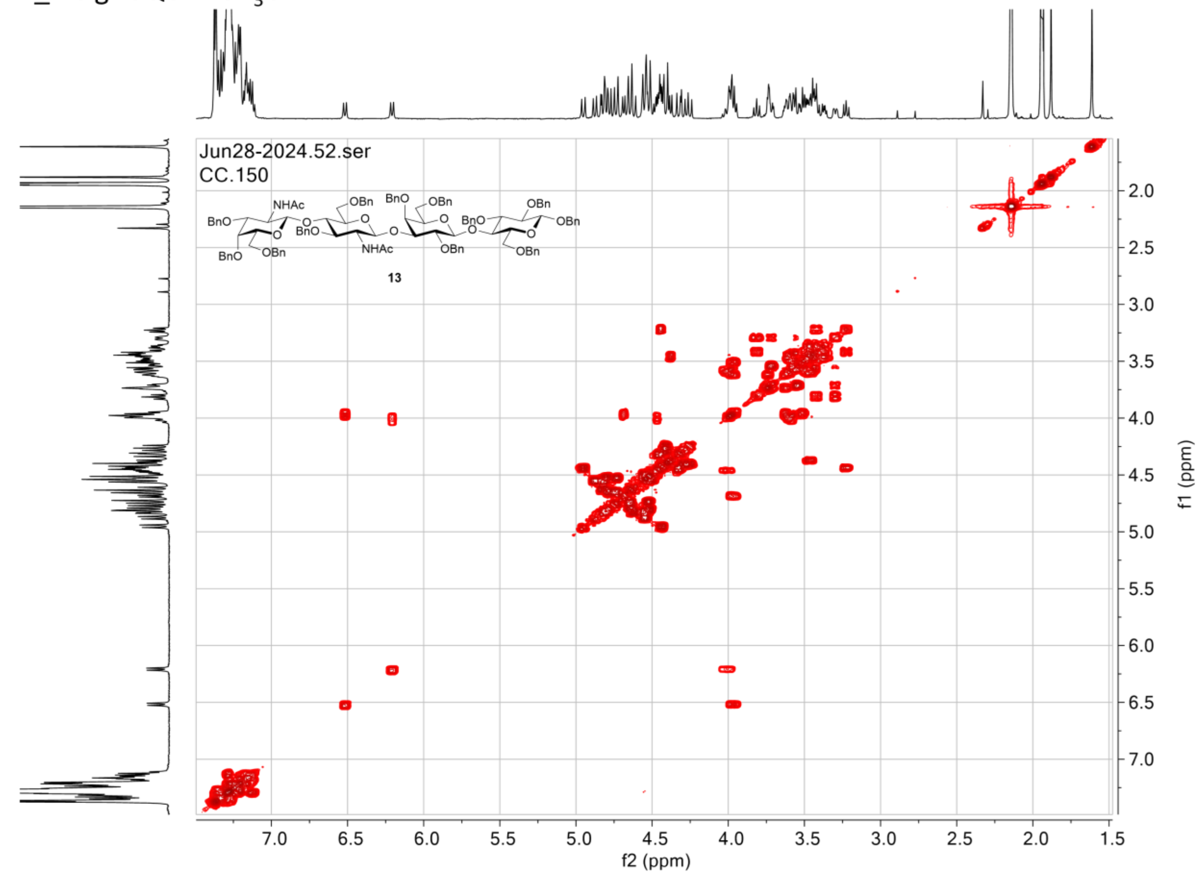

$^1H\_^{13}C$ gHSQC in $CD_3CN$

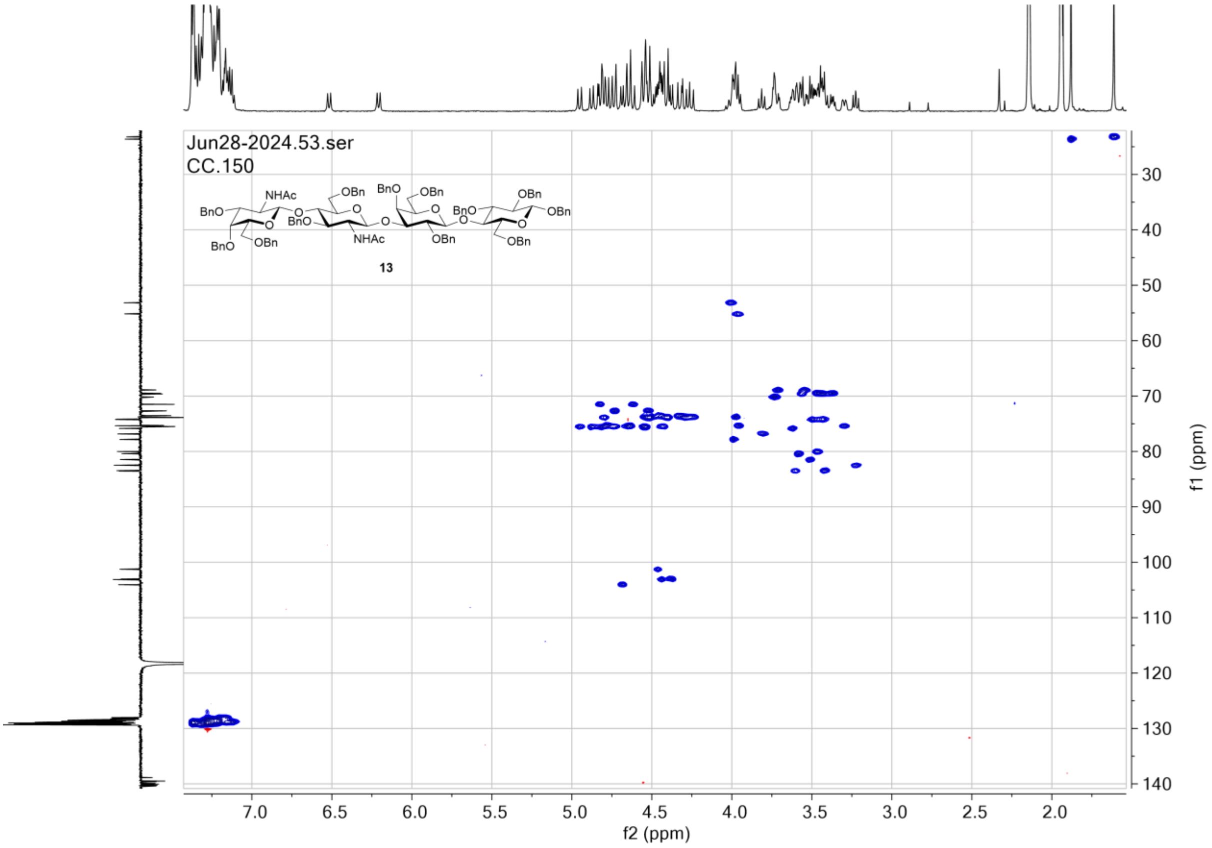

### NMR spectra for compound LdiNnT

[1]H NMR in $D_2O$ (500 MHz)

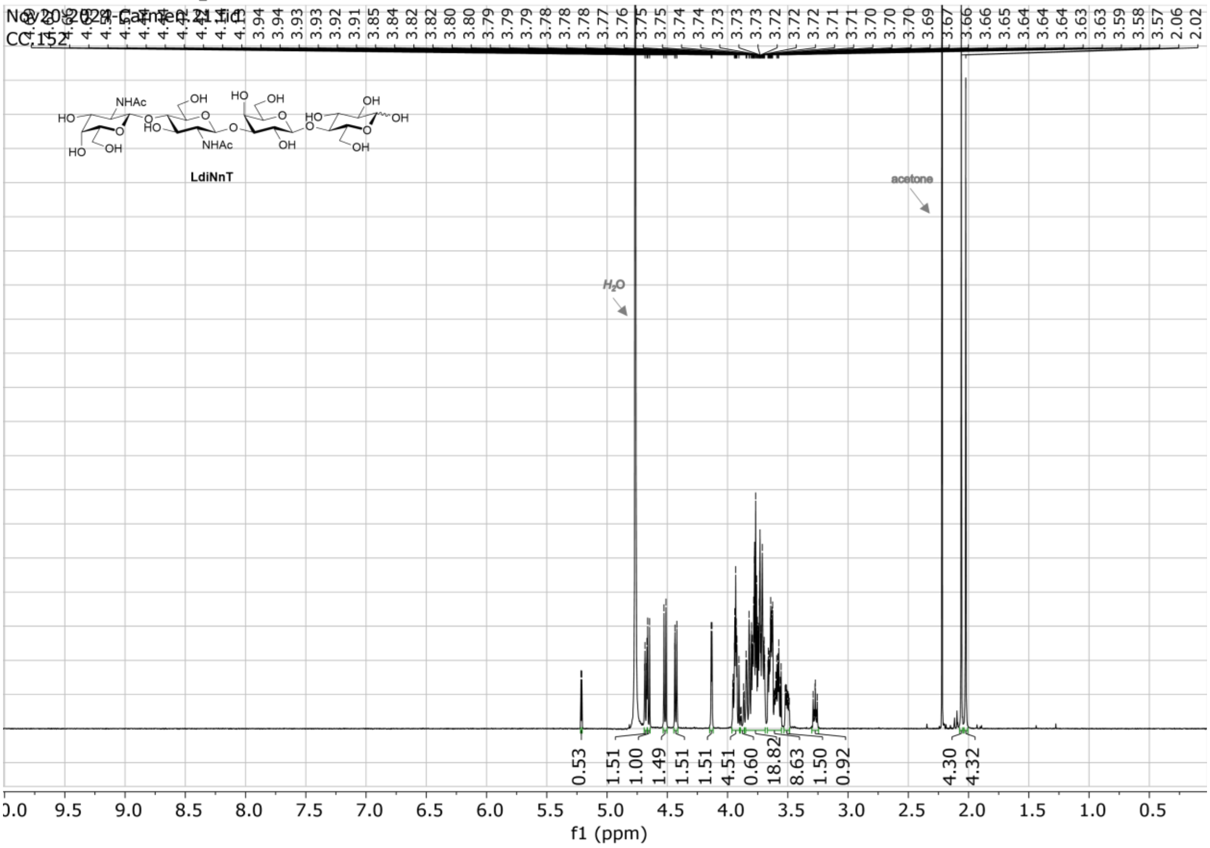

$^{13}$C NMR (APT) in $D_2O$ (125 MHz)

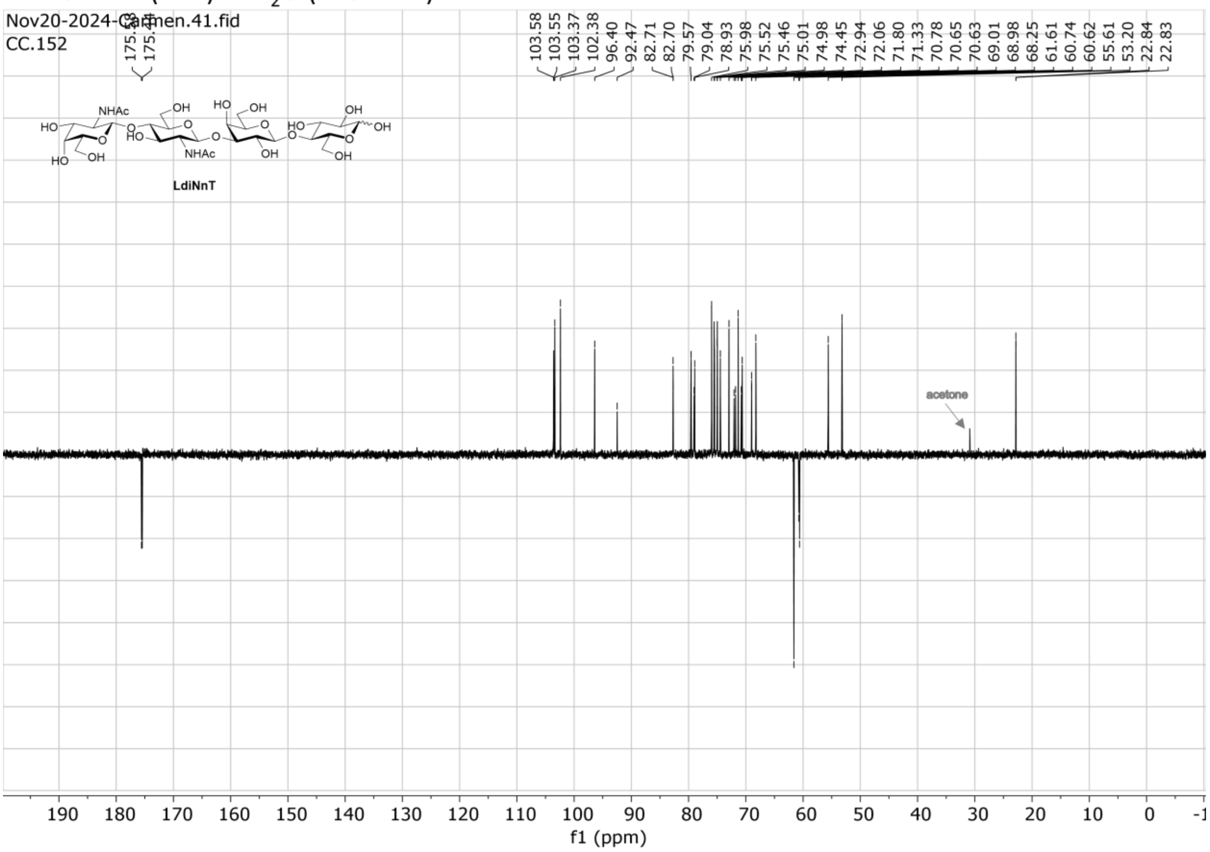

$^1$H_$^1$H gCOSY in $D_2O$

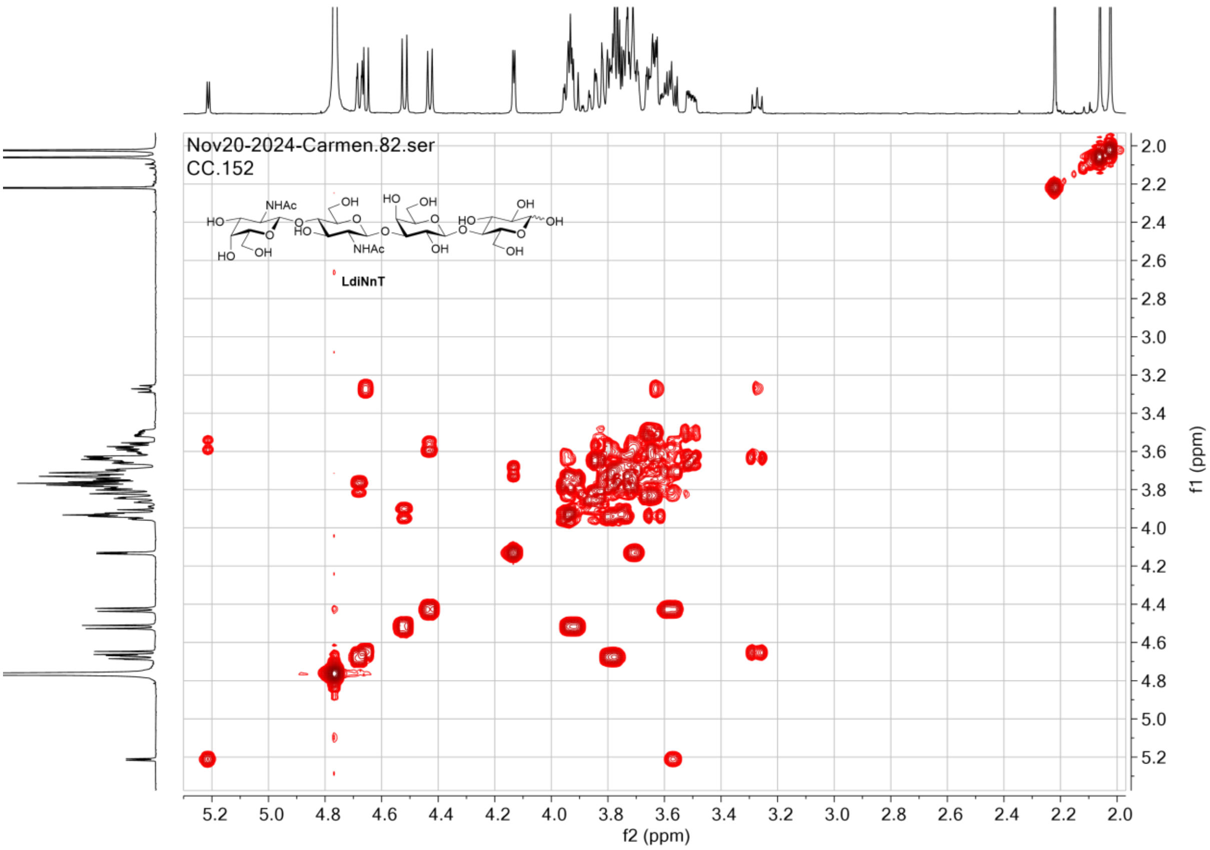

$^{1}H\_^{13}C$ gHSQC in $D_2O$

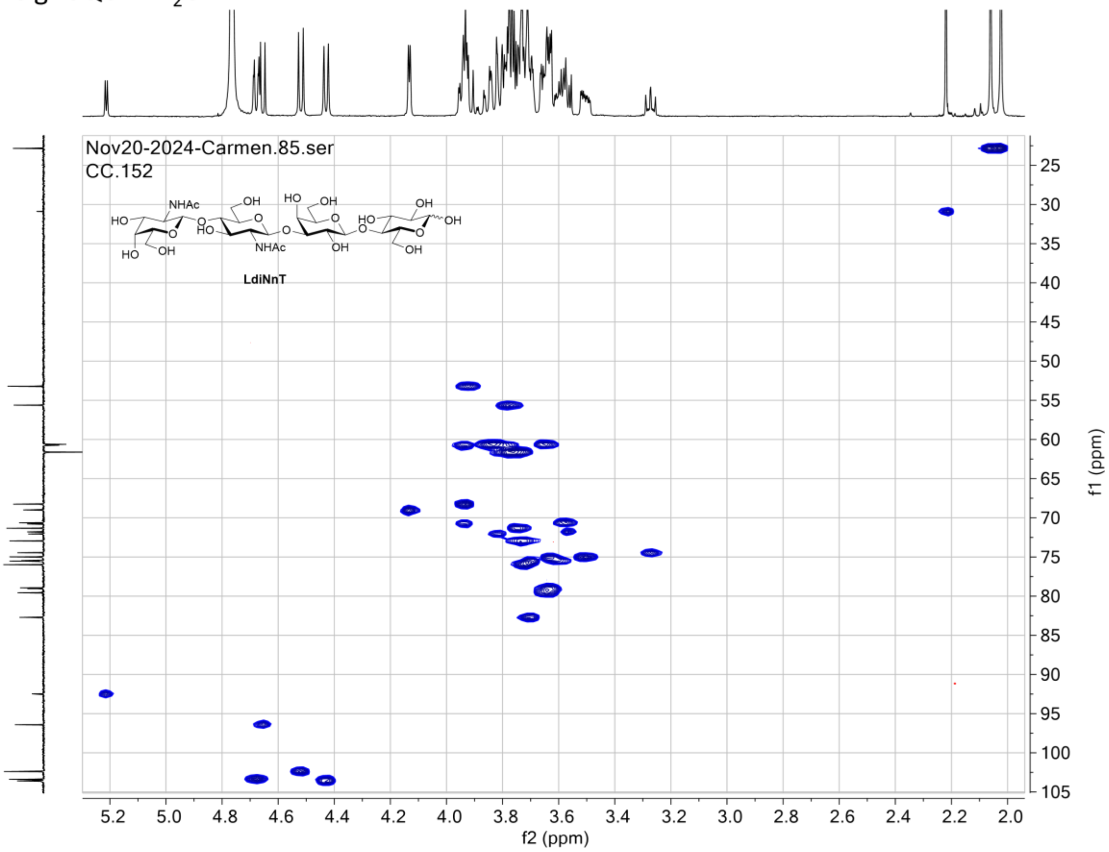

### HPLC purity analysis of LdiNnT

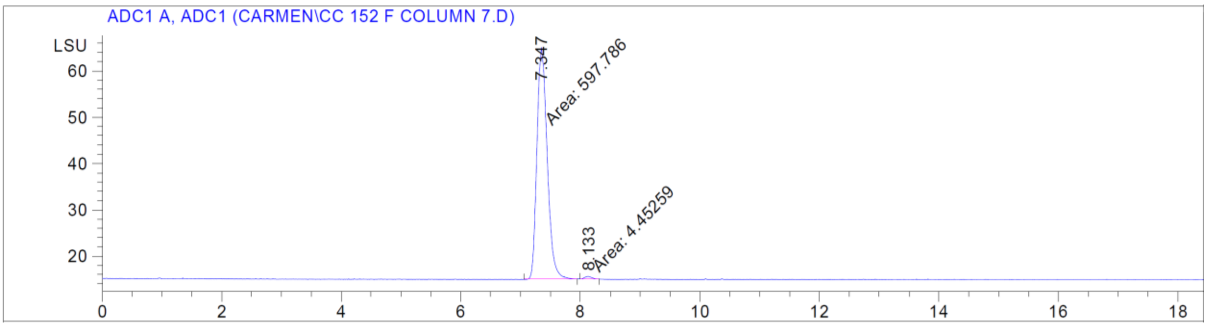
